# Mixed positive and negative feedback loops drive diverse single-cell gene expression dynamics

**DOI:** 10.1101/2025.01.14.632931

**Authors:** Torkel E. Loman, Christian P. Schwall, Teresa Saez, Yujia Liu, James C. W. Locke

## Abstract

Genetic circuits with only a few components can generate complex gene regulatory dynamics. Here, we combine stochastic modelling and single-cell time-lapse microscopy to reveal the possible dynamics generated by a key gene circuit motif: the mixed positive/negative feedback loop. Our minimal stochastic model of this motif reveals ten distinct classes of dynamic output, including stochastic pulsing, oscillations, and bistability. We systematically map how the circuit’s core parameters can be tuned to generate each of the behaviours. Experimental validation in two different mixed feedback circuits in the bacterium *Bacillus subtilis*, σ^B^ and σ^V^, confirms our model’s predictive power. Guided by our simulations, we are able to transition between dynamic behaviours by modulating in vivo parameters. Together, these results demonstrate how mixed feedback loops generate diverse single-cell dynamics, improving our understanding of this common biological network motif and informing our efforts to engineer them for synthetic biology applications.

## Introduction

Single-cell approaches have revealed that the outputs of gene circuits can be much more heterogeneous and dynamic than previously thought. Even small circuits with a few components can generate a wide range of regulatory dynamics, including bistability (1,2), oscillations (3,4), excitability (5,6), and pulsatile activation (7,8). Stochastic effects due to low molecule numbers often play a key role in generating these, sometimes beneficial, dynamics (9,10). In this study, we systematically explore the dynamic behaviours produced by a key circuit motif, the mixed positive/negative feedback loop, in which a single component both activates and represses its own expression.

Mixed positive/negative feedback loops are a widespread regulatory motif found in processes such as circadian rhythms (11), neural signalling (12), bacterial competence (13), the cell cycle (14), and DNA damage control (15). Mixed feedback architectures have also been proposed as useful regulatory motifs for synthetic biology applications (16). Despite their broad relevance, we lack a general understanding of the range of behaviours these circuits can generate, and how these are influenced by circuit properties.

To address this, we develop a minimal stochastic model of a mixed positive/negative feedback loop (Figure 1A). Stochastic modelling is essential for capturing intrinsic noise from low molecule numbers and extrinsic fluctuations from cellular processes like growth or energy availability (17–19). While prior models have explained behaviours such as excitability in bacterial competence (13,20) or circadian oscillations (21), these efforts typically model circuit-specific topologies and lack general predictive scope. By instead directly modelling effective properties of a mixed feedback circuit, like feedback strength, time delay, noise, and ultrasensitivity, we can predict what behaviours these motifs can generate.

**Figure 1:**
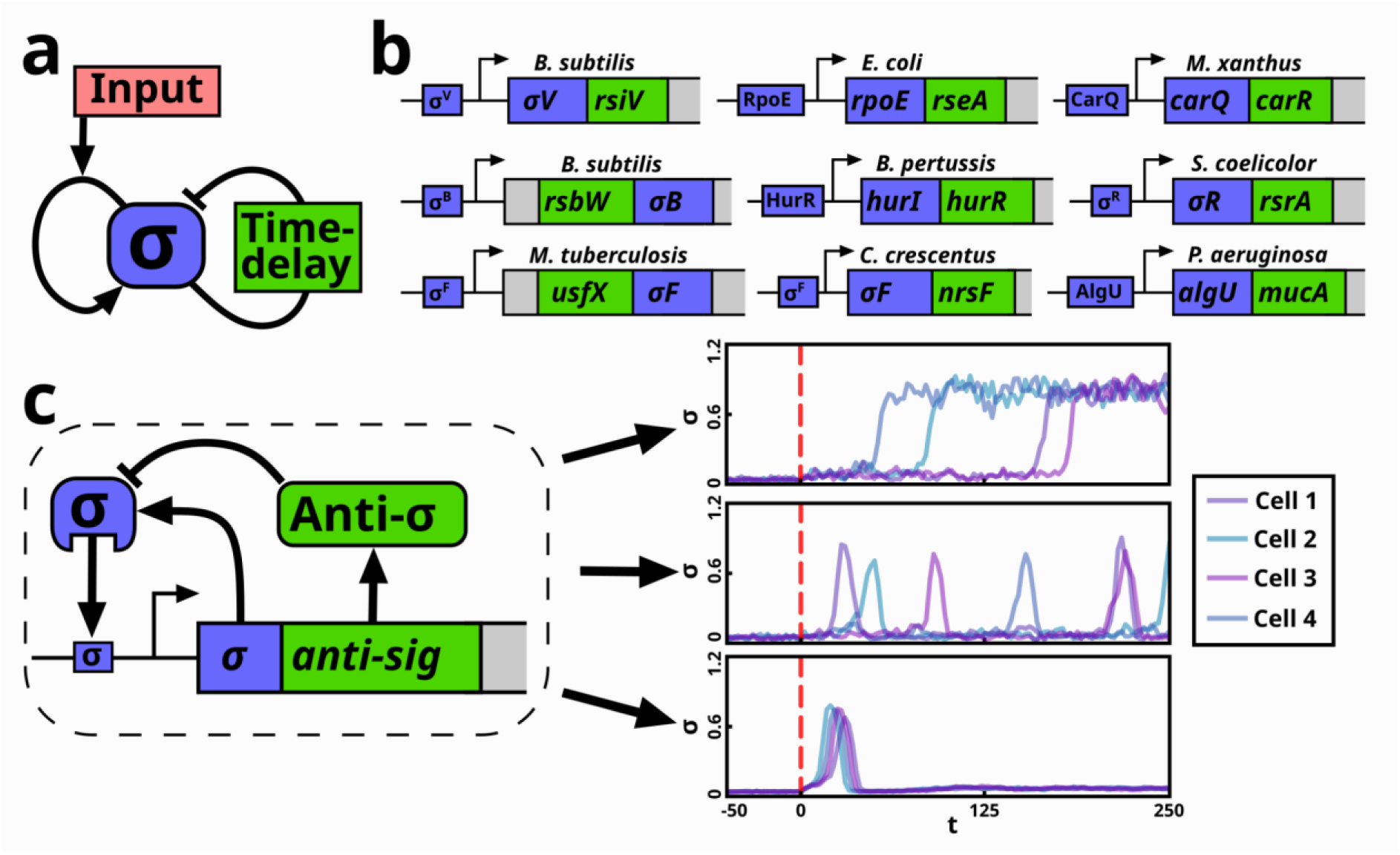
A general model of a mixed positive/negative feedback loop can reproduce experimentally observed sigma factor dynamics. **(a)** Schematic of the general mixed positive/negative feedback loop model. Here, a single component (σ) both activates and deactivates its own production. The activation depends on the presence of an input, and the deactivation is subject to a time delay. **(b)** The mixed positive/negative feedback loop is a common motif among alternative sigma factor circuits. Shown are simplified operon structures of alternative sigma factor circuits in a range of bacterial species, all consisting of a mixed positive/negative feedback. Anti-sigma factors (that repress sigma factor activity) are depicted in green, and sigma factors are depicted in blue. **(c)** Model simulations can capture known activation behaviours, including heterogeneous activation (top, as observed in e.g. σ^V^ (30)), stochastic pulsing (middle, as observed in e.g. σ^M^ (8)), and single pulse dynamics (bottom, as observed in σ^B^ under environmental stress (33)).

To validate our model predictions, we turn to bacterial alternative sigma factor circuits. Alternative sigma factors direct RNA polymerase to specific downstream targets and control key processes such as stress response, pathogenicity, and chemotaxis (22). The number of alternative sigma factors varies between bacterial species, ranging from 0 (as found in e.g. *Mycoplasma pneumoniae*), to over 63 (as found in *Streptomyces coelicolor*) (23–25).

Crucially, many alternative sigma factor circuits share a conserved mixed feedback motif: the sigma factor activates an operon that encodes both itself and an anti-sigma factor that inhibits its function (Figure 1b) (26–28). This simple, well-characterised, structure makes sigma factor circuits ideal systems for dissecting how mixed feedback architecture shapes dynamic behaviour.

Single-cell time-lapse fluorescence microscopy has revealed diverse, and often stochastic, gene expression dynamics across alternative sigma factor circuits. Using microfluidic devices such as the mother machine (29), studies have uncovered behaviours including heterogeneous activation (σ^V^ in *B. subtilis* (*30*)), stochastic pulsing (σ^M^, σ^W^, σ^X^, σ^D^, σ^B^ in *B. subtilis* and RpoS in *E. coli* (*8,31,32*)), and a single response pulse (σ^B^ in *B. subtilis* (*33*)). Here we use σ^V^ as a model system for heterogeneous activation and σ^B^ as a model system for pulsatile dynamics (See Supplementary Figures 1, 2 for full network details).

Both systems contain mixed feedback architectures. σ^V^ is activated by lysozyme stress via cleavage of its anti-sigma factor RsiV (34–40), and in turn activates its own operon containing both *sigV* and *rsiV* (Supplementary Figure 1). σ^B^ is regulated through a partner-switching mechanism involving its anti-sigma factor RsbW and the anti–anti-sigma RsbV (41–45), forming a more complex feedback structure (Supplementary Figure 2). Despite their biological differences, both systems share a core regulatory motif in which a sigma factor promotes expression of both itself and its anti-sigma factor.

In this work, we develop a general stochastic model of a mixed positive/negative feedback loop. The model predicts ten distinct dynamic behaviours, some of which have been previously observed and others newly predicted, and maps how they emerge from variations in a few biologically meaningful parameters. Using genetically rewired σ^V^ and σ^B^ circuits in *Bacillus subtilis* (Supplementary Figures 3, 4), we experimentally validate the predicted transitions between behaviours. Our work develops general principles of how these behaviours depend on circuit properties, demonstrating how native or engineered systems can be tuned to generate target behaviours.

## Results

### A general mixed positive/negative feedback loop model can reproduce experimentally observed sigma factor dynamics

To explore the range of dynamics that mixed feedback loop genetic circuits can generate, we developed a minimal stochastic model (Methods, Figure 1a). In this work, we apply it to sigma factor networks, biologically relevant systems that frequently implement mixed feedback loops (Figure 1b). The model is implemented as a stochastic differential equation (SDE) tracking the concentration of a single component (*σ*), which both activates and represses its own production through a Hill-type regulatory function. The self-repression is subject to a time delay (*τ*). In the context of sigma factors, this delay can be motivated either by the anti-sigma factor undergoing additional steps before it can interact with the sigma factor (46), or due to the presence of additional feedback mechanisms (47).

The initial formulation of our model depended on 9 parameters. Because our aim was to understand general qualitative behaviours rather than fit specific datasets, we performed nondimensionalisation, reducing the model to six biologically interpretable parameters (Supplementary Notes 2, Supplementary Tables 1, 2). These are: the system’s degree of self-activation (*S*), self-deactivation (*D*), the time delay in the self-deactivation (*τ*), base operon activity (*v_0_*), ultrasensitivity (*n*), and noise amplitude (*η*). The deterministic version of the model, which excludes the noise parameter *η*, is given by:

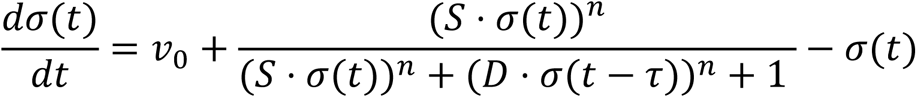

We implemented noise using the chemical Langevin equation (CLE) (48), a well-established method for modelling noise within gene regulatory networks (49). By representing the system as a stochastic differential equation (SDE), the CLE enables fast stochastic simulations when compared to the more exact Gillespie algorithm (50,51) (Supplementary Notes 3). Our results were not specific to our choice of stochastic interpretation, as we verified that simulating noise through the Gillespie algorithm generated equivalent results (Supplementary Notes 3, Supplementary Figures 5, 6).

We implemented the delay, *τ*, as a distributed delay using the linear chain trick (Supplementary Notes 4.1, Supplementary Figures 7, 8) (52,53). This enabled us to model the system as an SDE, rather than as a Stochastic Delay Differential Equation (SDDE), which is advantageous for model simulation and analysis (Methods). For completeness, we also implemented an SDDE version of the model and confirmed it reproduces key results (Supplementary Notes 4.2, Supplementary Figure 9, Supplementary Table 3). Finally, we note that our minimal mixed feedback loop model captures the range of sigma factor response behaviours reported in the literature, including heterogeneous activation, stochastic pulsing, and single response pulse (Figure 1C) (8,30,33).

### The model generates ten distinct response behaviours

To explore the range of behaviours the model can exhibit, and to understand how these depend on its parameters, we developed a behaviour classification scheme. The scheme is based on steady state analysis, where we note that the system can have either a fixed point in an inactive state, an active state, both, or a single fixed point of intermediate activity (Supplementary Notes 5, Supplementary Figures 10, 11). Asymptotically, it will approach one of these states. However, perturbations, either due to noise or onset of the input, can cause a transition to another state. By enumerating all possible combinations of these transitions, we find a total of ten distinct behavioural classes (Figure 2, Supplementary Notes 6, Supplementary Figures 12, 13, 14). By simulating the model across a large set of randomised parameter values and checking that each simulation aligned with one of these ten classes, we confirmed that these classes are robust. (Supplementary Figures 15, 16). Rare additional behaviours that can be considered subclasses of these 10 are discussed in Supplementary Notes 6.4.

**Figure 2.**
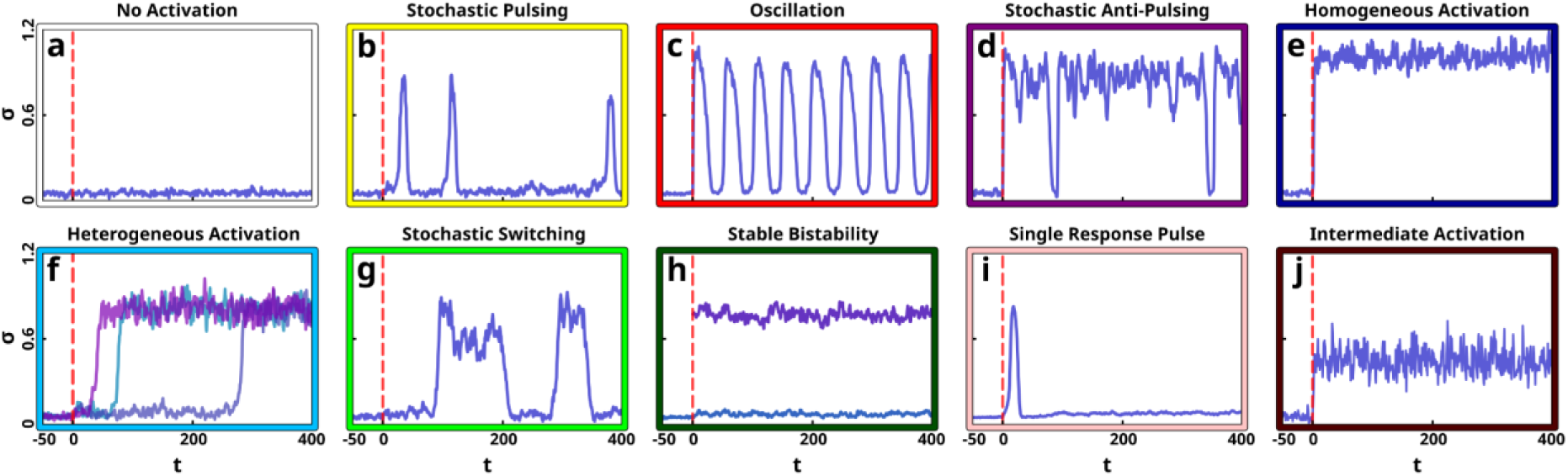
The model generates 10 distinct response behaviours. Simulations for 10 different parameter sets, demonstrating the 10 different behaviours the model can achieve. Initially, the system remains inactive; upon triggering the input (marked by the red dashed lines), distinct behaviours emerge: **(a)** No activation. **(b)** Stochastic pulsing. **(c)** Oscillation. **(d)** Stochastic anti-pulsing. **(e)** Homogeneous activation. **(f)** Heterogeneous activation (three different simulations displayed). **(g)** Stochastic switching. **(h)** Stable bistability. The figure depicts two simulations. The first (dark blue) is initiated at the initial (inactive) state at the same time as the simulations in the other figures. The second (purple) is initiated at the time of the input (dashed red line) in the active state. There is no intrinsic way for the system to switch between these two states (however, it could happen by e.g. the addition of exogenous σ). **(i)** Single response pulse. **(j)** Intermediate activation.

Through our classification scheme, we identified ten distinct response behaviours predicted by the model (Figure 2):

1. No activity: The system remains inactive. In the absence of the input, this is also the system’s default behaviour (Figure 2a, Supplementary Figure 17).
2. Stochastic pulsing: The system exhibits randomly distributed pulses of activity in time, from an otherwise inactive state (Figure 2b, Supplementary Figure 18).
3. Oscillation: The system alternates between low and high activity in a periodic manner (Figure 2c, Supplementary Figure 19).
4. Stochastic anti-pulsing: The system activates in response to the input, but displays randomly distributed pulses of inactivity over time (Figure 2d, Supplementary Figure 20).
5. Homogeneous activation: In response to the input, the system transitions from an inactive to an active state. The response times are instantaneous (and thus homogeneous across simulations) (Figure 2e, Supplementary Figure 21).
6. Heterogeneous activation: In response to the input, the system transitions from an inactive to an active state. The response times are random (and thus heterogeneous across simulations) (Figure 2f, Supplementary Figure 22).
7. Stochastic switching: The system randomly alternates between an inactive and an active state (Figure 2g, Supplementary Figure 23).
8. Stable bistability: The system has an inactive and an active state, but cannot switch between them intrinsically. An extrinsic way to induce such a switch would be to add additional σ from an external source (Figure 2h, Supplementary Figure 24).
9. Single response pulse: The system exhibits a single pulse of activity at the input’s onset, but no significant activity thereafter (Figure 2i, Supplementary Figure 25).
10. Intermediate activation: Following input onset, the system transitions to and remains in a state of intermediate activity (Figure 2j, Supplementary Figure 26). Here, while the activation times are typically homogeneous, certain low-*D* valued regions display heterogeneous intermediate activation (Supplementary Notes 6.4).

These behaviours include all previously observed dynamics among bacterial sigma factors (a single response pulse, stochastic pulsing, heterogeneous activation, and homogeneous activation) (8,30,31,33), as well as several novel behaviours. Detailed descriptions of each behaviour are provided in the supplementary material (Supplementary Notes 7).

### Systematic mapping of behaviours reveals their dependency on model parameters

Next, we investigated how the system’s behaviour depends on its six parameters, (*S*, *D*, *τ*, *v_0_*, *n*, and *η*). To do this, we developed a heuristic classifier that takes a parameter set as input and assigns it to one of the 10 behavioural classes (Supplementary Notes 8, Supplementary Figure 12). The classifier uses a decision tree in which each node performs simulations initialised at the model’s steady states and measures the transition times to other steady states (Supplementary Figure 14). We confirmed the accuracy of the classifier by empirically validating its classifications across a large random sample (Supplementary Figures 15, 16). Applying the classifier across the 6d parameter space allowed us to construct a map showing which behaviour occurs for each parameter combination (Supplementary Notes 8). To visualise how individual system properties influence resulting dynamic behaviours, we plotted 2d slices of this parameter space while holding the remaining four parameters constant (Figure 3).

**Figure 3.**
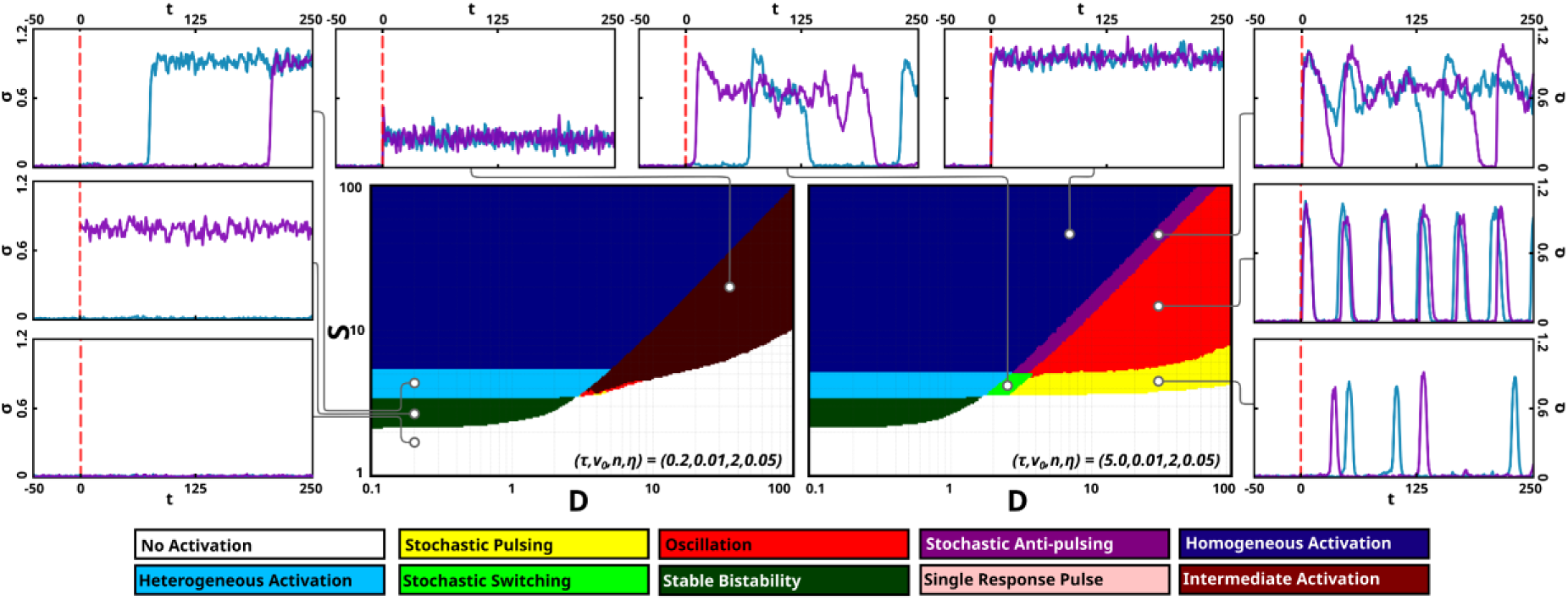
Mapping system behaviours across parameter space reveals distinct dynamic regimes. We applied our behaviour classifier to each point in 6d parameter space of the model and visualised 2d slices in log-scale S-D space, with the remaining parameters, v_0_, n, and η held constant. The two maps shown here differ only in the value of the time delay τ: τ = 0.2 (left) and τ = 5.0 (right). Each point is coloured by the classified dynamic behaviour, and example trajectories from selected points illustrate representative single-cell simulations. While the exact boundaries shift with τ, both maps exhibit a common structure: a D ≲ 1 region with bistable and switching behaviours, and a D ≲ 1 region that supports either intermediate activation (left) or oscillatory and pulsing dynamics (right). In these two maps, the transition occurs at around D ≈ 2, and in general, the transition depends on v_0_ but is typically close to 1.This general division by D is robust across parameter choices and is supported by stability analysis (Supplementary Figure 27). The effects of varying τ, v₀, n, and η are further explored in Supplementary Figures 28-32 (which shows additional behaviour maps).

When examining parameter maps for various values of *τ*, *v_0_*, *n*, and *η*, we found that they can be broadly divided into two regions: *D≲1* and *D≲1* (Supplementary Figure 27). Each region is characterised by a distinct behavioural transition as the system’s self-activation strength (S) increases. We note that *S* is a composite parameter which, after non-dimensionalisation, also incorporates the system’s input strength. Hence, *S* can be tuned by either modifying the self-activation strength or by varying the input. In the *D≲1* region, the model exhibits behaviours associated with bistability, such as stochastic switching, stable bistability, and heterogeneous activation. Conversely, the *D≲1* region exhibits behaviours related to oscillations and excitability, including stochastic pulsing, oscillations, and stochastic anti-pulsing, although it can also exhibit intermediate activation under some conditions (Supplementary Notes 8, Supplementary Figures 28-32).

Next, we characterised how the behaviours depend on the parameters *τ*, *v₀*, *n*, and *η* by measuring their effect on the prevalence of each behaviour (Supplementary Figures 17-26). Generally, the time delay *τ* had the largest effect on system behaviour (after *S* and *D*), being especially influential in the *D* ≲ 1 region (Supplementary Table 4). Figure 3 illustrates this effect: two maps are shown in which all parameters are held constant except *τ*, which is 0.2 in the left panel and 5.0 in the right. At low τ, the D ≲ 1 region supports intermediate activation, while longer delays give rise to stochastic pulsing and oscillations. In contrast, *τ* has minimal impact in the D ≲ 1 region, where weak self-deactivation renders the delay less consequential. Conversely, *v_0_*(the base activity of the operon) had only minor effects on the D≲1 region, while all behaviours in the D≲1 grew less prevalent as *v_0_* grew larger (Supplementary Notes 7, Supplementary Figure 30, 31). Almost all behaviours were affected by *η* (the noise amplitude), while *n* (the degree of system ultrasensitivity) had the least pronounced effect on the system’s behaviour (Supplementary Notes 7, Supplementary Figure 29, 31, 32).

Our behaviour maps also permit us to directly observe how noise warps behaviour space. At low noise levels, behavioural transitions coincide well with the bifurcation points of the deterministic system. However, as *η* (the noise amplitude) is increased, the “effective” bifurcation points, such as those marking the emergence of oscillatory behaviours, increasingly shift in behaviour space (Supplementary Figure 33).

Finally, we note that σ^V^ and σ^B^ naturally occupy distinct regions of parameter space, with σ^V^ exhibiting heterogeneous activation in the *D≲1* regime and σ^B^ showing stochastic pulsing in the D≲1 regime. This separation allows us to explore different parts of the behavioural map by selectively tuning parameters in each system. In the following sections, we use σ^V^ to probe transitions across the *D* axis and σ^B^ to examine transitions driven by changes in *S*, experimentally validating our model’s predictions in both cases.

### The predicted ability to transition between D≲1 and D≲1 behavioural domains is validated in the σ^V^ system

We first experimentally tested the transitions predicted for the bistability-dominated *D≲1* region, using the *B. subtilis* alternative sigma factor σ^V^ as a model system. σ^V^ regulates the lysozyme stress response in *B. subtilis*, activating genes related to cell wall protection and repair (34,35,37,38,54). In a previous study, we documented the behavioural transition of the wild-type system as *S* is varied (30). In that study, lysozyme stress input was varied, which corresponds to a modulation of *S* in our simulations. Using single-cell time-lapse microscopy, the study demonstrated a transition from no activation, to heterogeneous activation, to more homogeneous activation, as lysozyme was increased, similar to the transitions observed when *S* is increased in the *D≲1* region of the parameter map (Figure 3). Although our model often (but not always) includes a stable bistable regime between the no-activation and heterogeneous-activation regions, this behaviour will appear as no activity in these simple input-onset experiments (Supplementary Tables 5, 6). With this consideration, the experimental observations align well with the model predictions.

Our model predicts that the *D≲1* and *D≲1* regions coexists within the same system, and that increasing *D* (the strength of self-deactivation) can shift the system from one regime to the other. To test this, we engineered a series of modified σ^V^ circuits in which *D* was increased by adding additional copies of the anti-sigma factor (*rsiV*) (Methods) (Supplementary Figure 3). As *rsiV* expression is upregulated by σ^V^ and RsiV inhibits σ^V^ activity (39,40), this manipulation allows us to tune the system’s self-deactivation strength.

According to our model, increasing *D* should progressively reduce σ^V^ activity in the ON state while preserving heterogeneous activation; at sufficiently high *D*, all activity should be suppressed (Figure 4a–b, d). We tested these predictions using single-cell microscopy in the mother machine microfluidic device. Six P*_sigV_*-*YFP* reporter strains, each containing different copy numbers of *rsiV* (and therefore different *D* levels), were subjected to 1 µg/mL lysozyme stress. As the copy number of *rsiV* increased, the trend of σ^V^ ON-state activity decreased as expected, while its heterogeneity remained, consistent with the model’s predictions. At the highest *D* levels, σ^V^ activity was entirely suppressed (Figure 4c,e, Supplementary Figures 34, 35).

**Figure 4.**
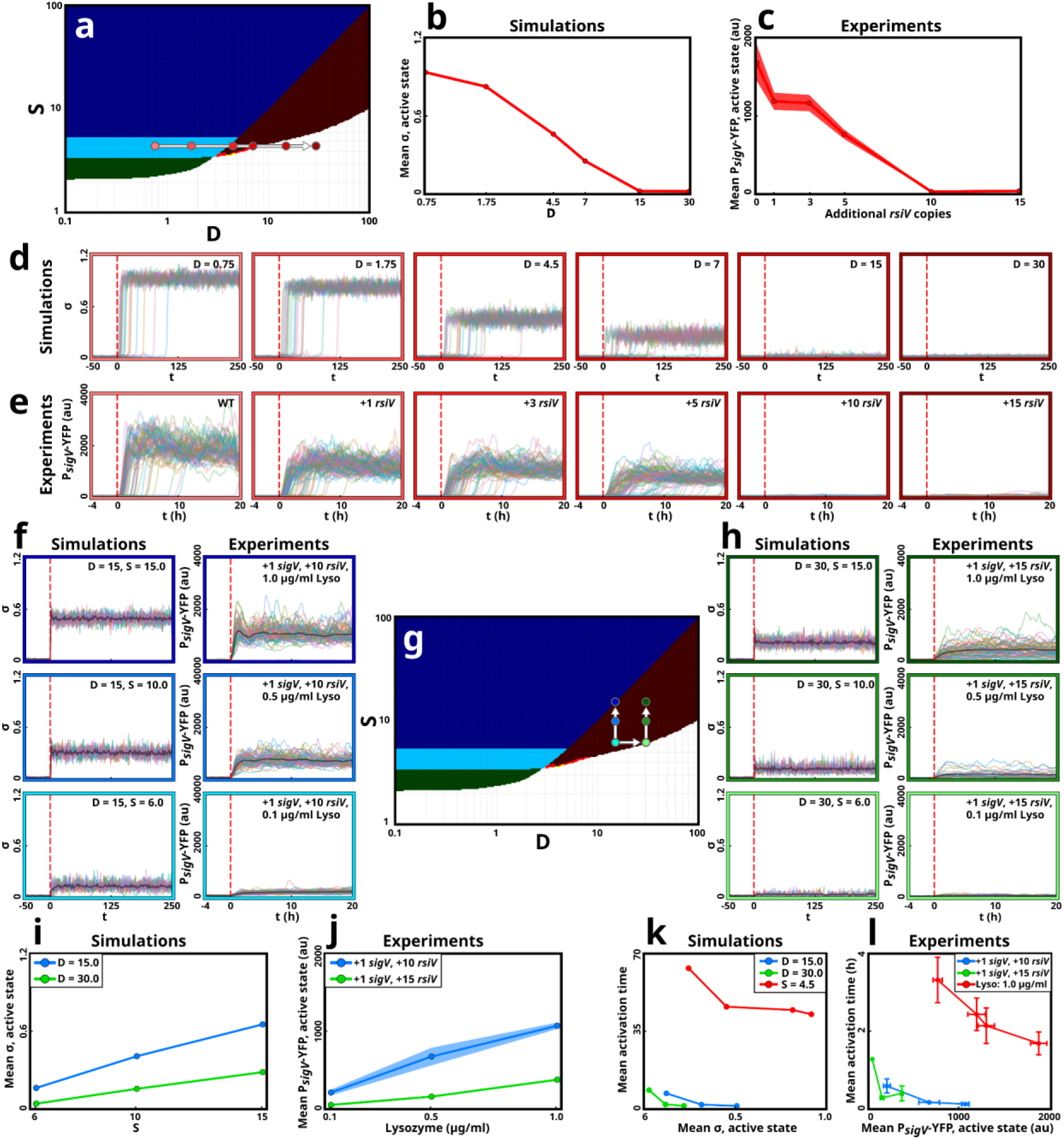
The model’s predicted transition between the heterogeneous and intermediate activation behavioural domains are validated in the σ^V^ system. The model predicts that, as D is increased, σ’s steady state activity is reduced while its heterogeneous activation remains. For high enough D levels, the system is fully inactivated. Here, it exhibits an intermediate activation transition as S is varied (unlike the heterogeneous activation transition previously observed for low D levels). **(a)** Six representative D values sampled from the model’s *S*-*D* behaviour map. **(b)** Simulated ON-state mean *σ* levels for the six points marked in a declines as *D* increases. **(*c*)**. For six different experimental *D* values, mean P_sigV_-YFP expression after 1 µg/mL lysozyme stress are shown. This confirms the decrease in ON-state activity with increasing *D* as predicted by the model. (**d**) Single-cell simulation traces (*n = 20*) for each condition in **a**. **(e)** Experimental single-cell fluorescence trajectories under 1 µg/mL lysozyme, recapitulating the simulated shift from heterogeneous activation to inactivity. **(f)** For fixed *D = 15*, the model (left) shows an intermediate activation transition as *S* is increased (simulations for *S = 6, 10, 15*). The ON-state activity increases with *S*. Experimental single-cell traces (right) for three lysozyme concentrations (0.1, 0.5, 1 µg/mL) validates this transition in a high-*D* σ^V^ strain (which carries 10 additional *rsiv* and one additional *sigV* copy). **(g)** The six *S*-*D* combinations used in **f** and **h**’s location in parameter space. **(h)** As *D* is increased (to *30*), the model (left) predicts a similar transition as in **f**, but with lower ON-state activity. We replicate this *D* increase by adding 5 *rsiV* copies to the strain used in **f**. Just like the model, this strain (right) replicates the transition in **f**, but with reduced ON-state activity. **(i,j)** Mean ON-state activity levels, as depending on *S*, for the simulations **(i)** and experiments **(j)** shown in **f** and **h**. **(k,l)** In the model **(k)** and experiment **(l)** the intermediate activation transitions (red and green lines) exhibit homogeneous activation, unlike the transition in **a** (red lines) which activates heterogeneously. All experiments have n=3-4 independent repeats.

Finally, we examined the σ^V^ system’s behaviour once it has transitioned into the *D≲1* regime. According to our model, this regime supports two possible *S*-transitions depending on the values of the remaining parameters *τ*, *v_0_*, *n*, and *η*, with *τ* having the primary effect (Supplementary Notes 8). These transitions include either pulsing/oscillatory behaviours, or intermediate activation (Figure 3, Supplementary Figures 28-32).

To identify in which region of parameter space the system is in, we modulated *S* in our two high *D* strains by adding an additional copy of *sigV* driven by its own promoter and by varying lysozyme stress levels. We found that σ^V^’s transition specifically falls into the intermediate activation category (Figure 4f-h). Our model predicts that the intermediate activation behaviour’s steady state activity is smoothly tuned by the levels of *S* and *D,* with *S* increasing activity whilst D reduces it (Figure 4i). We confirmed this experimentally (Figure 4j, Supplementary Movie 1). We also note that activation times are homogeneous in both simulation and experiment, further suggesting that the system has transitioned out of the heterogeneous activation domain observed for lower *D* values (Figures 4k-l). Together, these results confirm our prediction that, by tuning system parameters, a mixed feedback loop can transition between dynamically distinct behavioural regions.

### The predicted pulsing/oscillatory behavioural transition is validated in the σ^B^ system

Next, we experimentally tested the prediction from Figure 3 that increasing *S* can drive the system from stochastic pulsing into sustained oscillations. To do so, we used σ^B^, the master regulator of the general stress response in *B. subtilis* (*55–57*). Under persistent energy stress, σ^B^ exhibits stochastic pulsing (8,31). Our parameter maps predict a range of behaviours that can be generated from stochastic pulsing, such as oscillations, by tuning the value of *S*.

Normally, observation of potential high-*S* behaviours (such as oscillations) by tuning energy stress levels is not possible in σ^B^, as high energy stress causes secondary effects such as cell death. To avoid this issue, we rewired the σ^B^ circuit to synthetically drive σ^B^ via its energy stress pathway, adapting an approach previously used to re-wire σ^B^ via the environmental stress pathway (31). In a *B. subtilis* strain containing a reporter for σ^B^ activity, P*_sigB_*-*YFP*, we first knocked out *rsbP* and *rsbU*, the upstream phosphatases that activate the anti-anti-sigma factor RsbV under stress (Methods). Next, we chromosomally integrated a mutated form of RsbP, which is constitutively active, under the control of an IPTG inducible promoter to drive σ^B^ expression (Supplementary Figure 4). The mutated RsbP had the PAS and coiled-coil region removed causing it to be constitutively active (58).This enabled us to experimentally vary *S* by adding IPTG to the growth medium, allowing us to investigate the system’s response across a range of values.

We scanned our parameter space maps to determine which transition the system undergoes, given it starts in stochastic pulsing (the behaviour found in σ^B^), as *S* is modulated (Supplementary Notes 8). We note that stochastic pulsing is almost exclusively preceded (for lower *S*) by no activation, and proceeded (for higher *S*) by oscillation (Supplementary Tables 5). In addition, for high enough *S*, the system activates homogeneously. Finally, for some systems, the oscillating and the homogeneous activation regions are separated by a region of stochastic anti-pulsing (Figure 5a, Supplementary Tables 5, 6, Figures 28-32).

**Figure 5.**
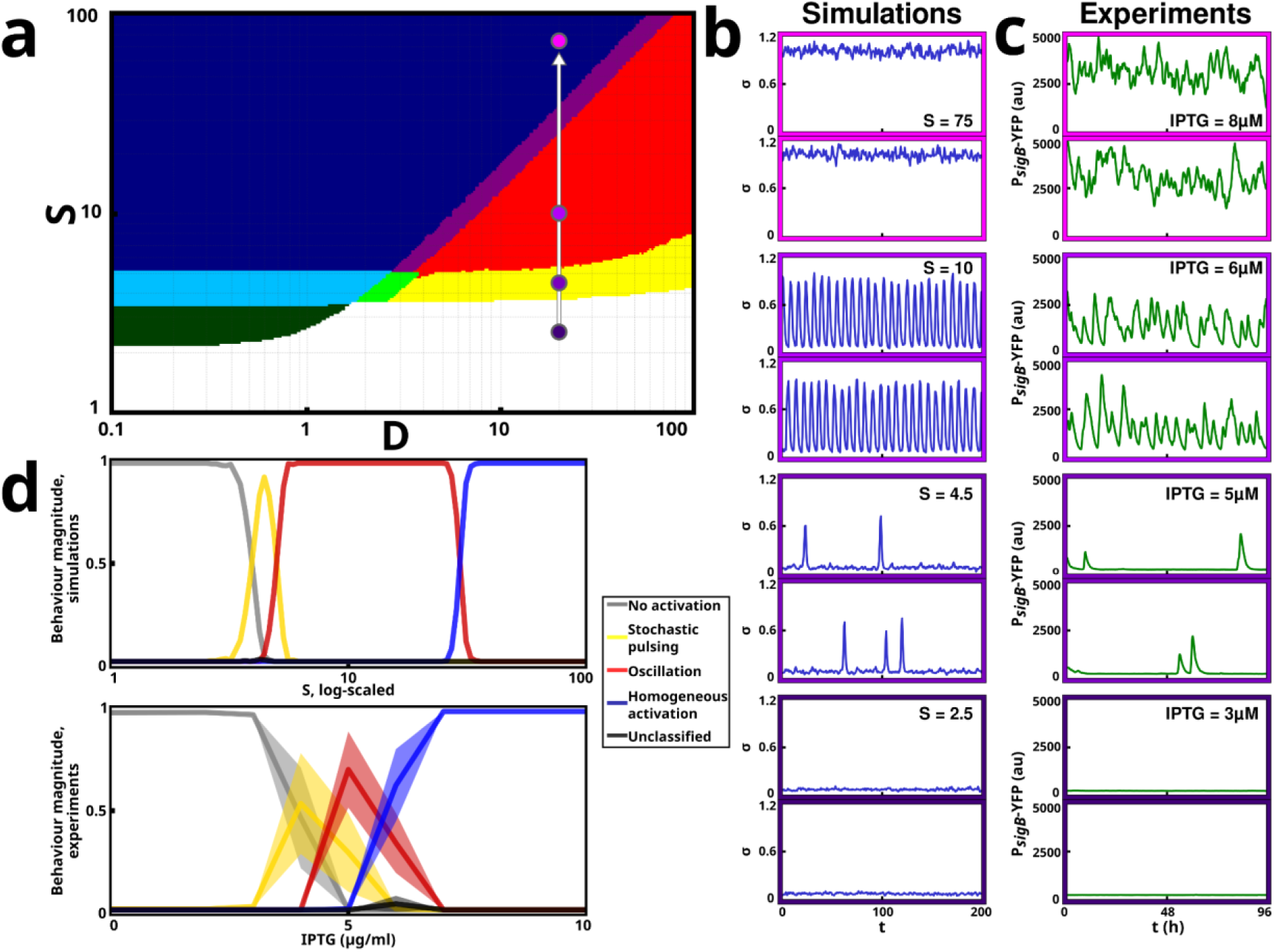
The model’s predicted transition through the pulsing/oscillatory region is validated in the σ^B^ system. **(a)** A behaviour map in which increasing self-activation strength (S) in the D ≲ 1 region drives a transition from stochastic pulsing to oscillations. **(b)** Simulated trajectories at four different S values (purple dots in **a**) under constant input, illustrating the predicted progression through no activation, stochastic pulsing, oscillations, and homogeneous activation. **(c)** Representative single-cell traces of the rewired B. subtilis P_sigB_-YFP strain (Methods) at increasing IPTG concentrations, showing the same behavioural sequence as in the model. **(d)** A classifier applicable to both simulation and experiment was developed to assign time-trajectories to one of four classes: no activation, stochastic pulsing, oscillation, or homogeneous activation (Supplementary Notes 11). Applied to simulated (from the parameter slice in **a**) and experimental (2–9 µg/mL IPTG) trajectories, the classifier confirms that both systems exhibit the same transition sequence. Shaded regions show standard deviation over n = 3-5 experimental replicates. The “unclassified” behaviour (black) indicates the (very few) trajectories the classifier failed to assign to either one class.

Next, we measured the σ^B^ activity of our rewired P*_sigB_*-*YFP* strain in the mother machine microfluidics device across various concentrations of IPTG (Methods). As IPTG levels increased from low to high, the σ^B^ system transitioned through no activation, stochastic pulsing, oscillation, and homogeneous activation behaviours, similar to as predicted by the model (Figure 5b, c, Supplementary Figures 36-39, Supplementary Movie 2). To further validate these findings, we developed a new classifier capable of quantifying these four behaviours’ occurrences in both model simulations and experimental time trajectories. Unlike the original classifier, which relies on analytical steady-state analysis of parameter values and cannot be applied to experimental data, this new classifier works directly with time-series data (Supplementary Notes 11, Supplementary Figures 40, 41). Applying it to both model and experimental data revealed that they undergo the same transition (Figure 5d).

We note that the stochastic anti-pulsing behaviour predicted by the model is not reproduced experimentally. This discrepancy could be explained by the half-life of the fluorescent marker for σ^B^ not providing high enough time resolution to distinguish the anti-pulses, or by additional complexity of the σ^B^ pathway enabling a different transition than what is predicted by our minimal model. Furthermore, the stochastic anti-pulsing behaviour only occurs for some combinations of *τ*, *v_0_*, *n* and *η*, while for others, oscillation transitions directly to homogeneous activation. While the former transition is about six times more common, we cannot here confirm which one σ^B^ exhibits (Supplementary Tables 5, 6, Figures 28-32).

Notably, the transition from stochastic pulsing to oscillations and then to homogeneous activation, as predicted by our minimal model and supported by experimental observations, also matches the transition identified in a previous stochastic model of the full σ^B^ circuit (59).

## Discussion

In this work, we present a general model of a mixed positive/negative feedback loop, which defines ten distinct dynamic behaviours this motif can generate (Figure 2). By employing an automated classification scheme, we mapped the occurrence of these behaviours across parameter space, revealing how each behaviour depends on the system’s properties (parameters) (Figure 3). To demonstrate our model’s applicability to real systems, we apply its predictions to bacterial sigma factors, a group of commonly studied systems whose circuits contain mixed positive/negative feedback loops.

We divide parameter space into two approximate regions, *D≲1* and *D≲1*, each undergoing a distinct behavioural transition as the input (or self-activation) parameter, *S*, is varied (Supplementary Figure 27). For *D≲1* the transition is characterised by bistable behaviours, while for *D≲1* a range of pulsing and oscillatory behaviours are exhibited. To validate our model, we examined the behaviours of two real systems (the alternative sigma factors σ^V^ and σ^B^, both from *B. subtilis*) as *S* is increased from small to large. Here, both systems undergo the predicted transition (Figures 4, 5). Furthermore, we showed that by increasing the value of *D*, a system in the *D≲1* region (e.g σ^V^) can be induced to produce *D≲1*-type behaviours. This shows that these two regions, as predicted by the model, can coexist in the same system (Figure 4).

The mixed positive/negative feedback loop is a common motif in cellular pathways. Our work provides a comprehensive understanding of how these systems generate a wide range of behaviours. A detailed analysis of each behaviour and their dynamics is provided in Supplementary Notes 7. Specifically for bacterial sigma factors, several behaviours predicted by our model, including single response pulse, stochastic pulsing, and heterogeneous activation, have been observed using single-cell microscopy (30,32,33,60). We demonstrate that these behaviours can be explained by the core mixed positive/negative feedback loop inherent in these systems (Figure 1c). Crucially, our model enables both forward and inverse predictions: it can predict which behaviours a circuit will exhibit given its properties, and conversely, infer likely circuit properties from observed behaviours. For example, the σ^B^ circuit, which displays stochastic pulsing, is predicted to operate at a higher self-deactivation strength (*D*) than the σ^V^ circuit, which displays heterogeneous activation.

As more sigma factor systems are explored, our model predicts that additional predicted behaviours may also be observed. Beyond sigma factors, our findings may generalise to mixed feedback loops in other biological and non-biological systems, including electronic control circuits, ecological models, and neural networks (12,61–63).

A central advantage of our approach is the use of a minimal, coarse-grained model to capture the essential dynamics of gene regulatory circuits. By not reproducing the detailed biochemistry of specific systems, our model allows us to make conclusions that can be generalised across mixed feedback loops. Furthermore, by testing these predictions *in vivo*, we confirm that our analysis holds when applied to real systems containing additional complexity, supporting the use of minimal models to make biologically relevant predictions. Finally, we show that noise not only enables qualitatively distinct dynamic behaviours, such as stochastic pulsing and heterogeneous activation, but also causes non-intuitive shifts of the boundaries between behavioural regimes in parameter space, effectively ‘warping’ the behavioural boundaries (Supplementary Figure 33).

Our insights are also relevant to synthetic biology, where mixed positive/negative feedback loops, such as those involving sigma factor circuits, have been proposed as potential regulators of synthetic systems (64–66). Here, we have only experimentally tuned two of the six parameters (*S* and *D*). Furthermore, the large number of additional genes required to tune *D* limits the practicality of our approach for most applications. In the future, it will be important to explore whether synthetic mixed positive/negative feedback loops that allow for tuning additional properties can be developed. Here, our work provides a blueprint for predicting and engineering circuit behaviours from minimal models, offering a foundation for designing synthetic regulators with tailored dynamic responses.

## Materials and Methods

### Mathematical model

Our minimal model of a general mixed positive/negative feedback loop tracks the concentration of a single component (*σ*). Degradation/dilution was assumed to be linear. To make production as general as possible, it was modelled as a Hill function. The model is based on the following three assumptions:

- *σ* activates its own production.
- *σ* deactivates its own production.
- The deactivation is subject to a time delay.

To take these into account, production depended on both an activating term (*σ*) and a repressing term (*σ* subject to a time delay).

The time delay was implemented using the linear chain trick (LCT) (52,53). While this introduces additional variables to the system, it has the advantage that the model can be represented as ODEs and SDEs (instead of DDEs and SDDEs). After confirming that the number of intermediaries used in the LCT had little effect on the model, we decided to implement it using three intermediaries (Supplementary Figures 7, 8, Supplementary Notes 4.1). Finally, we also confirmed that all ten response behaviours could be recreated in a model with a discrete delay (Supplementary Figure 9, Supplementary Notes 4.2).

Initially, our model depended on 9 parameters. Since we only sought to investigate qualitative behaviours, nondimensionalisation was performed. This section will describe the nondimensionalised model. The nondimensionalisation process and the non-nondimensionalised model are described in Supplementary Notes 2.

#### Reactions

Initially, our model contains only two reactions, production and degradation (or dilution) of the sigma factor (*σ*, the model’s only species). An additional three production and degradation reactions are added by the LCT (which also adds three intermediary species, *A_1_*, *A_2_*, and *A_3_*). These reactions are described in Table 1.

**Table 1:**
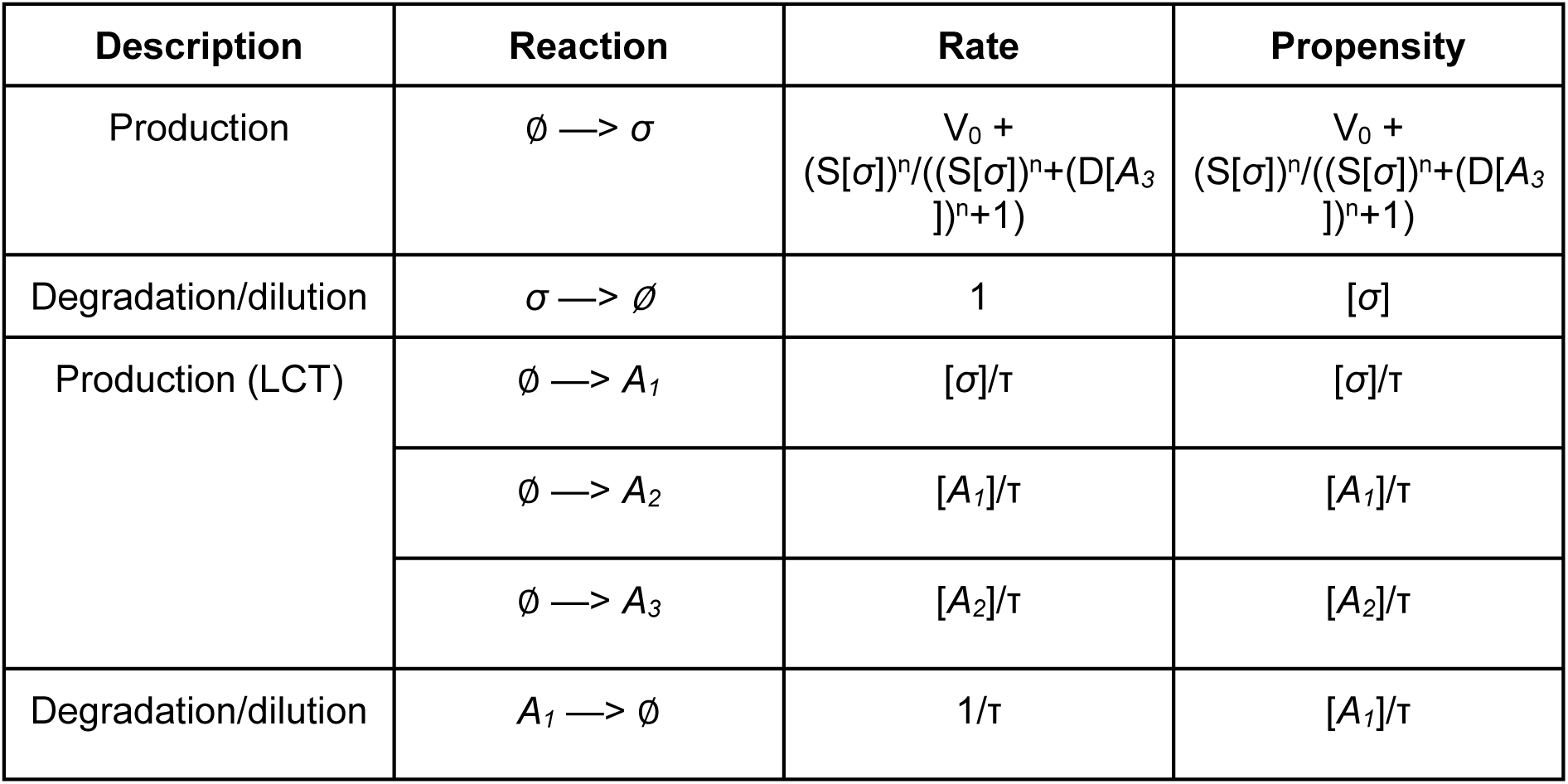
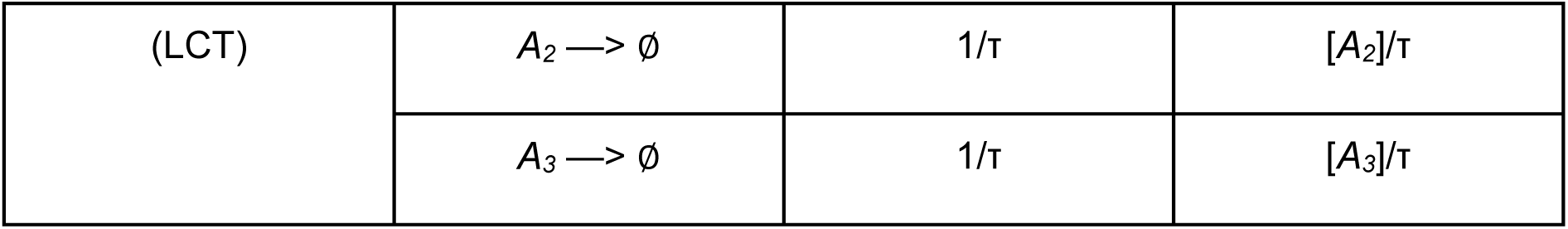
The reactions of the general sigma factor circuit model. The model primarily consists of two reactions, production and degradation of the sigma factor (σ). However, since the time delay (of the negative feedback of σ on its own production) is implemented using the LCT, an additional 3 species (A_1_, A_2_, and A_3_) are introduced. This also adds an additional three production and three degradation reactions.

#### Parameters

Our nondimensionalised model consists of 6 parameters, as described in Table 2. The model does not have specific parameter values. Instead, every instance of the model, with specific values of *(S*,*D*,*τ*,*v_0_*,*n*,*η)* represents a potential mixed positive/negative feedback loop. E.g. by letting *S* be large and *D* small, we can investigate the behaviour of circuits with high degrees of self-activation and low degrees of self-deactivation. As we scan these parameters’ values we can investigate how systems’ properties affect their behaviours. These scans are made possible through the nondimensionalisation process, which reduces the dimension of parameter space.

**Table 2:** The parameters of the (non-dimensionalised) general sigma factor model.

| Parameter | Description |
| --- | --- |
| $S$ | Degree of self-activation (also includes input strength). |
| $D$ | Degree of self-deactivation. |
| $\tau$ | Length of the self-deactivation time delay. |
| $v_0$ | Operon base activity/leakage. |
| $n$ | Hill coefficient (degree of non-linearity in the circuit). |
| $\eta$ | Degree of noise in the circuit. |

We note that the parameter *S* is a composite of the pre-nondimensionalised parameters *S*^ and *I*^, where *S*^ is the degree of system self-activation, and *I*^ the magnitude of the input signal (Supplementary Notes 2). Hence, modulation of a system’s input corresponds to modulation of the parameter *S*.

### Model Simulations

We wish to investigate the model’s behaviour in response to an input. In the absence of the input (in response to which we wish to investigate the response), *S=0*. When we simulate our model for a specific parameter set *(S*,*D*,*τ*,*v_0_*,*n*,*η)*, we will initialise it with *S=0*, with the parameter *S* only attaining its designated value at the time of input onset (always at time *t=0*). E.g. a model instance may be defined by the parameter set *(S*,*D*,*τ*,*v_0_*,*n*,*η)=(5.0,5.0,10.0,0.1,2,0.1)*, and simulated throughout the time interval (*-500,2000*). The simulations start with the parameter values *(0.0,5.0,10.0,0.1,2,0.1)*, which are changed to *(5.0,5.0,10.0,0.1,2,0.1)* at time *0*, at which point the system activates.

It would be possible to vary the value of *S* throughout a simulation, to model inputs with varying magnitude. However, for simplicity, this work will only consider systems with a step increase in the input.

Practically, the model was implemented using in the Julia programming language using the Catalyst.jl modelling package (67). All simulations were carried out using the DifferentialEquations.jl package (68). The ImplicitEM solver was used as it handles stiff SDE systems well, and allows for adaptive time stepping (69). Stochastic chemical kinetics were simulated using Gillespie’s direct algorithm using the JumpProcesses.jl package (70).

### Automated behaviour classification

Initial steady state and stability analysis of our model determined that it may have either 0, 1, or 2 stable steady states (Supplementary Notes 5). While a dynamic system asymptotically approaches a stable steady state, perturbations may cause a transition towards another steady state. Such perturbations may either be transient (change in input magnitude) or asymptotic (noise). Based on the possible transitions between the system’s potential steady states, we designed a classification scheme for the system’s behaviour (Supplementary Notes 8). This scheme takes into account the possibility of excitable and oscillatory behaviours. A total of 10 system behaviours (as described in the Results) were predicted.

Further behaviours could be created by subdividing the initial ten, however, we deemed such subdivisions non-beneficial (Supplementary Notes 6.4). Further description and analysis of the individual behaviours can be found in Supplementary Notes 7.

To enable our mapping of behaviour across parameter space, we created a decision tree-based heuristic classifier algorithm. When provided a parameter set *(S*,*D*,*τ*,*v_0_*,*n*,*η)*, the algorithm simulates the model multiple times, and assigns it one of the behavioural classes described in Figure 2. The full algorithm is described in Supplementary Notes 8. In addition, to further prove its validity, we randomly generated 10 parameter sets from each behaviour (as classified by the algorithm). By showing that simulations of these parameter sets align with their designated behaviours, we demonstrate the efficacy of the algorithm (Supplementary Figures 15, 16).

### Bifurcation analysis

Bifurcation diagrams were generated using the BifurcationKit.jl software (71).

### Experiments

#### Strains and Media

All strains are derived from *B. subtilis* 168 or PY79. Deletions were made by recombination of linear DNA fragments homologous to the region of interest and replacing the gene of interest with an antibiotic resistance cassette. All strains had a housekeeping σ^A^ promoter-driven mCherry as a constitutive control and for segmentation.

Cells were grown in Spizizen’s Minimal Media (SMM) (72). It contained 50 µg/ml Tryptophan as an amino acid source and 0.5% glucose as a carbon source. Cultures were started from frozen stock in SMM and grown overnight at 30°C to an OD between 0.3 and 0.8. The overnight cultures were resuspended to an OD of 0.01 and regrown to an OD of 0.1 at 37°C.

#### Plasmids

E. coli strain DH5α was used to clone all plasmids. The cloning was done with a combination of standard molecular cloning techniques using Clonetech In-Fusion Advantage PCR Cloning kits and non-ligase dependent cloning. Plasmids were chromosomally integrated into the 168 or PY79 background via double crossover using standard techniques. The list below provides a description of the used plasmids, with details on selection marker and integration position/cassette given at the beginning. Note that all plasmids below replicate in *E. coli* but not in *B. subtilis*.

1. ppsB::P*_trpE_* - *mCherry* ErmR This plasmid was used to provide uniform expression of mCherry from a σ^A^-dependent promoter, enabling automatic image segmentation (cell identification) in time-lapse movie analysis. A minimal σ^A^ promoter from the *trpE* gene was cloned into a vector with *ppsB* homology regions (31). The original integration vector was a gift from A. Eldar (73).
2. sacA::P*_sigB_*-*YFP* CmR The target promoter of *sigB* was cloned into the EcoRI/BamHI sites of AEC127 (73) yielding a Venus (YFP) reporter for σ^B^ activity (gift from M. Elowitz, CalTech).
3. amyE::P*_hyperspank_*-*rsbP*Δ4 (ΔPAS-CC) The coding region of a *rsbP* mutant lacking PAS and the coiled-coil region along with a 5’ transcriptional terminator, was cloned downstream of the hyperspank IPTG-inducible promoter in plasmid pDR-111 (gift from D. Rudner, Harvard). *rsbP*Δ4 (ΔPAS-CC) is described in (58).
4. sacA::P_sigV_-YFP CmR The target promoter of *sigV* was cloned into the EcoRI/BamHI sites of AEC12 (Eldar *et al,* <u>2009</u>) (gift from M. Elowitz, CalTech).
5. *amyE*::P*_sigV_* -? x *rsiV* SpecR The target promoter of *sigV* with multiple copies of *rsiV* (?= 1x, 3x, 5 x) were cloned into the sites EcoRI/BAmHI and BamHI/HindIII of pdL30 (Locke et al, 2011).
6. *lacA*::P*_sigV_* - 5 x *rsiV* TetR The target promoter of *sigV* with five copies of *rsiV* was cloned into the sites Ecorl/Nde of a variation of ECE174 (gift from M. Elowitz, CalTech).
7. *pyrD*::P*_sigV_* - 5 x *rsiV* NatR The target promoter of *sigV* with five copies of *rsiV* was cloned into the sites EcoRI/BAmHI of a variation of ECE174 (gift from M. Elowitz, CalTech).
8. hag::P_sigV_-sigV ErmR The target promoter of *sigV* driving *sigV* and 400 upstream and downstream homology regions of hag was cloned into EcorV of pBS II SK

#### Strain List

**Table 3:**
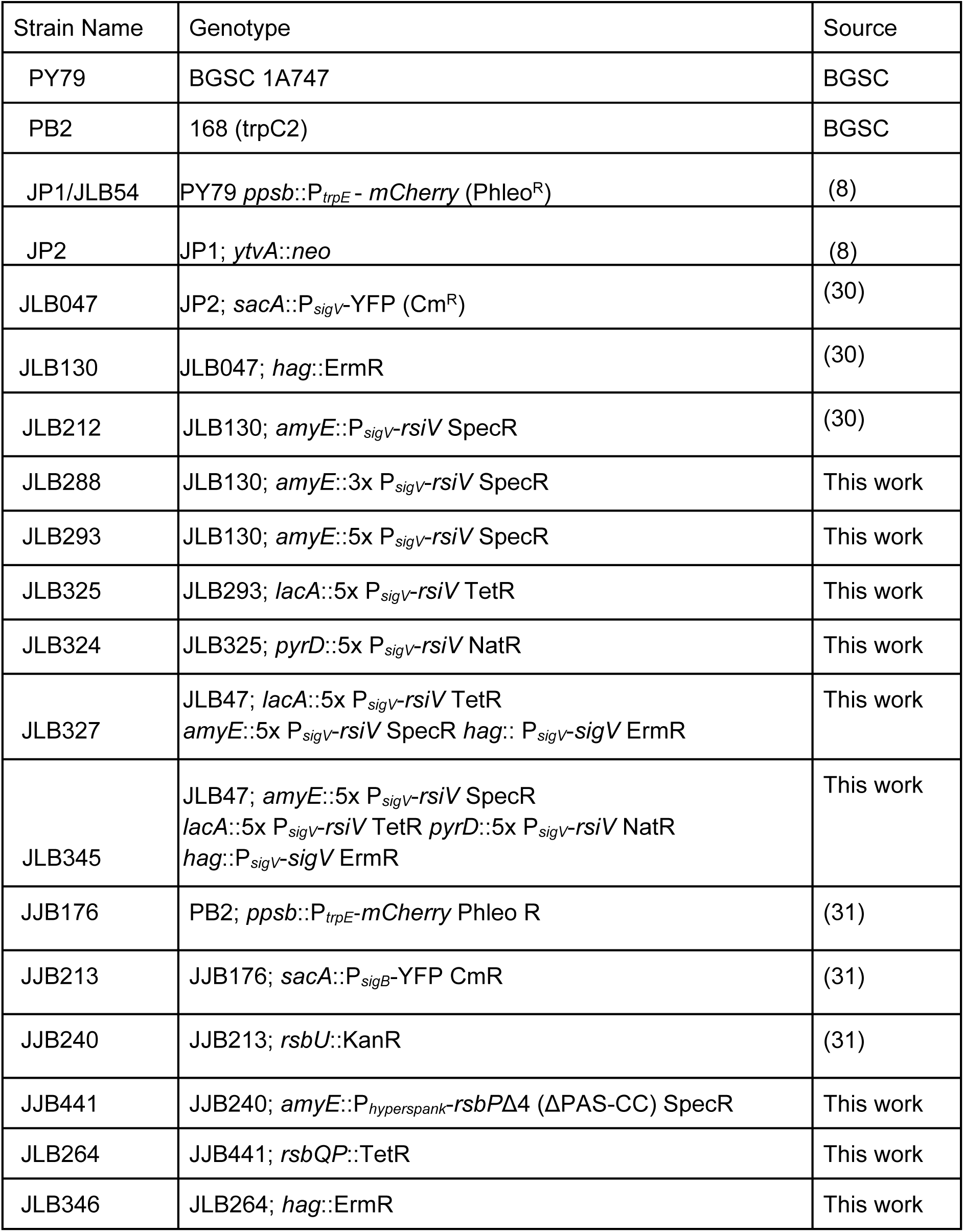
List of strains used in this article.

#### Microscopy

An inverted Nikon Ti-E microscope (Nikon, Netherlands) with a Photometrics Prime sCMOS camera (Photometrics, USA) and a 100x phase Plan Apo (NA 1.4) objective (Nikon, Netherlands) were used for all experiments. Brightfield illumination was provided by a LED lamp (CoolLED, UK) and for Epifluorescence illumination a Lumencore Solar II light engine (Lumencore USA) was used. Chroma filters (Chroma, Bellows Falls, USA) #41027 for the RFP channel and #49003 for the YFP channel were used in combination with the epifluorescence. For temperature control a stage incubator (Solent Scientific, UK) was set at 37°C for all experiments. Metamorph (Molecular Device, Sunnyvale, CA, USA) controlled the camera, the motorised stage (Nikon, Amsterdam, Netherlands) and the microscope.

#### Microfluidic Designs

Two microfluidic designs were used in this study. All experiments on σ^B^ were done in a traditional mother machine device (29). The master for these chips was fabricated by exploiting a Kloe Dilase 650, direct laser writer (Kloe, France). More details on the fabrication can be found in the Supplementary Notes 12. All experiments on σ^V^ were done in a mother machine device with side trenches (74). This epoxy master was a kind gift from the Ellowitz lab.

#### Microfluidic Chip fabrication

All microfluidics chips were made using Sylgard 184 polydimethylsiloxane (PDMS) (Dow Corning, USA) using a 10:1 base to curing agent ratio. The PDMS was poured onto the silicon master and cured at 65C for 2h. Chips were cut out and inlets were punched into the PDMS using a 1 mm biopsy puncher (WPI, UK). The chips were then plasma bonded to glass bottom dishes (Wilco Wells, Netherlands) using a Femto Plasma System (Diener, Germany) (12s at 30W and 0.35 mbar). To strengthen the bonding of the PDMS to the glass dish the chips were put into the oven for 10 min at 65°C.

#### Microfluidic Experiments

10 ml of the regrown overnight culture with an of OD 0.1 were spun down at 4000 rpm (3000 g) for 10 min and resuspended in 100 μl. This cell suspension was loaded into the feeding channels and spun for 5 min at 3000 rpm using a spin coater (Polos SPIN150i, SPS) to improve the loading of the cells into the growth channels.

The loaded chips were then mounted onto the microscope stage and connected to a syringe pump (KD Scientific, USA) using tygon tubing (Cole Parmer, USA). A constant flow rate of 0.15 ml/h and a temperature of 37°C was maintained throughout the experiment. In all experiments with P_sigV_-YFP cells were first grown in SMM overnight for ∼100 frames before the media was switched to contain various levels of lysozyme (0.1, 0.5 or 1 μg/ml). Switching was provided by disconnecting the tubing from media with 0 ug/ml lysozyme and connecting the tubing to a new syringe with lysozyme stress. Subsequently, the tubing was flushed for 10 min at 10 ml/h.

All movies were analysed using a modified version of the Schnitzcells package (75) for MATLAB (Mathworks, USA) optimised for mother machine experiments.

## Data availability

All data used in this paper, as well as all code for carrying out all analysis, and the generation of all figures, can be found here: https://gitlab.developers.cam.ac.uk/slcu/teamjl/loman_schwall_etal_2025. Repository README files and code comments direct users on its use. Julia’s default package manager can be used to reproduce and install the exact package environment used. The repository contains the 6d behaviour maps, which is available for download and further investigation.

## Acknowledgments

We thank Ozgur Akman and Diana Fusco for their extensive feedback on the modelling. We thank Romain Veltz for helping with the computation of the bifurcation diagrams. We are grateful for the high performance computing resources provided by the Cambridge Service for Data Driven Discovery. This research was made possible by the award of a European Research Council under the European Union’s Seventh Framework Programme (FP/2007-2013)/ERC Grant Agreement 338060. The work in the Locke laboratory is further supported by a fellowship from the Gatsby Foundation (GAT3272/GLC), and a BBSRC grant award no. BB/V006088/1. Torkel Loman has received funding from the European Union’s Horizon 2020 research and innovation programme under the Marie Skłodowska-Curie grant agreement No. 721456.

