## Supplementary Information for "Mixed positive and negative feedback loops drive diverse single-cell gene expression dynamics"

### Loman & Schwall et. al. 2025 - Supplementary Material

### Contents

**1 Sigma factor circuits** **3**

**2 Model formulation and nondimensionalisation** **7**

**3 Replication of results using Gillespie's algorithm** **11**

**4 Modelling the time delay** **13**

#### 1 Sigma factor circuits

In this article, we validate our model using the $\sigma^V$ and $\sigma^B$ systems (which control the lysozyme and general stress response of *Bacillus subtilis*, respectively). Circuit diagrams for these can be found in Supplementary Figures 1 and 2 (for $\sigma^V$ and $\sigma^B$, respectively). Additionally, Supplementary Figures 3 and 4 describe the rewired circuits used for the validation experiments (for $\sigma^V$ and $\sigma^B$, respectively).

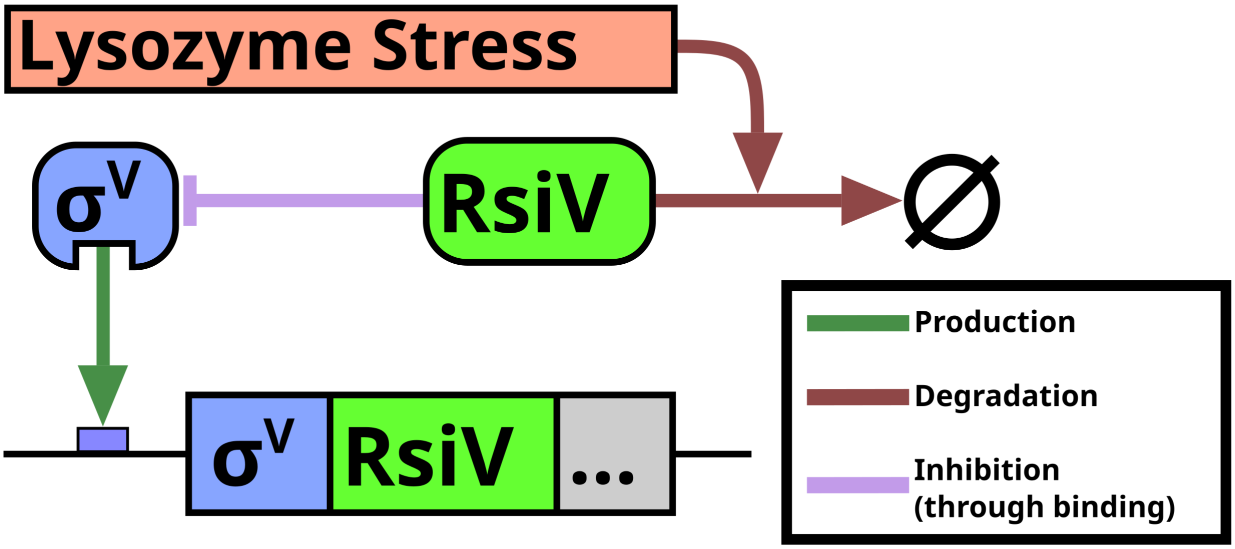

Supplementary Figure 1: **Simplified schematic of the $\sigma^V$ regulatory circuit highlighting the core mixed feedback motif.** Under non-stress conditions, $\sigma^V$ is bound to, and kept inactive by, its anti-sigma factor RsiV. Lysozyme stress will activate the degradation of RsiV, in the process of which $\sigma^V$ is released. Free $\sigma^V$ activates the genes required for the lysozyme stress response. It also activates the production of itself ($\sigma^V$), and its anti-sigma factor (RsiV), creating a mixed positive/negative feedback loop. Additional components involved in $\sigma^V$ activation are omitted here for clarity

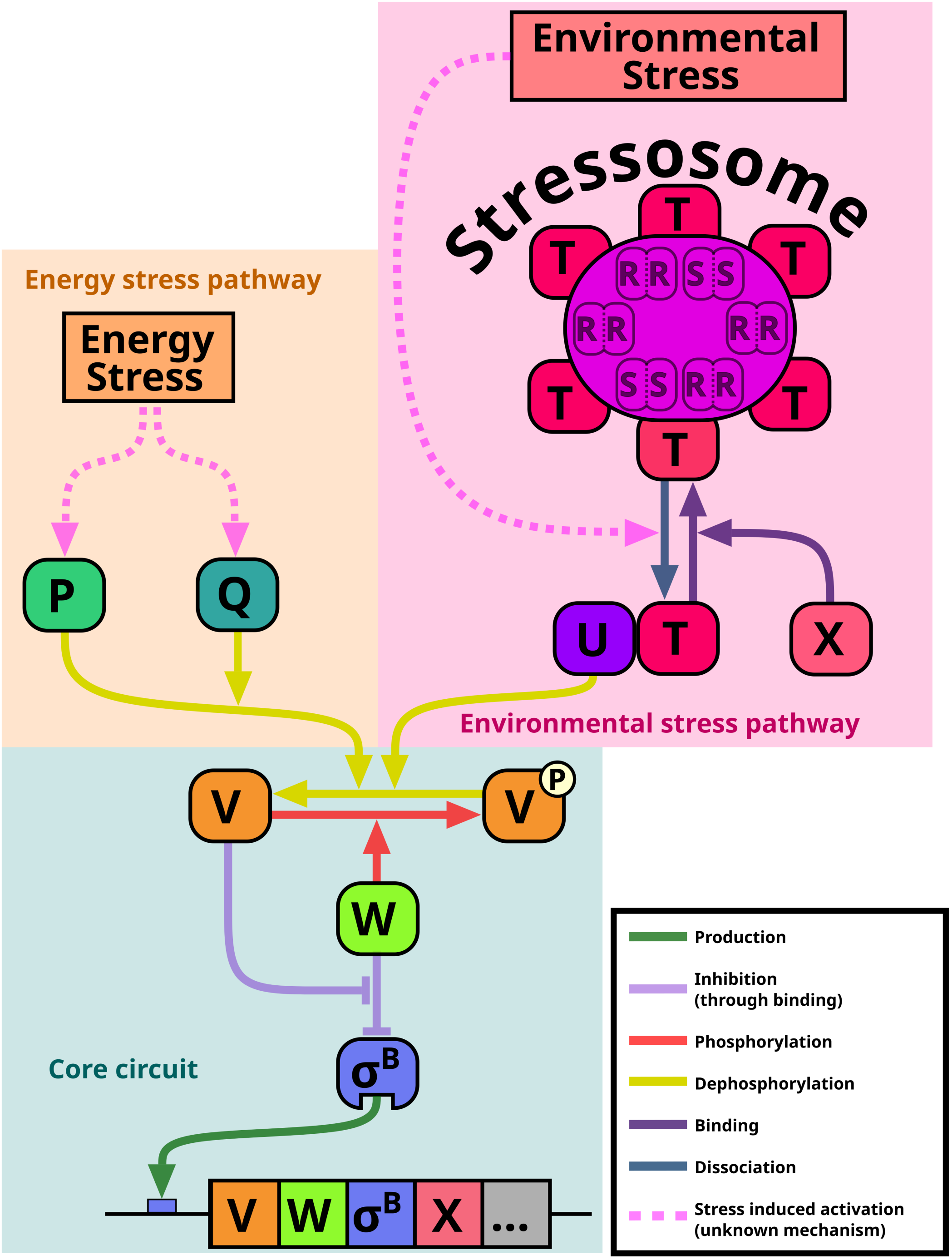

Supplementary Figure 2: **The $\sigma^B$ regulatory network consists of a core circuit, which is activated by two distinct upstream pathways.** Under non-stress conditions, $\sigma^B$ is bound, and held inactive by, its anti-sigma factor (RsbW, W in the figure). An anti-anti-sigma factor (RsbV, V in the figure) is inactivated by phosphorylation. The upstream pathways are triggered by two distinct types of stress (environmental and energy stress), each activating their respective phosphatase (RsbTU and RsbP, respectively). These then dephosphorylate, and thus activate, RsbV. Once activated, RsbV binds RsbW, which simultaneously releases $\sigma^B$ in a partner switching mechanism. This permits $\sigma^B$ to activate the general stress response of *B. subtilis*. In addition, $\sigma^B$ activates the production of itself, RsbW, and RsbV, creating a mixed positive/negative(/positive) feedback loop. The environmental stress response phosphatase, RsbTU, consists of RsbU and RsbT (U and T, respectively, in the figure). Under non-stress conditions, RsbT is bound to the stressosome protein complex. The stressosome also consists of RsbS and RsbR (S and R, respectively, in the figure) dimers. It senses environmental stress (via an unknown mechanism), in response to which RsbT is released. Free RsbT can bind RsbU and activate the core $\sigma^B$ circuit. In addition, RsbX (X in the figure), which is produced by active $\sigma^B$, facilitates the rebinding of RsbT to the stressosome. The details of the energy stress response are less well understood. The phosphatase RsbP (P in the figure), together with its associated factor RsbQ (Q in the figure), mediates $\sigma^B$ activation in response to energy stress. In this work, we only use a rewired version of the $\sigma^B$ circuit (where the WT energy and environmental stress sensing pathways have been disabled, Supplementary Figure 4).

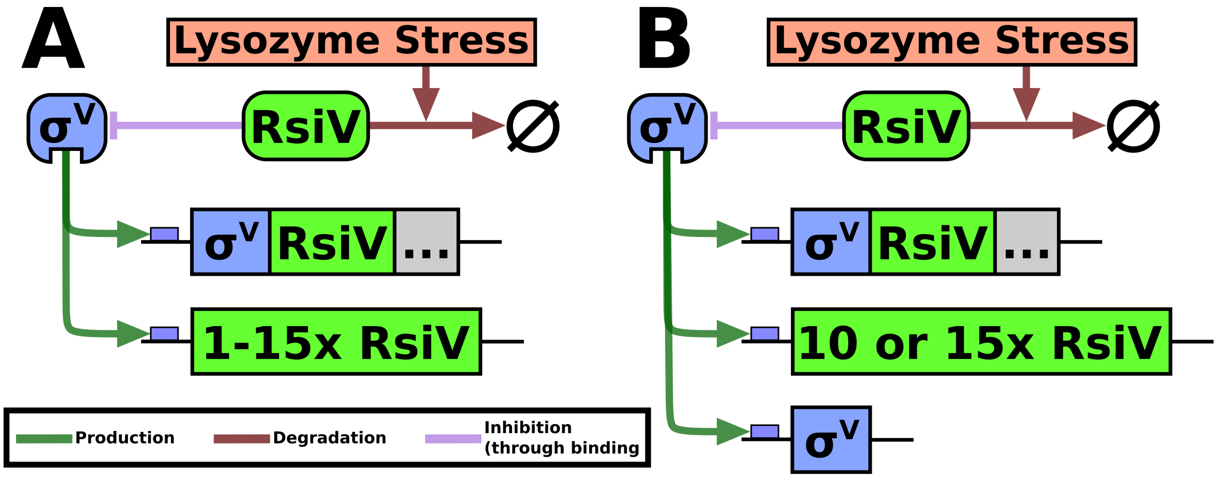

Supplementary Figure 3: **By rewiring the $\sigma^V$ circuit, in addition to $S$, we can control $D$ levels.** (A) We add an additional $P_{sigV}$-driven operon containing either 1, 3, 5, 10, or 15 copies of *rsiV*. By modulating the number of copies of *rsiV* we can control the circuit's self-deactivation strength, $D$. (B) To the strains in A with either 10 or 15 copies of *rsiV*, we add a single $P_{sigV}$-driven operon containing one copy of *sigV* only. This increases the circuit's self-activation strength, $S$. This enables us to reach high $S$ levels with less severe stress strengths (reducing secondary effects on the bacteria).

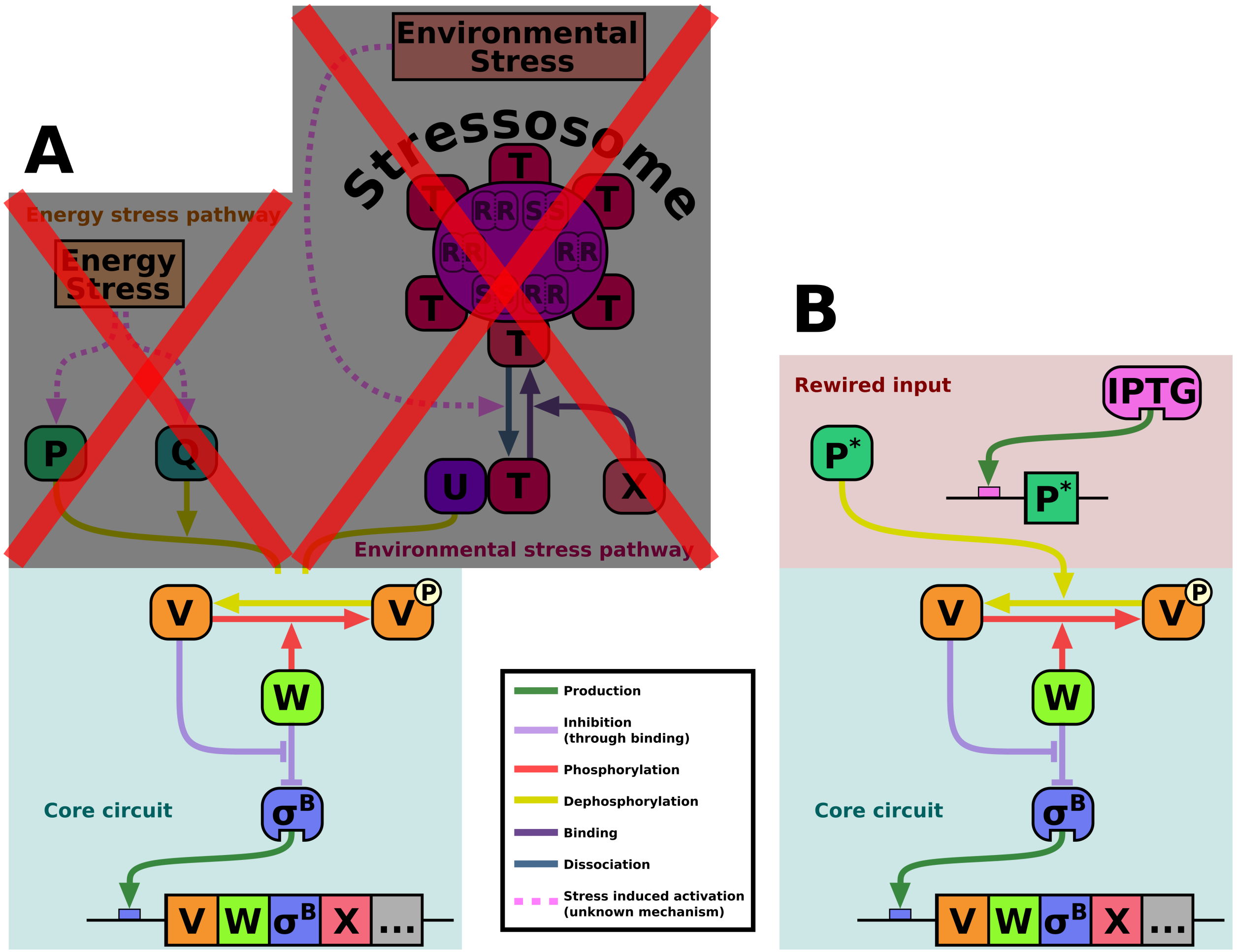

Supplementary Figure 4: **By rewiring the $\sigma^B$ circuit we can control $S$ levels.** (A) First, we delete the native energy and environmental stress response upstream pathways to $\sigma^B$. This prevents any wild-type activation of the circuit. (B) Next, we add a constitutively active RsbP ($P^*$) to the circuit. The production of $P^*$ is controlled by IPTG. This way, we can control the circuit input ($S$) by modulating the IPTG levels.

### 2 Model formulation and nondimensionalisation

Our model is a chemical reaction network (CRN) type model. It initially consisted of a single species ($\sigma$), which has a corresponding production reaction (modelled as a Hill function) and a (linear) degradation/dilution reaction. Furthermore, by implementing the time delay using the linear chain trick (LCT) (Supplementary Section 4.1) we introduce additional species and reactions. We chose to use three intermediaries in the LCT (Supplementary Section 4.1, which generates an additional three production and degradation reactions (all linear). All the reactions of the initial model is described in Supplementary Table 1.

| **Description** | **Reaction** | **Rate** | **Propensity** |
|---|---|---|---|
| Production | $\varnothing \longrightarrow \sigma$ | $\hat{v}_0 + \hat{v}\frac{(\hat{I}\cdot\hat{S}[\hat{\sigma}])^{\hat{n}}}{(\hat{I}\cdot\hat{S}[\hat{\sigma}])^{\hat{n}}+(\hat{D}[\hat{A}_3])^{\hat{n}}+\hat{K}^{\hat{n}}}$ | $\hat{v}_0 + \hat{v}\frac{(\hat{I}\cdot\hat{S}[\hat{\sigma}])^{\hat{n}}}{(\hat{I}\cdot\hat{S}[\hat{\sigma}])^{\hat{n}}+(\hat{D}[\hat{A}_3])^{\hat{n}}+\hat{K}^{\hat{n}}}$ |
| Degradation/dilution | $\sigma \longrightarrow \varnothing$ | $\hat{k}_{deg}$ | $\hat{k}_{deg}\cdot[\hat{\sigma}]$ |
| Production (LCT) | $\varnothing \longrightarrow A_1$ | $\frac{[\hat{\sigma}]}{\hat{\tau}}$ | $\frac{[\hat{\sigma}]}{\hat{\tau}}$ |
| | $\varnothing \longrightarrow A_2$ | $\frac{[\hat{A}_1]}{\hat{\tau}}$ | $\frac{[\hat{A}_1]}{\hat{\tau}}$ |
| | $\varnothing \longrightarrow A_3$ | $\frac{[\hat{A}_2]}{\hat{\tau}}$ | $\frac{[\hat{A}_2]}{\hat{\tau}}$ |
| Degradation/dilution (LCT) | $A_1 \longrightarrow \varnothing$ | $\frac{1}{\hat{\tau}}$ | $\frac{[\hat{A}_1]}{\hat{\tau}}$ |
| | $A_2 \longrightarrow \varnothing$ | $\frac{1}{\hat{\tau}}$ | $\frac{[\hat{A}_2]}{\hat{\tau}}$ |
| | $A_3 \longrightarrow \varnothing$ | $\frac{1}{\hat{\tau}}$ | $\frac{[\hat{A}_3]}{\hat{\tau}}$ |

Supplementary Table 1: **The reactions of the (non-nondimensionalised) general sigma factor circuit model.** Here, ˆ is used to denote variables and parameters before nondimensionalisation (it is, however, not used to modify the species names in the reactions). The reactions' propensities are formed by multiplying their rates with their substrates.

Before nondimensionalisation, the model consists of nine parameters. These are described in Supplementary Table 2.

Through the chemical Langevin equations (CLE) [Gillespie, 2000], the reactions in Supplementary Table 1 generate the following system of stochastic differential equations (SDEs):

$$\begin{cases} d[\hat{\sigma}] &= d\hat{t}\cdot\left(\hat{v}_0 + \hat{v}\frac{(\hat{I}\cdot\hat{S}[\hat{\sigma}])^{\hat{n}}}{(\hat{I}\cdot\hat{S}[\hat{\sigma}])^{\hat{n}}+(\hat{D}[\hat{A}_3])^{\hat{n}}+\hat{K}^{\hat{n}}} - \hat{k}_{deg}[\hat{\sigma}]\right) \\ &\quad +\sqrt{\hat{v}_0 + \hat{v}\frac{(\hat{I}\cdot\hat{S}[\hat{\sigma}])^{\hat{n}}}{(\hat{I}\cdot\hat{S}[\hat{\sigma}])^{\hat{n}}+(\hat{D}[\hat{A}_3])^{\hat{n}}+\hat{K}^{\hat{n}}}}\cdot d\hat{W}_1 - \sqrt{\hat{k}_{deg}[\hat{\sigma}]}\cdot d\hat{W}_2 \\ d[\hat{A}_1] &= d\hat{t}\left(\frac{[\hat{\sigma}]}{\hat{\tau}} - \frac{[\hat{A}_1]}{\hat{\tau}}\right) \\ &\quad +\sqrt{\frac{[\hat{\sigma}]}{\hat{\tau}}}\cdot d\hat{W}_3 - \sqrt{\frac{[\hat{A}_1]}{\hat{\tau}}}\cdot d\hat{W}_4 \\ d[\hat{A}_2] &= d\hat{t}\left(\frac{[\hat{A}_1]}{\hat{\tau}} - \frac{[\hat{A}_2]}{\hat{\tau}}\right) \\ &\quad +\sqrt{\frac{[\hat{A}_1]}{\hat{\tau}}}\cdot d\hat{W}_5 - \sqrt{\frac{[\hat{A}_2]}{\hat{\tau}}}\cdot d\hat{W}_6 \\ d[\hat{A}_3] &= d\hat{t}\left(\frac{[\hat{A}_2]}{\hat{\tau}} - \frac{[\hat{A}_3]}{\hat{\tau}}\right) \\ &\quad +\sqrt{\frac{[\hat{A}_2]}{\hat{\tau}}}\cdot d\hat{W}_7 - \sqrt{\frac{[\hat{A}_3]}{\hat{\tau}}}\cdot d\hat{W}_8 \end{cases}$$

where $d\hat{W}_i$ is a Wiener process. There exists one such noise term for each reaction (subscript correspond to order of the reactions in Supplementary Table 1). We will next describe the nondimensionalisation

| Parameter | Description | Unit |
|---|---|---|
| $\hat{v}_0$ | Transcriptional leakage of the operon. | µM·min$^{-1}$ |
| $\hat{v}$ | Maximal operon activity. | µM·min$^{-1}$ |
| $\hat{I}$ | Input signal magnitude. | - |
| $\hat{S}$ | Degree of system self-activation. | - |
| $\hat{D}$ | Degree of system self-deactivation. | - |
| $\hat{K}$ | Activation threshold. | µM |
| $\hat{n}$ | Hill coefficient (degree of non-linearity in the circuit). | - |
| $\hat{k}_{deg}$ | Degradation/dilution rate. | min$^{-1}$ |
| $\hat{\tau}$ | Rate parameter of the LCT (length of the time-delay). | min |

Supplementary Table 2: **The parameters of the (non-nondimensionalised) general sigma factor circuit model.** Here, ˆ is used to denote parameters before nondimensionalisation. Dimensions for the parameters are given using minutes for time, and µM for species amounts, however, these specifics will not be used further.

procedure where we reduce the number of parameters of this model, while preserving its dynamics. We start by considering the first SDE only:

$$d[[\hat{\sigma}] = d\hat{t} \cdot (\hat{v}_0 + \hat{v}\frac{(\hat{I} \cdot \hat{S}[\hat{\sigma}])^{\hat{n}}}{(\hat{I} \cdot \hat{S}[\hat{\sigma}])^{\hat{n}} + (\hat{D}[\hat{A}_3])^{\hat{n}} + \hat{K}^{\hat{n}}} - \hat{k}_{deg}[\hat{\sigma}]) + \sqrt{\ldots} \cdot d\hat{W}_1 - \sqrt{\ldots} \cdot d\hat{W}_2.$$

Here, we multiply both sides by $\frac{\hat{k}_{deg}}{\hat{v}}$ :

$$d\frac{\hat{k}_{deg}[\sigma]}{\hat{v}} = d(\hat{k}_{deg} \cdot \hat{t}) \cdot (\frac{\hat{v}_0}{\hat{v}} + \frac{(\hat{I} \cdot \hat{S}[\hat{\sigma}])^{\hat{n}}}{(\hat{I} \cdot \hat{S}[\hat{\sigma}])^{\hat{n}} + (\hat{D}[\hat{A}_3])^{\hat{n}} + \hat{K}^{\hat{n}}} - \frac{\hat{k}_{deg}[\hat{\sigma}]}{\hat{v}}) + \frac{\hat{k}_{deg}}{\hat{v}}(\sqrt{\ldots} \cdot d\hat{W}_1 - \sqrt{\ldots} \cdot d\hat{W}_2).$$

Next, we multiply both the numerator and denominator of the fraction by $\frac{1}{K^n}$. Within the fraction, multiply the three terms (not identical to 1) by $(\frac{\hat{v}}{\hat{v}})^{\hat{n}}$ and $(\frac{\hat{k}_{deg}}{\hat{k}_{deg}})^{\hat{n}}$. This yields:

$$d\frac{\hat{k}_{deg}[\sigma]}{\hat{v}} = d(\hat{k}_{deg} \cdot \hat{t}) \cdot (\frac{\hat{v}_0}{\hat{v}} + \frac{(\frac{\hat{I}\cdot\hat{S}\cdot\hat{v}}{\hat{k}_{deg}\cdot\hat{K}}\frac{\hat{k}_{deg}[\hat{\sigma}]}{\hat{v}})^{n}}{(\frac{\hat{I}\cdot\hat{S}\cdot\hat{v}}{\hat{k}_{deg}\cdot\hat{K}}\frac{\hat{k}_{deg}[\hat{\sigma}]}{\hat{v}})^{n} + (\frac{\hat{D}\cdot\hat{v}}{\hat{k}_{deg}\cdot\hat{K}}\frac{\hat{k}_{deg}[\hat{A}_3]}{\hat{v}})^{n} + 1} - \frac{\hat{k}_{deg}[\hat{\sigma}]}{\hat{v}}) + \frac{\hat{k}_{deg}}{\hat{v}}(\sqrt{\ldots} \cdot d\hat{W}_1 - \sqrt{\ldots} \cdot d\hat{W}_2).$$

In a similar manner, the noise terms can be rewritten as:

$$\frac{\hat{k}_{deg}}{\hat{v}}\sqrt{\hat{v}_0 + \hat{v}\frac{(\hat{I} \cdot \hat{S}[\hat{\sigma}])^{\hat{n}}}{(\hat{I} \cdot \hat{S}[\hat{\sigma}])^{\hat{n}} + (\hat{D}[\hat{A}_3])^{\hat{n}} + \hat{K}^{\hat{n}}}}d\hat{W}_1 = \frac{1}{\sqrt{\hat{v}}}\sqrt{\frac{\hat{v}_0}{\hat{v}} + \frac{(\frac{\hat{I}\cdot\hat{S}\cdot\hat{v}}{\hat{k}_{deg}\cdot\hat{K}}\frac{\hat{k}_{deg}[\hat{\sigma}]}{\hat{v}})^{\hat{n}}}{(\frac{\hat{I}\cdot\hat{S}\cdot\hat{v}}{\hat{k}_{deg}\cdot\hat{K}}\frac{\hat{k}_{deg}[\hat{\sigma}]}{\hat{v}})^{\hat{n}} + (\frac{\hat{D}\cdot\hat{v}}{\hat{k}_{deg}\cdot\hat{K}}\frac{\hat{k}_{deg}[\hat{A}_3]}{\hat{v}})^{\hat{n}} + 1}}d(\hat{k}_{deg}\hat{W}_1)$$

$$\frac{\hat{k}_{deg}}{\hat{v}}\sqrt{\hat{k}_{deg} \cdot [\hat{\sigma}]}d\hat{W}_2 = \frac{1}{\sqrt{\hat{v}}}\sqrt{\frac{\hat{k}_{deg}[\hat{\sigma}]}{\hat{v}}}d(\hat{k}_{deg}\hat{W}_2).$$

Finally, we consider the equations of the time-delay intermediaries:

$$\begin{cases} d[A_1] = dt(\frac{[\sigma]}{\tau} - \frac{[A_1]}{\tau}) + \sqrt{\frac{[\sigma]}{\tau}} \cdot dW_3 - \sqrt{\frac{[A_1]}{\tau}} \cdot dW_4 \\ d[A_2] = dt(\frac{[A_1]}{\tau} - \frac{[A_2]}{\tau}) + \sqrt{\frac{[A_1]}{\tau}} \cdot dW_5 - \sqrt{\frac{[A_2]}{\tau}} \cdot dW_6 \\ d[A_3] = dt(\frac{[A_2]}{\tau} - \frac{[A_3]}{\tau}) + \sqrt{\frac{[A_2]}{\tau}} \cdot dW_7 - \sqrt{\frac{[A_3]}{\tau}} \cdot dW_8 \end{cases}$$

$$\Longleftrightarrow$$

$$\begin{cases} d\frac{d\cdot[A_1]}{v} = d(d\cdot t)(\frac{d\cdot[\sigma]}{v}\frac{1}{d\tau} - \frac{d\cdot[A_1]}{v}\frac{1}{d\tau}) + \frac{1}{\sqrt{v}}\sqrt{\frac{d\cdot[\sigma]}{v}\frac{1}{d\tau}} \cdot d(d\cdot W_3) - \frac{1}{\sqrt{v}}\sqrt{\frac{d\cdot[A_1]}{v}\frac{1}{d\tau}} \cdot d(d\cdot W_4) \\ d\frac{d\cdot[A_2]}{v} = d(d\cdot t)(\frac{d\cdot[A_1]}{v}\frac{1}{d\tau} - \frac{d\cdot[A_2]}{v}\frac{1}{d\tau}) + \frac{1}{\sqrt{v}}\sqrt{\frac{d\cdot[A_1]}{v}\frac{1}{d\tau}} \cdot d(d\cdot W_5) - \frac{1}{\sqrt{v}}\sqrt{\frac{d\cdot[A_2]}{v}\frac{1}{d\tau}} \cdot d(d\cdot W_6) \\ d\frac{d\cdot[A_3]}{v} = d(d\cdot t)(\frac{d\cdot[A_2]}{v}\frac{1}{d\tau} - \frac{d\cdot[A_3]}{v}\frac{1}{d\tau}) + \frac{1}{\sqrt{v}}\sqrt{\frac{d\cdot[A_2]}{v}\frac{1}{d\tau}} \cdot d(d\cdot W_7) - \frac{1}{\sqrt{v}}\sqrt{\frac{d\cdot[A_3]}{v}\frac{1}{d\tau}} \cdot d(d\cdot W_8). \end{cases}$$

Now, we can perform the following substitutions:

$$\begin{cases} [\sigma] & = \frac{\hat{k}_{deg}\cdot[\hat{\sigma}]}{\hat{v}} \\ [A_1] & = \frac{\hat{k}_{deg}\cdot[\hat{A}_1]}{\hat{v}} \\ [A_2] & = \frac{\hat{k}_{deg}\cdot[\hat{A}_2]}{\hat{v}} \\ [A_3] & = \frac{\hat{k}_{deg}\cdot[\hat{A}_3]}{\hat{v}} \\ t & = \hat{k}_{deg}\hat{t} \\ W_i & = \hat{k}_{deg}\hat{W}_i \\ S & = \frac{\hat{S}\cdot\hat{i}\cdot\hat{v}}{\hat{k}_{deg}\cdot\hat{K}} \\ D & = \frac{\hat{D}\cdot\hat{v}}{\hat{k}_{deg}\cdot\hat{K}} \\ \tau & = \hat{k}_{deg}\hat{\tau} \\ v_0 & = \frac{\hat{v_0}}{\sqrt{\hat{v}}} \\ n & = \hat{n} \\ \eta & = \frac{1}{\sqrt{\hat{v}}} \end{cases}$$

which generate the following, nondimensionalised, set of SDEs:

$$\begin{cases} d[\sigma] & = dt\cdot(v_0 + \frac{(S[\sigma])^n}{(S[\sigma])^n+(D[A_3])^n+1} - [\sigma]) \\ & +\eta\sqrt{v_0 + \frac{(S[\sigma])^n}{(S[\sigma])^n+(D[A_3])^n+1}} \cdot dW_1 - \eta\sqrt{[\sigma]}\cdot dW_2 \\ d[A_1] & = dt(\frac{[\sigma]}{\tau} - \frac{[A_1]}{\tau}) \\ & +\eta\sqrt{\frac{[\sigma]}{\tau}}\cdot dW_3 - \eta\sqrt{\frac{[A_1]}{\tau}}\cdot dW_4 \\ d[A_2] & = dt(\frac{[A_1]}{\tau} - \frac{[A_2]}{\tau}) \\ & +\eta\sqrt{\frac{[A_1]}{\tau}}\cdot dW_5 - \eta\sqrt{\frac{[A_2]}{\tau}}\cdot dW_6 \\ d[A_3] & = dt(\frac{[A_2]}{\tau} - \frac{[A_3]}{\tau}) \\ & +\eta\sqrt{\frac{[A_2]}{\tau}}\cdot dW_7 - \eta\sqrt{\frac{[A_3]}{\tau}}\cdot dW_8. \end{cases}$$

A corresponding set of ODEs is also computed, and primarily used for steady state analysis:

$$\begin{cases} \frac{d[\sigma]}{dt} & = v_0 + \frac{(S[\sigma])^n}{(S[\sigma])^n+(D[A_3])^n+1} - [\sigma] \\ \frac{d[A_1]}{dt} & = \frac{[\sigma]}{\tau} - \frac{[A_1]}{\tau} \\ \frac{d[A_2]}{dt} & = \frac{[A_1]}{\tau} - \frac{[A_2]}{\tau} \\ \frac{d[A_3]}{dt} & = \frac{[A_2]}{\tau} - \frac{[A_3]}{\tau}. \end{cases}$$

The reactions and parameters of the model after nondimensionalisation are provided in the Methods section.

### 3 Replication of results using Gillespie's algorithm

To make the model simulations directly comparable to the stability analysis (which is carried out in continuous space), and to allow for the reduction in parameters using nondimensionalisation, we used the chemical Langevin equation (CLE) to implement noise [Gillespie, 2000]. Continuous space is also required for our definition of inactive and active states (Supplementary Section 6.2). However, *stochastic chemical kinetics* (which we here simulate using Gillespie's algorithm) typically yields more realistic trajectories [Gillespie, 1976, Gillespie, 1977]. Especially due to the general nature of our model, we consider the CLE a sound approach (checks also suggest that we are simulating the CLE in a noise regime where its approximations hold). However, to show that the results hold in discrete space, we used a non-nondimensionalised version of our model, simulated using the Gillespie algorithm, to recreate all of the response behaviours (Supplementary Figure 5). This demonstrates that the observed behaviours were not dependent on the CRN interpretation.

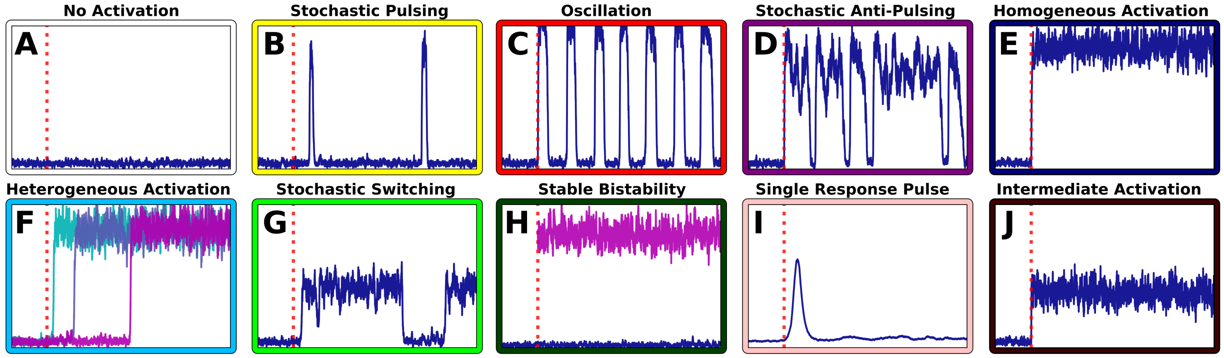

Supplementary Figure 5: **All the model behaviours can be recreated using Gillespie simulations.** Example simulations of the system's ten behaviours, but simulated using the Gillespie algorithm. The time scales in the various plots are different ($k_{deg}$ was used to modulate copy numbers, and thus stochasticity, which also affected the time scales). Simulations were carried out using the non-nondimensionalised model, as described in Supplementary Table 1.

Next, we wanted to show that our primary results (that the model's behaviours can be mapped across parameter space) could also be recreated using Gillespie simulations. Here, we constructed a rudimentary version of our classifier (Supplementary Section 6), but which was based on Gillespie simulations. Using it, we mapped the model's behaviours (Supplementary Figure 6). We note that this Gillespie simulation-based map reproduce similar behavioural patters as which was observed in the CLE-based maps (described in Supplementary Section 8).

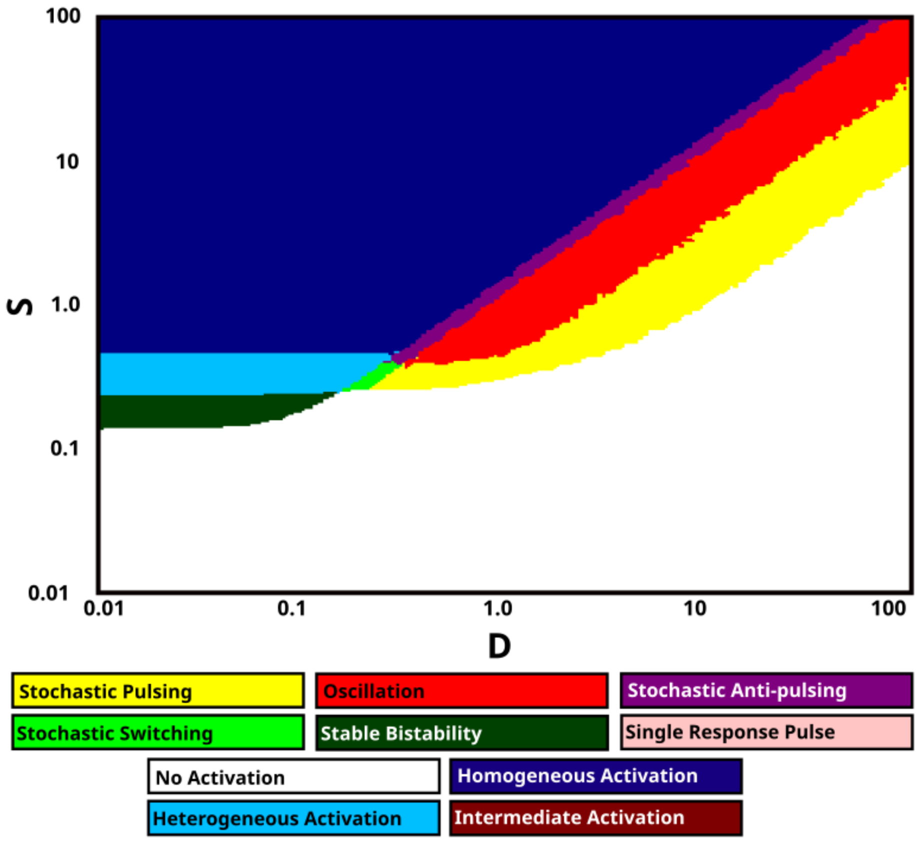

Supplementary Figure 6: **A Gillespie simulation-based classifier reproduces the behaviour maps generated by the CLE-based classifier.** Our Gillespie simulation-based classifier performs simulations based on the non-nodimensionalised model (Supplementary Section 2). This map shows the model behaviours in (logarithmic) $S, D$-space for the parameter set $(v_0, v, I, K, n, d, \tau) = (0.1, 2.0, 1.0, 15.0, 3, 0.01, 100.0)$. The map shows a similar behaviour distribution to which is generated by the CLE-based classifier (Supplementary Section 8).

### 4 Modelling the time delay

#### 4.1 Determining the number of intermediaries in the linear chain trick

The simplest implementation of a time-delay is a discrete delay, which generates a *delay differential equation* (DDE) (such an implementation is investigated in Supplementary Section 4.2). Here, the delay is defined by a single parameter only ($\tau$, the length of the delay). An alternative approach is to use a distributed delay, generating an *integro-differential equation* (IDE) [Cushing, 1977, Hu et al., 2018, Korsbo and Jönsson, 2020]. It has been shown that distributed delay may produce qualitatively different results from discrete delays, and these differences may have important biological implications [Beretta and Breda, 2016]. Distributed delays often depends on more than one parameter, e.g. the gamma distributed delay depends on a rate parameter $\tau$ (determining the length of the delay) and a shape parameter $n$. A gamma-distributed delay can be approximated using the *linear chain trick* (LCT), which replaces the delay term with a set of additional species that are produced and degraded at a linear rate [Fargue, 1973, MacDonald, 1978, Metz and Diekmann, 1991]. This has the advantage that an IDE can be replaced with an ODE, which are significantly easier to analyse and simulate (in our stochastic case, an SIDE is replaced by an SDE). While the LCT depend on only a single parameter ($\tau$, the rate), in its implementation, one also has to determine the number of intermediaries introduced (this number correspond to the shape parameter, $n$, of the corresponding gamma-distributed delay). We postulated that the shape would only have a minor effect on the behaviour, and that the complexity added to the model by making the number of intermediaries variable would not merit the introduction of an additional parameter. We thus decided to choose a fixed number of intermediaries.

According to [Heinrich et al., 2002], for two LCT implementations, with rate parameters $\tau_1$ and $\tau_2$, and shape parameters $N_1$ and $N_2$, the length of their delays is equivalent if

$$\frac{\tau_1}{N_1} = \frac{\tau_2}{N_2}.$$

We test this in practice by investigating the period of our general model for a range of values of $\tau$ and $N$ (Supplementary Figure 7). Our tests show that the estimate improves as $N$ becomes large. Especially, only for $N = 1$ or $N = 2$ is there a noticeable difference. Hence, we select $N = 3$ (the smallest value for which the estimate holds well). Finally, we also note that (in practical simulations) the $A_1$ and $A_3$ variables are highly correlated (with the latter being a delay of the former) (Supplementary Figure 8. This further suggests that the choice of the number of intermediaries has little effect on the model.

#### 4.2 Replication of result using a discrete delay

In addition to our main model, where we use a distributed delay implemented through the LCT, we also developed a model depending on a discrete delay. This model contain only two reactions (Supplementary Table 3). The parameters are identical to those of the distributed delay implementation. Finally, the RRE generates a single ODE:

$$\frac{d[\sigma]}{dt} = v_0 + \frac{(S[\sigma](t))^n}{(S[\sigma](t))^n + (D[\sigma](t-\tau))^n + 1} - [\sigma](t)$$

with a corresponding SDE generated by the CLE:

$$\begin{aligned} d[\sigma] = {} & dt \cdot (v_0 + \frac{(S[\sigma](t))^n}{(S[\sigma](t))^n + (D[\sigma](t-\tau))^n + 1} - [\sigma](t)) \\ & + \eta\sqrt{v_0 + \frac{(S[\sigma](t))^n}{(S[\sigma](t))^n + (D[\sigma](t-\tau))^n + 1}} \cdot dW_1 - \eta\sqrt{[\sigma](t)} \cdot dW_2. \end{aligned}$$

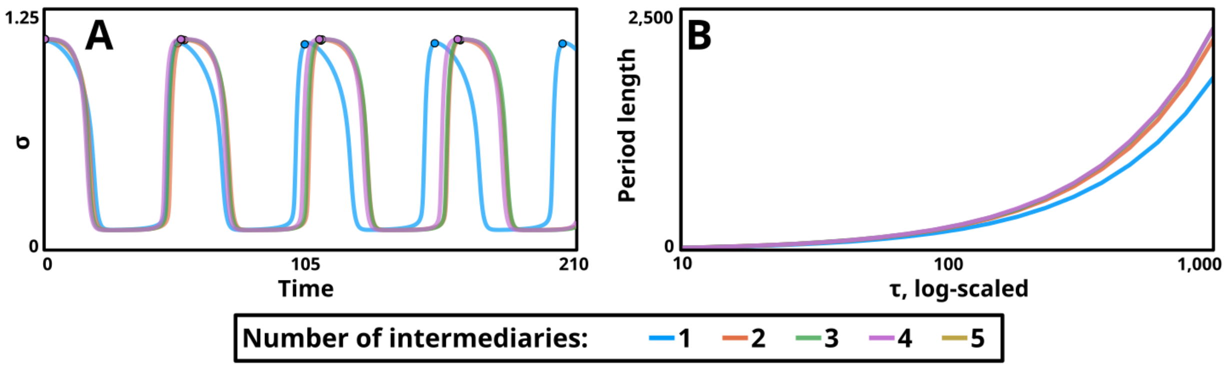

Supplementary Figure 7: **The number of intermediaries selected in the LCT has a small effect on the model's time-delay.** (A) Deterministic (using the reaction rate equation (RRE) interpretation) simulations of the model, implemented using various values of $N$ (intermediaries in the LCT). All parameters are identical, except for $\tau$, which is set according to the rule $\frac{\tau_1}{N_1} = \frac{\tau_2}{N_2}$, ensuring similar time-delay in all model implementations. As the oscillations persist, the simulations (especially the $N = 1$ and $N = 2$ ones) get desynchronised (B) For a range of values of $\tau$, the period length is measured for model implementations using $N = 1, 2, 3, 4, 5$. As $\tau$ grows large, the period lengths start to diverge. However, this difference is most distinct for $N = 1$ and $N = 2$. The previously mentioned rule is used to produce similar $\tau$ values for the different implementations, with the x-axis indicating the value of $\tau$ for $N = 1$.

| Description | Reaction | Rate | Propensity |
|---|---|---|---|
| Production | $\varnothing \longrightarrow \sigma$ | $v_0 + \frac{(S[\sigma](t))^n}{(S[\sigma](t))^n + (D[\sigma](t-\tau))^n + 1}$ | $v_0 + \frac{(S[\sigma](t))^n}{(S[\sigma](t))^n + (D[\sigma](t-\tau))^n + 1}$ |
| Degradation/dilution | $\sigma \longrightarrow \varnothing$ | 1 | $[\sigma](t)$ |

Supplementary Table 3: **The reactions of the discrete delay implementation of the general sigma factor model.** Here, $(t)$ is added at the end of species concentrations, to distinguish which terms are subject to a time-delay.

Finding steady states (and there stability) is much harder for DDEs than ODEs. In addition, there are much fewer algorithms available for simulations of DDEs and SDDEs (as compared to ODEs and SDEs). Hence, we will limit our analysis of this model to demonstrating its ability to generate the ten different response behaviour observed in the distributed delay implementation of our model (Supplementary Figure 9)

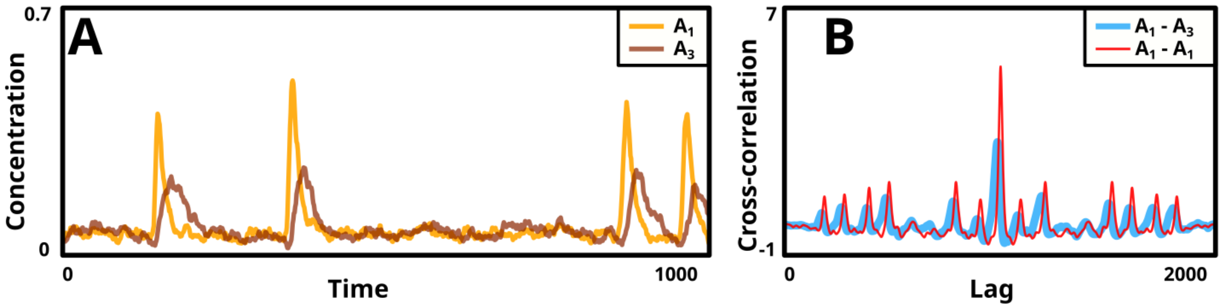

Supplementary Figure 8: **The $A_1$ and $A_3$ components can be compared using cross-correlation.** (A) The concentration of the $A_1$ (yellow) and $A_3$ (brown) components throughout a single (stochastic pulsing) simulation (input active throughout the simulation). The two trajectories are closely correlated. (B) The cross-correlation between $A_1$ and $A_3$ (blue) and between $A_1$ and itself (red). The distinct peak in the middle of the diagram indicates that the two curves are closely correlated (indeed, the $A_1$ to $A_3$ correlation is almost as good as $A_1$ to itself). The slight offset between the two curves along the x-axis indicates the length of the delay between the two components.

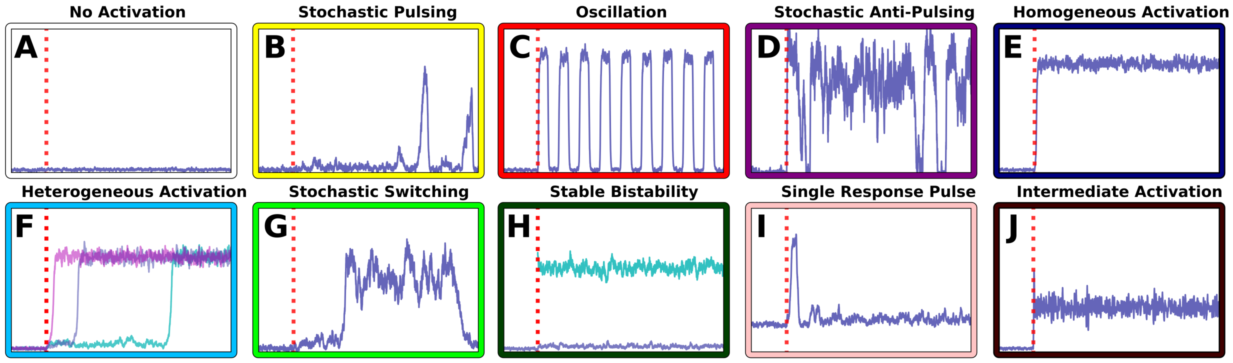

Supplementary Figure 9: **All the model behaviours can be recreated using the discrete delay implementation of the model.** Example simulations of the system's ten behaviours, but simulated using the discrete delay model.

### 5 Steady state analysis

#### 5.1 Steady state computations

There exists a range of approaches for finding the steady states of a system of ODEs. Here, we will use algebraic manipulation to find a single polynomial whose roots corresponds to the steady states of our system. Polynomial roots, and hence our model's steady states, can be computed reliably.

We start with the model's ODE system (generated through the reaction rate equations) and set the derivatives equal to 0 (to find the steady states):

$$\begin{cases} 0 &= v_0 + \frac{(S[\sigma])^n}{(S[\sigma])^n + (D[A_3])^n + 1} - [\sigma] \\ 0 &= \frac{[\sigma]}{\tau} - \frac{[A_1]}{\tau} \\ 0 &= \frac{[A_1]}{\tau} - \frac{[A_2]}{\tau} \\ 0 &= \frac{[A_2]}{\tau} - \frac{[A_3]}{\tau}. \end{cases}$$

here, the last three equations trivially yields $[A_i] = [\sigma]$, we thus only needs to solve:

$$0 = v_0 + \frac{(S[\sigma])^n}{(S[\sigma])^n + (D[\sigma])^n + 1} - [\sigma]$$

which we can rewrite as:

$$(v_0 - [\sigma])((S[\sigma])^n + (D[\sigma])^n + 1) = (S[\sigma])^n$$

$$\Longleftrightarrow$$

$$(S^n + D^n)[\sigma]^{n+1} - (S^n + v_0 \cdot S^N + v_0 \cdot D^n)[\sigma]^n + [\sigma] - v_0 = 0$$

which gives us our desired polynomial. Furthermore, according to Descartes' rule of signs, this polynomial has at most three real roots, setting a limit for the maximum number of steady states of our system. Since all three steady states cannot be stable we are left with the alternatives of either zero, one, or two stable steady states.

#### 5.2 Nullcline analysis

Nullclines (curves along which the rate of change, with respect to a specified system variable, is 0) are commonly used during phase space analysis. In this section, we will first compute our system's nullclines, and then investigate how there properties depend on the system's parameters. The nullclines will also be shown in Supplementary Section 7 and they are important for our classification of the system's behaviour (Suplementary Section 6).

The nullclines can be computed by solving the equations $[\dot{\sigma}] = 0$ and $[\dot{A}_i] = 0$ (generating the curves $NC_\sigma([\sigma])$ and $NC_{A_i}([\sigma])$, respectively). Let's briefly consider the LCT with $N$ intermediaries:

$$\begin{cases} 0 &= v_0 + \frac{(S[\sigma])^n}{(S[\sigma])^n + (D[A_N])^n + 1} - [\sigma] \\ 0 &= \frac{[\sigma]}{\tau} - \frac{[A_i]}{\tau} \end{cases}$$

The second set of nullclines are, trivially,

$$[A_i] = [\sigma].$$

The first one can be derived as:

$$([\sigma] - v_0)((S[\sigma])^n + (D[A_N])^n + 1) = (S[\sigma])^n$$

$$\Longleftrightarrow$$

$$(D[A_N])^n = \frac{(S[\sigma])^n}{[\sigma] - v_0} - (S[\sigma])^n - 1$$

$$\Longleftrightarrow$$

$$[A_N] = \frac{1}{D}\sqrt[n]{(S[\sigma])^n(\frac{1}{[\sigma] - v_0} - 1) - 1}$$

We note that the root of $NC_\sigma([\sigma])$ is not defined for negative arguments. While the initial equations might have had a solution for these values, they would result in $[A_N] < 0$. Since the phase space of a CRN is limited to non-negative species concentrations, these values of the nullcline are not of interest. We can safely do the root, and ignore the nullcline where the root is undefined.

We will use three intermediaries in our implementation of the LCT, however, for phase space analysis we will display the system in $[\sigma]$-$[A_3]$ space (ignoring the $[A_1]$ and $[A_2]$ dimensions). The two relevant nullclines are:

$$[A_3] = NC_\sigma([\sigma]) = \frac{1}{D}\sqrt[n]{(S[\sigma])^n(\frac{1}{[\sigma] - v_0} - 1) - 1}$$

$$[A_3] = NC_{A_3}([\sigma]) = [\sigma].$$

These intersect either once or trice (Supplementary Section 5.1), producing a system with either one or three steady states (Supplementary Figure 10).

While the curve $NC_A([\sigma])$ is unaffected by the parameters ($S$, $D$, $\tau$, $v_0$, $n$, and $\eta$), their values do affect the curve $NC_\sigma$. Changing $S$ (the degree of system self-activation) changes its shape. For large or small values of $S$, $NC_\sigma$ is a descending line, permitting only a single steady state. Only for intermediate values of $S$ is bistability possible (Supplementary Figure 10B). For $S = 0$ (corresponding to a system with no input), $NC_\sigma$ is a straight line at $[\sigma] = v_0$ (yielding a single steady state at $([\sigma], [A_i]) = (v_0, v_0)$ for the system before input). Changing $D$ will only scale $NC_\sigma$, but not affect its shape further (Supplementary Figure 10C). For some critical values of $S$ (where the system is close to, or in, the bistability region) $v_0$ may affect the shape of $NC_\sigma$. For most values of $S$, however, modulating $v_0$ has little effect on $NC_\sigma$ (Supplementary Figure 10D). Increasing $n$ increases the sharpness of the curve (Supplementary Figure 10E). Finally, $\eta$ and $\tau$ do not affect $NC_\sigma$, however, $\tau$ may affect stability properties.

#### 5.3 Bifurcation analysis

Additional information of our system can be inferred by considering its bifurcation diagrams. These describe how the system's steady states depends on its parameters. Here, we will only analyse the bifurcation diagrams with respect to the parameter $S$. This parameter describes the degree of input (or stress) to the system (as well as its degree of self-activation), and the bifurcation diagrams thus show how the steady states of the system are affected by the input's magnitude. We note two types of bifurcation diagrams (which one is observed mainly depends on the value of $D$, Supplementary Figure 11G). One displays bistability for intermediate values of $S$ (this is known as a bistable switch, Supplementary Figure 11A,D)). The other has a single steady state for all values of $S$ (Supplementary Figure 11C,F). For some parameter values, the bistable diagram may instead be monostable, with the two larger valued steady states being unstable (Supplementary Figure 11B,E). The value of $v_0$ has little effect on the single steady state diagrams, but affects the shape of the bistable switch (Supplementary Figure 11H). Next, $n$ also affects the shape of the bistable switch (which requires $n > 1$, Supplementary Figure 11I). Finally, the steady state values are unaffected by the value of $\tau$, but for large time-delays the single steady state diagrams develop an increasing region of instability (corresponding to an oscillation) (Supplementary Figure 11J). All bifurcation diagrams are computed using the BifurcationKit.jl Julia package [Veltz, 2020].

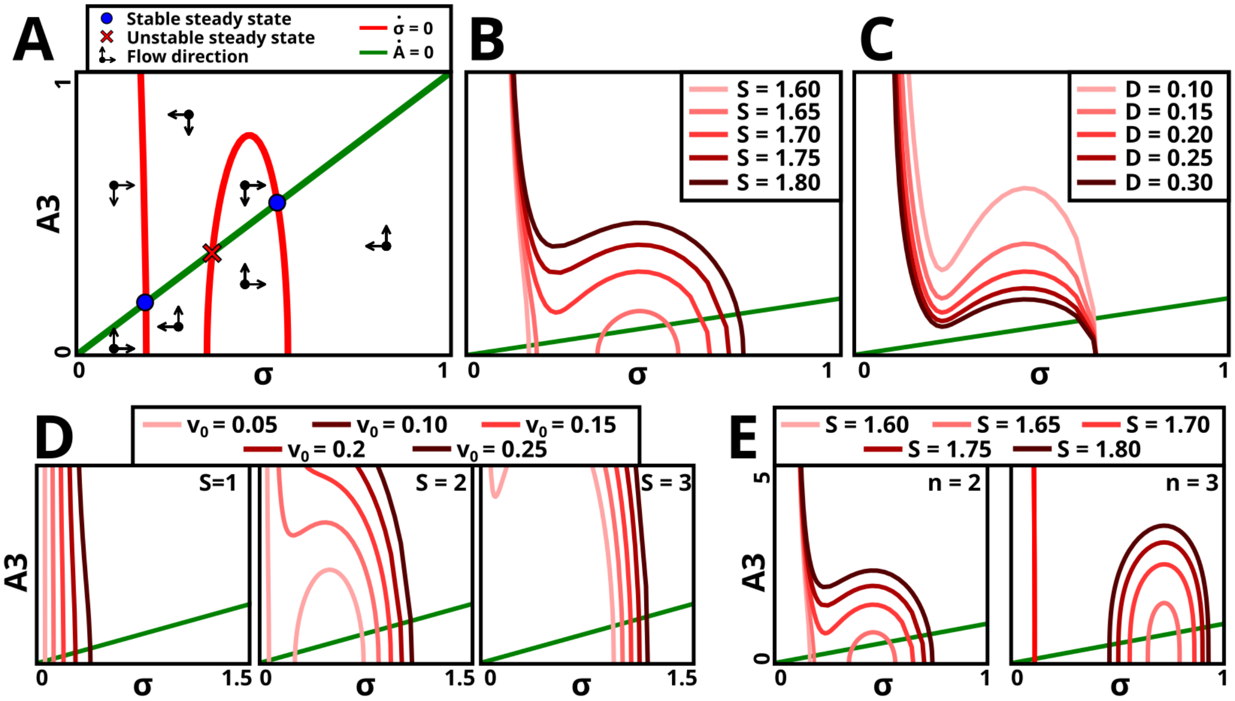

Supplementary Figure 10: **The model's nullclines plotted in phase space.** The two nullclines $NC_\sigma$ (shades of red) and the $NC_A$ (green). (A) The steady states of the system can be found as the nullclines' intersections. Also, in each region of the phase space, as separated by the nullclines, the direction of flow of the system can be determined, this is marked by the arrows. (B) The parameter $S$ affects the shape of $NC_\sigma$. For low and high $S$ there is only one steady state (with small and large $[\sigma]$, respectively). Only for intermediate values of $S$ is the system bistable. (C) Modulating the parameter $D$ only scales $NC_\sigma$, but does not affect the shape. (D) For small or large values of $S$, changing $v_0$ only has a minor effect on $NC_\sigma$. However, for intermediate values of $S$, modulating $v_0$ changes the shape. (E) Increasing $n$ makes the $NC_\sigma$ sharper, emphasising bistability. The parameters $\tau$ and $\eta$ do not affect $NC_\sigma$. Finally, we note that our analysis here is primarily carried out as if our 4-variable system (with variables $\sigma(t)$, $A_1(t)$, $A_2(t)$ and $A_3(t)$) is a 2-variable system (with variables $\sigma(t)$ and $A_3(t)$). However, the dynamics of the 4-variables is *very* similar to the 2-variable one with only a single intermediate ($A$) (Supplementary Figure 8), and that these two systems have identical nullclines.

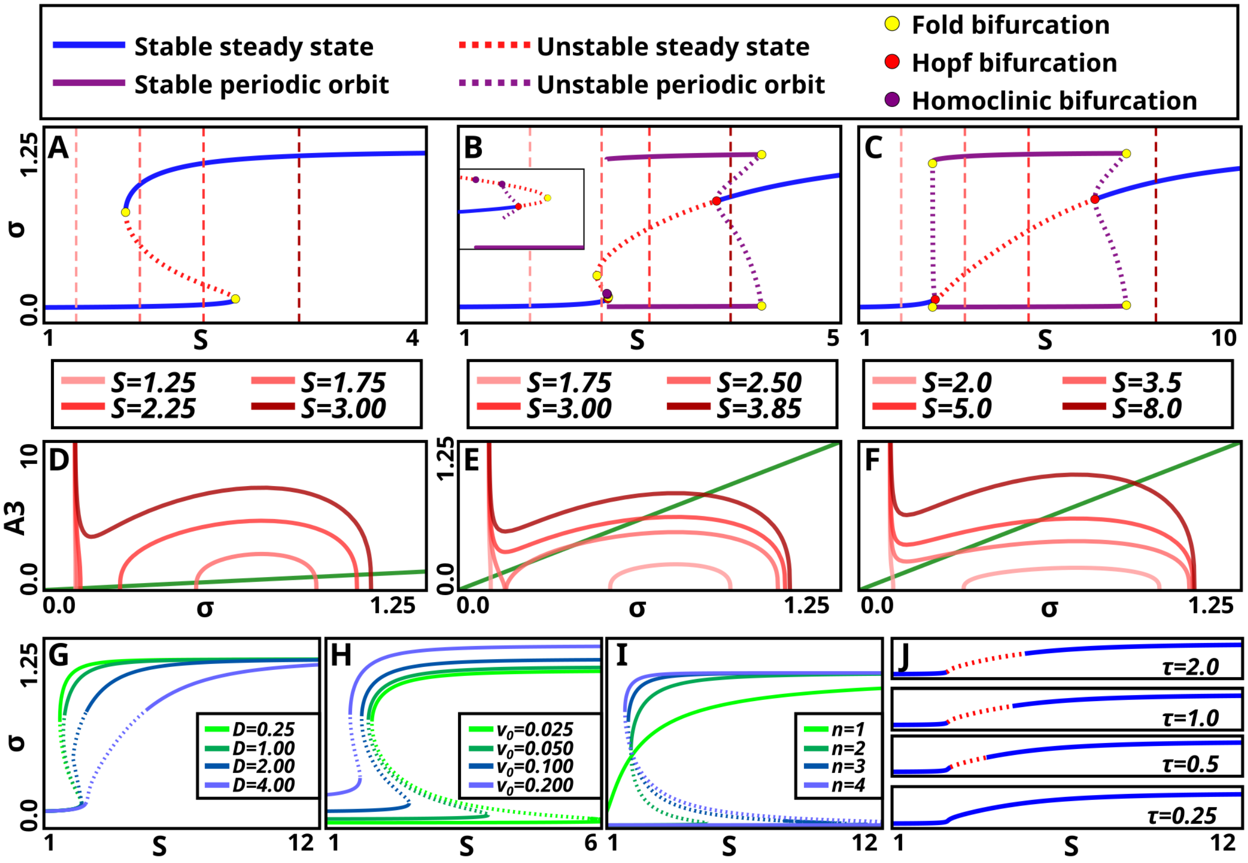

Supplementary Figure 11: **The model's bifurcation diagrams with respect to the parameter *S*.** (A-F) For three different model instances, the system's bifurcation diagram is plotted (top), and the nullclines of the same system plotted in phase space (bottom). The $[\dot{\sigma}] = 0$ nullcline is plotted for four different values of $S$ (these values are also marked with dotted lines in the bifurcation diagram). (A,D) The system where it exhibits a bistable switch. (B,E) The system may also exhibit a region with two unstable steady states, and a single stable one. (C,F) Finally, an instance that does not exhibit bistability, and smoothly transition from a single inactive state to a single active state. (G-I) Bifurcation diagrams plotted for a selection of parameter values. Solid lines indicate stable steady states and dotted unstable ones. (G) As the parameter $D$ is modulated, it exhibits the range of bifurcation diagrams in A-C. (H) Modulating the parameter $v_0$ affects the size of the bistability region. (I) Bifurcation diagrams for a few values of $n$, only for $n > 1$ does the system exhibit bistability. (J) As the parameter $\tau$ is increased, the size of mono-unstable region increase.

### 6 Behaviour classification

#### 6.1 Determining the model's possible response behaviours

To examine our model we determined the behaviours that it might exhibit. The behaviours were classified according to the system's possible transitions between its steady states. The system either contains a single unstable steady state (and thus oscillates), or one or two stable steady states (as shown in Supplementary Section 5.1). If the system is bistable, one (stable) steady state corresponds to an *inactive value*, and one to an *active value*. If it is monostable, the steady state corresponds to either an *inactive value*, an *active value*, or some *intermediate value*.

Given a system with at least a single steady state, the solution will asymptotically tend toward a steady state. However, fluctuations may push the system out of a steady state (either transitioning to another steady state, or, in an excitable behaviour, into a temporary pulse which eventually return to the initial steady state). Two types of fluctuations may push the system out of a steady state. One type is stochastic noise. Alternatively, at the onset of the input ($S$ changed from 0 to its designated value), the steady states of the system shift, causing the system to exist in a non-steady state state. This constitutes a perturbation to the system. The remaining paragraphs of this section will describe how we classify a model instance according to the types of transitions between states it exhibits.

In absence of the input ($S = 0$), the system has a single steady state at $([\sigma], [A_i]) = (v_0, v_0)$ (where it rests). At the input's onset, the system may shift into an active state (briefly ignoring the possibility of an intermediate state, which is handled separately at the end). This transition is driven by fluctuations both due to the input's onset, as well as noise. This time of activation we call the *initial activation time* (IAT). If the system reaches an active state, it may similarly transition back into an inactive state, this time we call the *deactivation time* (DT). Finally, if the system deactivates, it may again activate. The time to reactivate after having deactivated we call the *reactivation time* (RT, which is different from the initial activation time in that it is only driven by noise). Deactivation times that follow either a normal activation or an initial activation should be identically distributed (hence we do not need to split these into initial and normal deactivation times). Here, the initial activation is a transient behaviour, while deactivations and reactivations are asymptotic behaviours.

We classify the times for this state shift as either *instantaneous* (happens immediately upon input onset, activation, or deactivation), *heterogeneous* (the transition happens after some delay, and not necessarily immediately), and *infinite* (the transition never happens). The three types of transitions, to the power of the three classes of transitions, generate 27 different alternative patterns of transitions. However, several will not happen (e.g. a system with infinite activation time will not have a reactivation time). Also, some combinations will yield behaviours that we assign to the same class. The full classification can be found in Supplementary Figure 12.

This scheme does not work on a system with a single steady state of intermediate activity. For what remains of this section, we will describe how we classify these systems. If this state is unstable, we will classify it as an oscillation (using the observation that a well-behaved CRN with only unstable steady states adopts a limit cycle). If the steady state is stable, we will classify the system as an *intermediate activation* (where the system, at the input's onset, immediately and homogeneously, transitions into a stable state of intermediate activity). One could imagine separate behaviours using intermediate states (e.g. stochastic pulsing from an intermediate state). Most of these, however, would have little qualitative difference from behaviours already defined in Supplementary Figure 12 (e.g. stochastic switching between an inactive state and an intermediate state). We rigorously scanned for new behaviours depending on steady states of intermediate activity, and only found three potential such cases:

- A limited *heterogeneous intermediate activation*, where the system, at the onset of the input, heterogeneously transitioned to a stable steady state of intermediate activity. However, the activation times were short and bounded (and did not seem to depend on the system remaining in a quasi-stable steady state, as is the case for normal heterogeneous activation). The region in parameter space where this

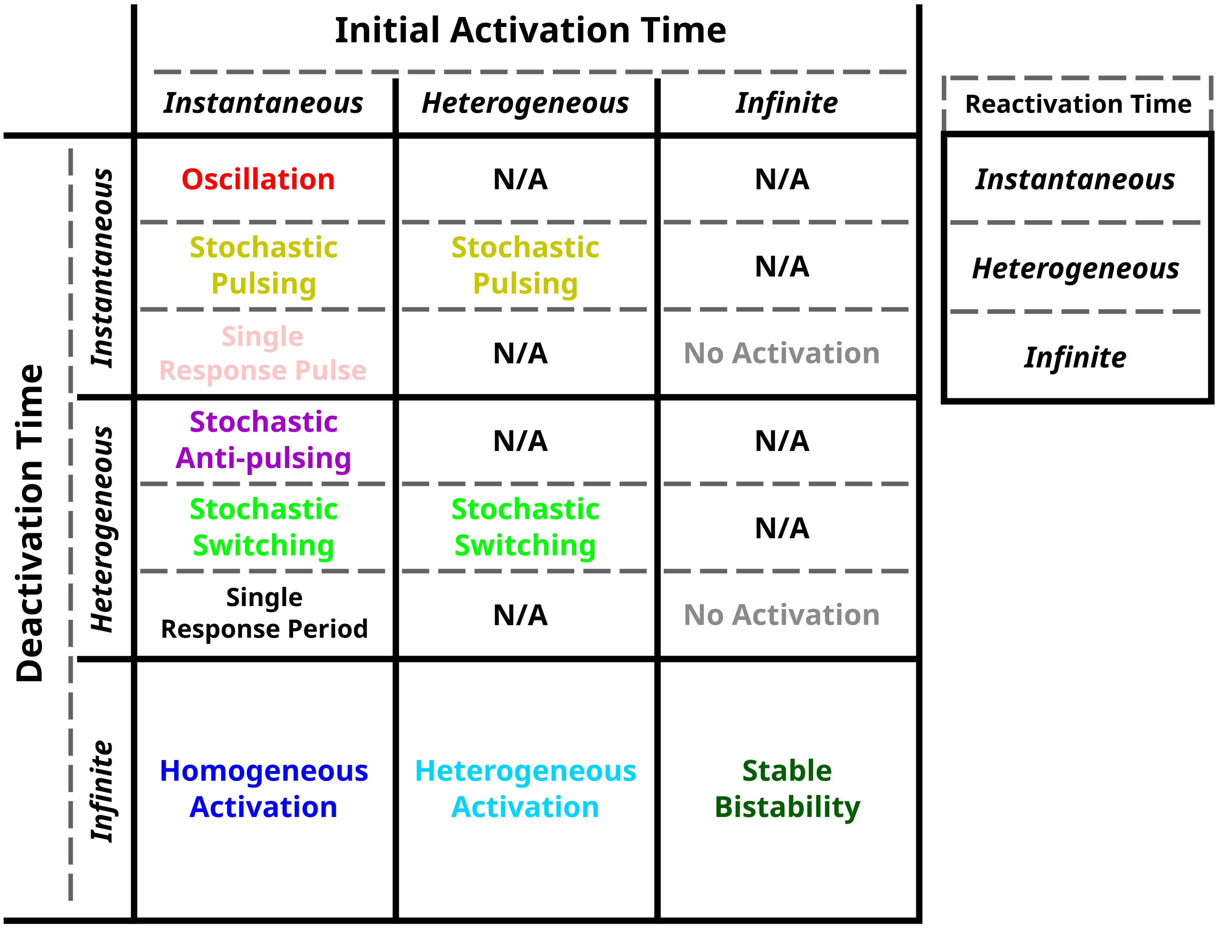

Supplementary Figure 12: **The model's potential behaviours are classified through its activation and deactivation times.** We define the model's *initial activation time* (IAT, time from input onset to it first reaching an active state), its *deactivation time* (DT, time from it reaching an active state to it reaching an inactive state) and its *reactivation time* (RT, time from it reaching an inactive state to it reaching an active state). Each set of times can either be classified as *instantaneous* (the transition is immediate), *heterogeneous* (the transition happens, but not necessarily instantaneously, over an ensemble the times are heterogeneous), and *infinite* (the transition never happens). The behaviours are located in a 3x3x3 grid, where the first dimension corresponds to the IAT type, the second to the DT type, and the third to the RT type. In the figure, the first two dimensions (IAT and DT) are shown in a 3x3 grid, with each field containing three subfields (corresponding to the third dimension, RT). The text in each subfield indicates what behaviour that combinations generate. A system with infinite DT cannot activate, hence no separate RT fields are given in these cases. Furthermore, if the IAT is infinite, we assume the RT is also infinite (the IAT is produced by perturbation due to input onset and noise, while the RT onset is only due to noise, if the formed cannot yield a transition neither should the later). Using a similar argument, we assume that if the IAT is heterogeneous so is the RT. This produces a couple of impossible combinations (e.g. IAT infinite, DT instantaneous, and RT heterogeneous), which are marked as N/A. Next, the stochastic switching and stochastic pulsing behaviours can be subdivided depending on whether the IAT is instantaneous or heterogeneous, however, both these cases are assigned the same class (hence these two classifications each occur twice in the grid). Finally, the IAT instantaneous, DT heterogeneous, and RT infinite combination would yield a behaviour we call *single response period*. However, we have never encountered this behaviour, which is purely theoretical.

behaviour occurred was also small, and adjacent to those of normal intermediate activation. Hence this sub-behaviour was classified as an intermediate activation.

- In some cases, the system exhibited a response pulse, followed by relaxation into an intermediate state. However, due to the higher levels of $[\sigma]$ in the steady state, the response pulse would be less distinct. This was classified as an intermediate activation.

- For some bistable systems, the active state was narrowly in the region of intermediate activation. Otherwise, the system behaved like a bistable system. Again, this happened in a very narrow parameter region (adjacent to that where normal bistability occurred). In these cases, the system was considered as a bistable system with an inactive and an active stable steady state.

We recognise that more granularity could be added to the scheme, further subdividing the classes. This could include re-defining stochastic pulsing depending on whether there is an initial pulse in response to the input's onset (or if they only occur asymptotically). However, we decided further subdivisions would not be beneficial. In Supplementary Section 6.4, we list potential subdivisions of the 10 behaviours defined in the section.

#### 6.2 Defining active and inactive states in the model

In Supplementary Section 6.3, we use the local minimum and maximum of the nullclines to define whether a system is in an active or in an inactive state (Supplementary Figure 13). These can be found as the values of $[\sigma]$ where $NC_\sigma([\sigma])'_{[\sigma]} = 0$. Again, while they cannot be found analytically, we can compute a polynomial with roots that both can reliably be found numerically and which corresponds to the nullcline's local minimum and maximum. First, we note that:

$$\frac{d}{dx} p \cdot f(x)^{1/n} = \frac{p}{n} f(x)^{1/n-1} \cdot f'(x)$$

since we are only interested in values where $NC_\sigma([\sigma]) \geq 0$ we only need to consider the inside function. We thus solve

$$\frac{d}{d[\sigma]}[(S[\sigma])^n(\frac{1}{[\sigma]-v_0} - 1) - 1] = 0$$

$$\iff$$

$$n \cdot S^n \cdot [\sigma]^{n-1}(\frac{1}{[\sigma]-v_0} - 1) - S^n[\sigma]^n(\frac{1}{([\sigma]-v_0)^2}) = 0$$

$$\iff$$

$$n(\frac{1}{[\sigma]-v_0} - 1) - [\sigma](\frac{1}{([\sigma]-v_0)^2}) = 0$$

$$\iff$$

$$-\frac{[\sigma]}{n} + [\sigma] - v_0 - ([\sigma]-v_0)^s = 0$$

$$\iff$$

$$[\sigma]^2 + (2v_0 + 1 - \frac{1}{n})[\sigma] + v_0^2 + v_0 = 0$$

which is our desired polynomial. Its roots can be computed as:

$$[\sigma]_0^\pm = \frac{1}{2n} - \frac{1}{2} - v_0 \pm \sqrt{(\frac{1}{2n} = \frac{1}{2} - v_0)^2 - v_0^2 - v_0}$$

$$\iff$$

$$[\sigma]_0^\pm = \frac{1}{2n} - \frac{1}{2} - v_0 \pm \sqrt{\frac{1}{4n^2} + \frac{1}{4} - \frac{1}{2n} - \frac{v_0}{n}}$$

which either generates two roots for positive $[\sigma]$, or, when $v_0$ exceeds a critical value $v_0^*$, complex roots (that is, no local minimum and maximum exists).

As $v_0 \longrightarrow v_0^*$ the two roots approach each other. As they do, the region where the system has an intermediate state diminishes (this region will be defined as $[\sigma]_0^- < [\sigma] < [\sigma]_0^+$). For $v_0 > v_0^*$ we will still want to define active and inactive states, but will not be able to do so through the nullclines local minimum and maximum (since these no longer exist). For $v_0 > v_0^*$, we define the activity threshold as the critical value of the two roots as they absorb each other at $v_0^*$ ($[\sigma]_0^* = [\sigma]_0^+(v_0^*) = [\sigma]_0^-(v_0^*)$. First, we note that the $v_0^*$ can be found as a function of $n$ only, by solving:

$$\frac{1}{4n^2} + \frac{1}{4} - \frac{1}{2n} - \frac{v_0}{n} = 0$$

which gives:

$$v_0^* = \frac{n}{4} - \frac{1}{2} + \frac{1}{4n}$$

and finally, we find $[\sigma]_0^*$ as the value of the roots where the square root is 0:

$$[\sigma]_0^* = \frac{1}{2n} - \frac{1}{2} - v_0$$

When classifying the behaviours of our model, we will need to define whether a state is to be considered *active*, *inactive*, or *intermediately active*. To do so, we use a scheme based on the local and minimum and maximum of the system's nullcline (more specifically, the $[\dot{\sigma}] = 0$ nullcline, also called $NC_\sigma$). Here, states with a value of $[\sigma] < T_{ia}$ (with $T_{ia}$ being the local minimum of the nullcline) are inactive. States with a value of $[\sigma] > T_a$ (with $T_a$ being the local maximum of the nullcline) are active. Remaining states (with $T_{ia} \geq [\sigma] \leq T_a$) are intermediately active. The scheme is described in detail in Supplementary Figure 13.

#### 6.3 Automated classification of the model's response behaviour

To enable us to classify the behaviour of the system at a given parameter set, we designed a classification algorithm. An initial algorithm was based on finding the steady states of the RRE interpretation of the system. While this algorithm yielded results similar to the final one, due to noise's tendency to shift a system's fixed points the algorithm had to be modified [Rao et al., 2002]. Over the next few paragraphs, as well as in Supplementary Figure 14, we present a heuristic algorithm that was created by modifying the initial one. Since the efficiency of the algorithm is not necessarily confirmed by its description, we demonstrate its performance through showing trial classifications. We generated a large number of parameter sets, and classified them. That the classifier generates appropriate classifications for this random sample can be seen in Supplementary Figures 15 and 16.

In the first step of our heuristic algorithm, the steady states of the parameter set are classified. First, the steady states of the deterministic RRE interpretation of the model are determined. If the system is bistable it is classified as "bistable". Else, the median $\sigma$ concentration of the system (measured over several long simulations, $11,000$ time units) is found. The system is then classified as "inactive", "active", or "intermediate" (corresponding according to what property the median value has according to the scheme in Supplementary Figure 13).

Next, the steady state(s) are computed (as the deterministic steady states in the bistable case, by using the median values of each component of the system in the single steady state cases). From these, the system's reactivation time (RT, inactive and bistable systems only) and deactivation time (DT, active and bistable systems only) are computed. This is done by starting a number of simulations ($N = 40$) in the inactive (for RT) or active (for DT) and measuring the mean times it takes the system to pass the threshold ($T_a$ for RT and $T_{ia}$ for DT). If the mean passage time is lower than an (arbitrarily selected) threshold (of

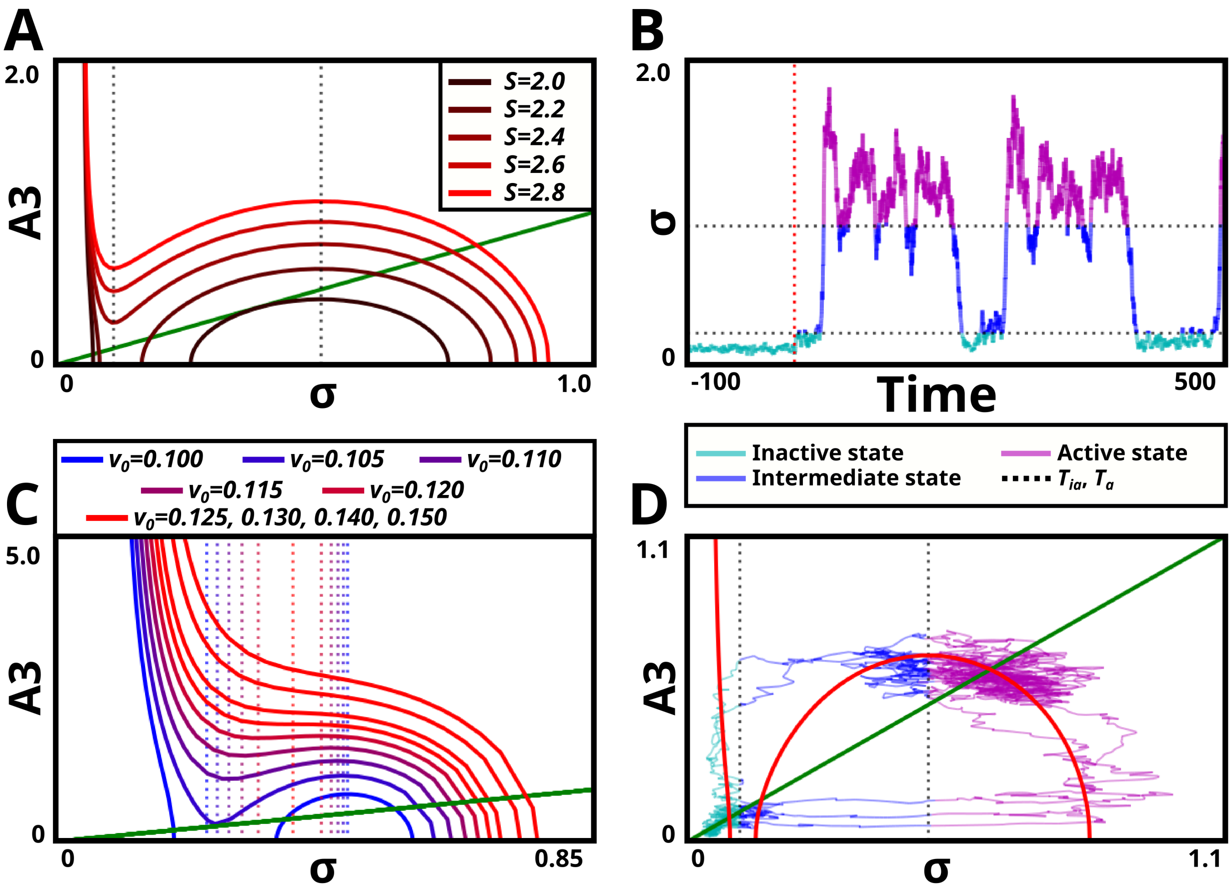

Supplementary Figure 13: **We use the nullclines to define what states of the model are active and which are inactive.** (A) The nullcline $NC_\sigma$ (for $[\dot{\sigma}] = 0$) may have a local minimum and maximum (their $[\sigma]$ values are called $T_{ia}$ and $T_a$). $T_{ia}$ and $T_a$ are independent of the values of $S$ (and $D$, $\tau$, and $\eta$). (B,D) A single simulation of the model, with $T_{ia}$ and $T_a$ marked in black dotted line. The trajectories are coloured according to whether the current state is inactive (teal), intermediate active (blue), or active (purple). (B) The simulation shown in time. (D) The simulation shown in phase space (the nullclines plotted in red and green). (C) As the value of $v_0$ increases, the thresholds $T_{ia}$ and $T_a$ approach each other. Here, for a range of values of $v_0$, the nullcline (solid line) and the corresponding thresholds (dotted lines) are plotted. For large values of $v_0$ ($v_0 > v_0^*$), neither local minimum nor maximum exists (bright red lines). Here, both $T_{ia}$ and $T_a$ are set to the boundary threshold obtained at the annihilation point.

25 time units), they are classified as *instantaneous*. If a certain percentage of the simulations (arbitrarily selected value of 85%) had not passed the threshold at the end of the simulation (length 2,000 time units), the times are classified as *infinite*. Else, they are classified as *heterogeneous*. A fourth case, with passage times being very similar, but with a mean higher than the instantaneous activation threshold can be considered (*delayed homogeneous passage*). However, this happened so rarely that its occurrence is most likely due to random chance. Hence we do not include this case.

Finally, for some cases, the IAT were measured and classified. These were measured by starting a number of simulations ($N = 40$) in the state corresponding to system with no input ($(\sigma, A_i) = (v_0, v_0)$) and measuring the mean time for the simulations to pass the threshold to an active state ($T_a$). If the mean was lower than a threshold (25 time units) the IAT were classified as instantaneous. Else they were classified as *heterogeneous or infinite*. In this case, the additional effect of the transient effect due to the change in parameter values would have worn off, and the passage times should be asymptotically identical to the

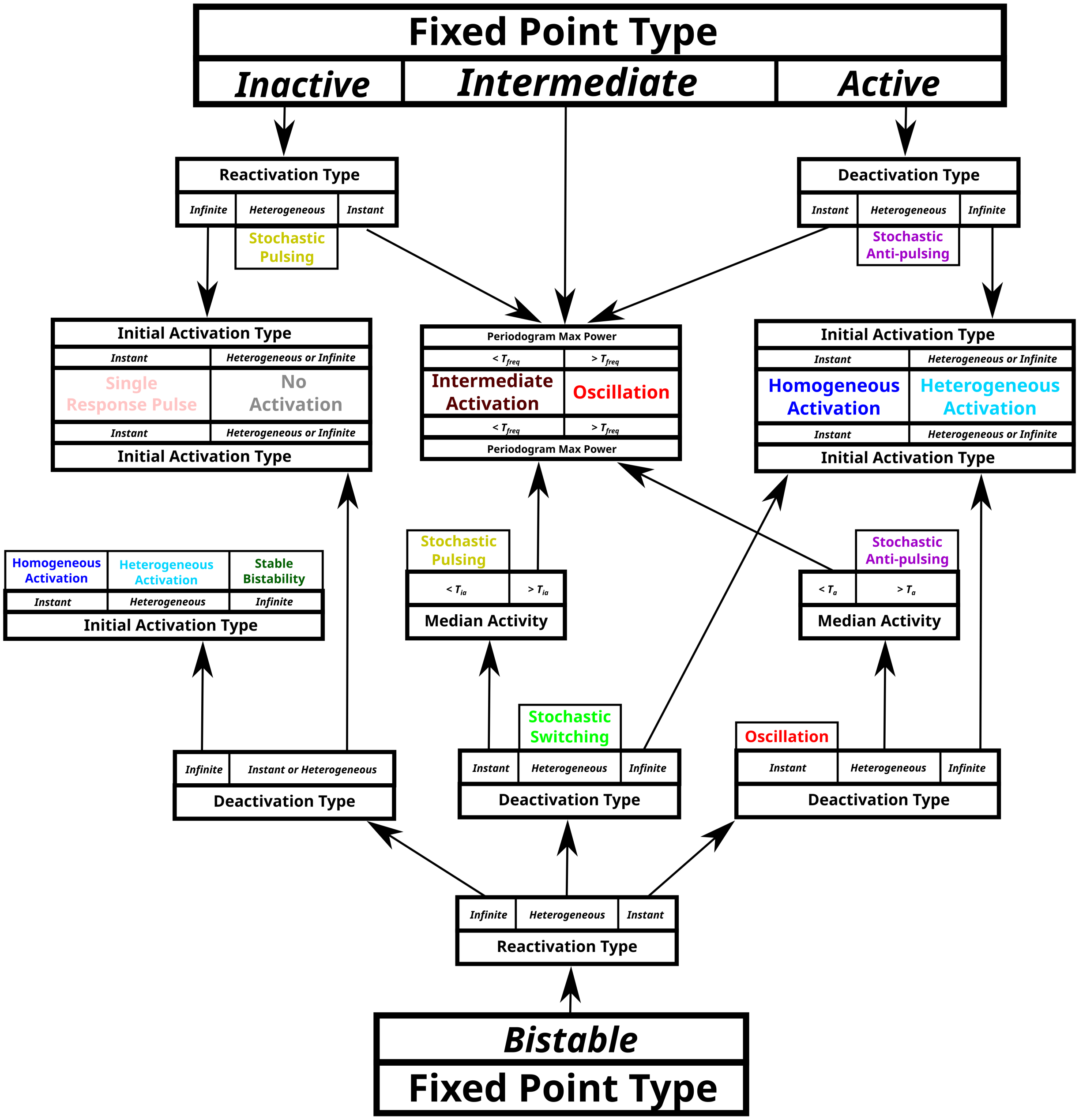

Supplementary Figure 14: **A decision tree heuristic is used to classify the response behaviour of a model instance.** The system's fixed point type (inactive, intermediate, active, or bistable) is first measured, then in succession its: reactivation type, deactivation type, and initial activation type (each either classified as infinite, heterogeneous, or instantaneous). Boxes with coloured texts indicate a classification result. In some cases, it is checked whether the system's median activity surpasses some threshold ($T_{ia}$ being the nullcline's local minimum and $T_a$ its local maximum, Supplementary Section 6.2). Similarly, it is checked whether the maximum value of the system's Welch type periodogram exceeds some threshold value ($T_{freq} = 10$).

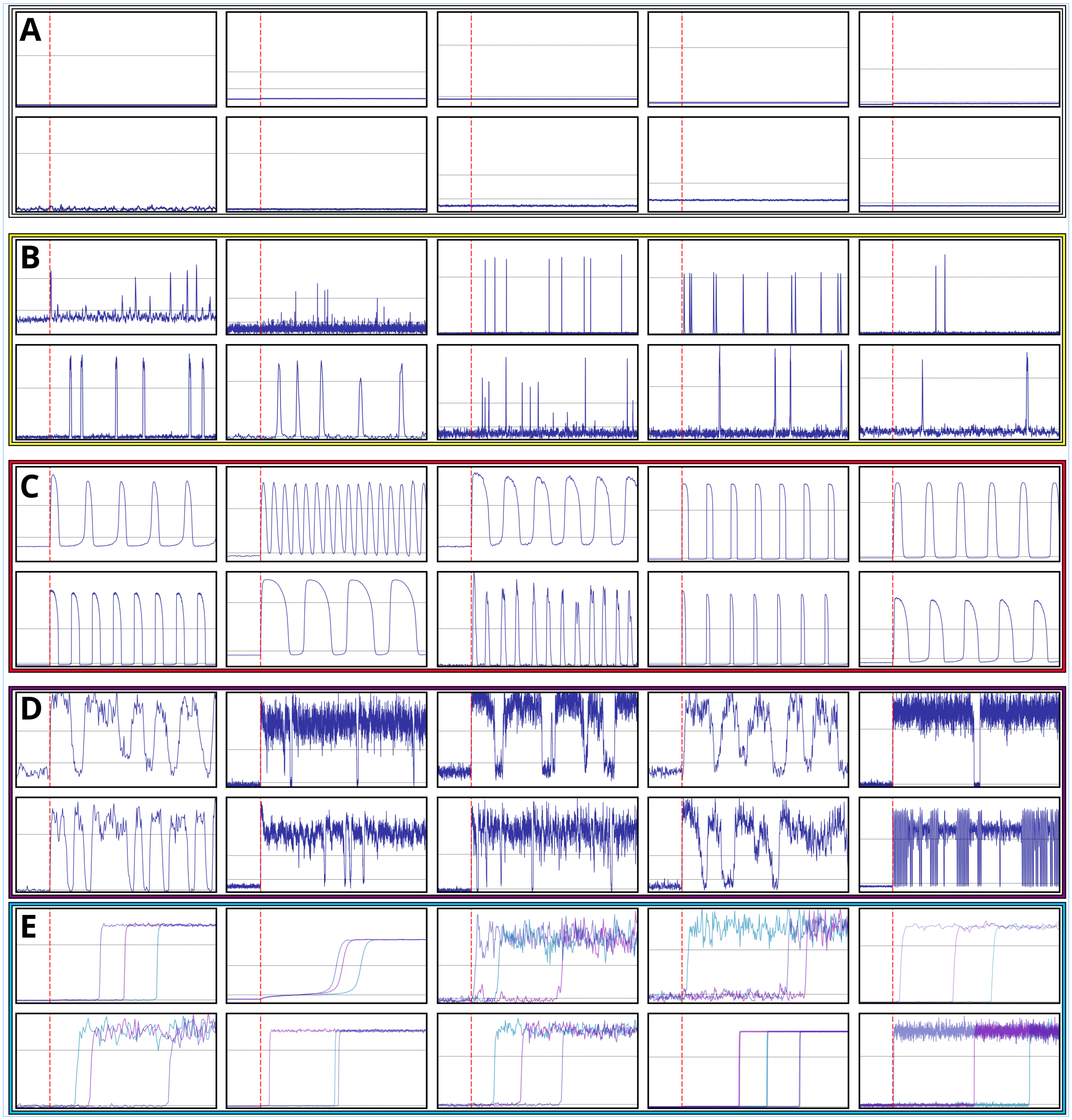

Supplementary Figure 15: **Example model simulations and the algorithm's corresponding classification (part 1).** Random parameter sets were generated and classified according to our algorithm (A-E) Example trajectories from the parameter sets ($\sigma(t)$ in blue, the dotted black lines indicates $T_{ia}$ and $T_a$, and dotted red line the activation of the input). The x-axis is time, and the y-axis is $\sigma$. The time scales may be different between the plots. (A) Simulations for model instances classified as the "no activation" behaviour. (B) Simulations for model instances classified as the "stochastic pulsing" behaviour. (C) Simulations for model instances classified as the "oscillation" behaviour. (D) Simulations for model instances classified as the "stochastic anti-pulsing" behaviour. (E) Simulations for model instances classified as the "heterogeneous activation" behaviour.

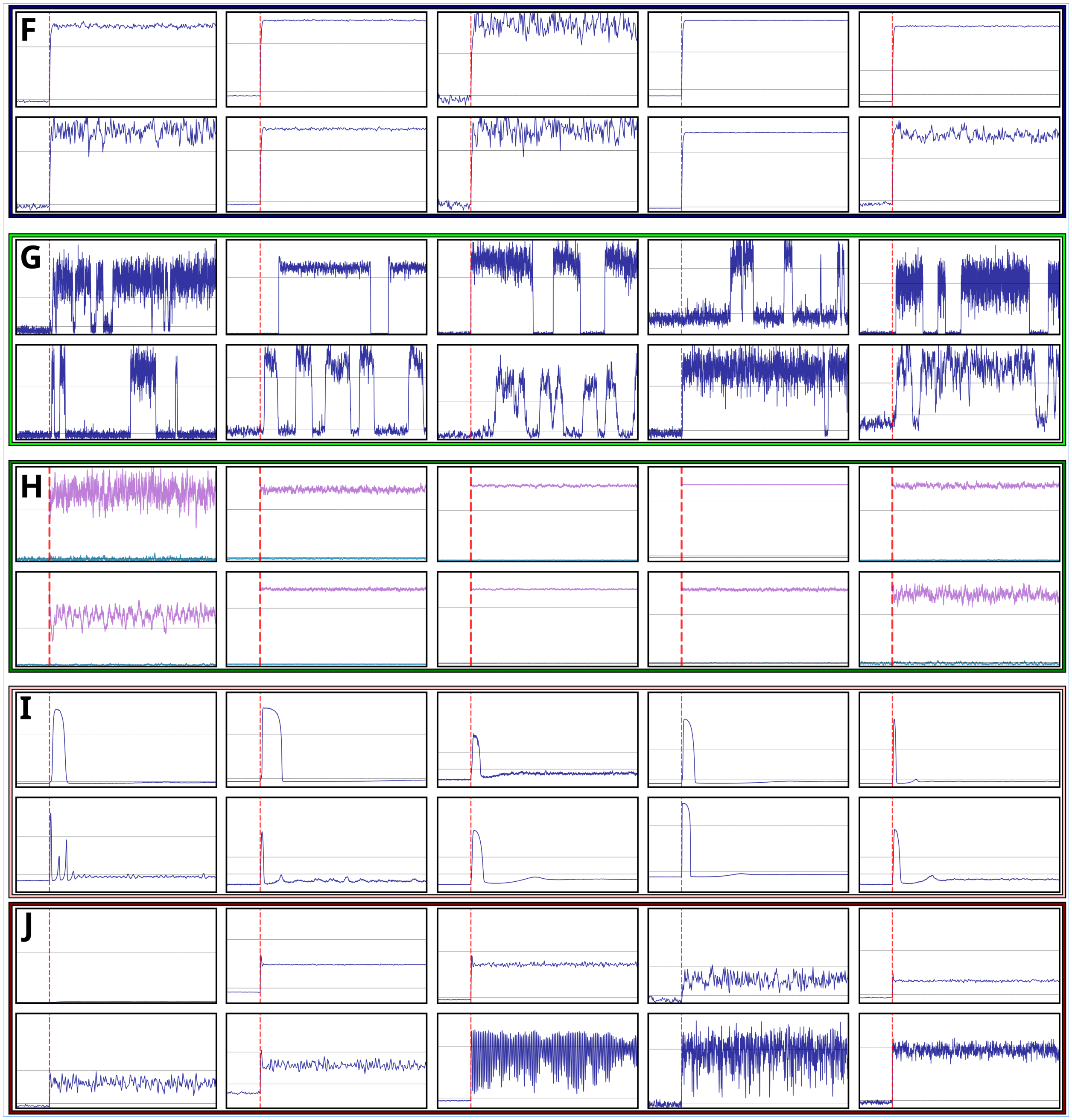

Supplementary Figure 16: **Example model simulations and the algorithm's corresponding classification (part 2).** Random parameter sets were generated and classified according to our algorithm (F-J) Example trajectories from the parameter sets ($\sigma(t)$ in blue, the dotted black lines indicates $T_{ia}$ and $T_a$, and dotted red line the activation of the input). The x-axis is time, and the y-axis is $\sigma$. The time scales may be different between the plots. (F) Simulations for model instances classified as the "homogeneou activation" behaviour. (G) Simulations for model instances classified as the "stochastic switching" behaviour. (H) Simulations for model instances classified as the "stable bistability" behaviour. (I) Simulations for model instances classified as the "single response pulse" behaviour. (J) Simulations for model instances classified as the "intermediate activation" behaviour.

reactivation times. Hence, in this case, the IAT distribution contributed no additional information of the system, and was not used.

Inactive systems with RT heterogeneous were classified as *stochastic pulsing*, while active systems with DT heterogeneous were classified as *stochastic anti-pulsing*. Inactive systems with RT infinite were classified according to their initial activation times (*single response pulse* if instantaneous, else *no activation*). Similarly, active systems were classified as *homogeneous activation* if the IAT were instantaneous, and else as *heterogeneous activation*.

Intermediate systems, inactive systems with instant activation times, and active systems with instant deactivation times were classified according to their frequency properties. The system was simulated for a long time ($11,000$ time units) and the Welch-type periodogram was computed using the last $10,000$ time units of the simulation (with the mean activity subtracted). It was then checked whether the maximum value of the periodogram passed a specific threshold ($T_{freq}$, arbitrarily set to $T_{freq} = 10$). If it did, the system was classified as an oscillation, else as an intermediate activation.

For a bistable system, RT was first classified. If it was infinite, then DT and IAT were found. For DT infinite, and IAT instant, the system was classified as homogeneous activation. For DT infinite, and IAT heterogeneous, the system was classified as *heterogeneous activation*. For DT infinite, and IAT infinite the system was classified as *stable bistability*. If DT was heterogeneous or infinite, the system was classified as a single response pulse if IAT was instantaneous, else it was classified as no activation.

If instead RT was heterogeneous, then DT was measured. If it also was heterogeneous, the system was classified as *stochastic switching*. If DT instead was instantaneous, the median activity of the system was measured (see the previous description). If the median activity was inactive, the system was classified as stochastic pulsing, else it was classified as either *oscillation* or *intermediate activation*, depending on its frequency properties (see the previous description). If DT was infinite, then IAT was measured. If IAT was instantaneous the system was classified as homogeneous activation, else as *heterogeneous activation*.

Finally, if RT was instantaneous, then DT was measured. If DT also was instantaneous the system was classified as oscillation. If it was heterogeneous the median activity of the system was measured (see the previous description). If the median activity was active, the system was classified as stochastic anti-pulsing, else it was classified as either oscillation or intermediate activation, depending on its frequency properties (see the previous description). Finally, if DT was infinite, then the IAT was measured. If IAT was instantaneous, the system was classified as homogeneous activation, else as heterogeneous activation.

#### 6.4 List of behavioural subclasses

While we in this work present 10 different potential response behaviours, these can potentially be subdivided further. While preparing this work, we decided that the 10 behaviours described well represent the potential dynamics of the mixed positive/negative feedback loop. However, for completeness, we here present additional potential behaviours that the model can exhibit.

- **Heterogeneous intermediate activation**: In our classifier, we classify any behaviour which reaches (and stays in) a state of intermediate activity as intermediate activation. However, potentially, it is possible to subdivide this behaviour into *heterogeneous intermediate activation* and *homogeneous intermediate activation* (depending on whether the initial activation time is heterogeneous or homogeneous). We note that while the mean activation time of the normal heterogeneous activation behaviour can get arbitrarily large, the activation times of the heterogeneous intermediate activation behaviour is fairly bounded. Furthermore, this behaviour only occurs in a limited region in the $D \approx 1$ region.
- **Stable bistability with intermediate activation**: Like the stable bistability behaviour, but where one stable steady state occurs for intermediate activity (and the other is an inactive state). Under our scheme, we would classify this as stable bistability.

- **Stochastic switching with intermediate activation**: Like the stochastic switching behaviour, but where one stable steady state occurs for intermediate activity (and the other is an inactive state). Under our scheme, we would classify this as stochastic switching. Indeed, some parameter sets classified as stochastic switching are technically stochastic switching with intermediate activation.

- **Single response pulse with intermediate activation**: A behaviour where there is an initial pulse of activity which reaches the active state, but which then settles to an intermediate activation state. Under our scheme, this behaviour is classified as intermediate activation.

- **Stochastic switching with oscillation and no activation**: In this behaviour, the system randomly switches between an oscillation and an inactive state. This behaviour can be observed in the region of the bifurcation diagrams where the limit cycle coexists with a stable inactive state (Supplementary Figure 11). Generally, this behaviour occurs between the stochastic pulsing and oscillation regions. Under our scheme, this behaviour is classified as stochastic pulsing.

- **Stochastic switching with oscillation and activation**: In this behaviour, the system randomly switches between an oscillation and an active state. This behaviour can be observed in the region of the bifurcation diagrams where the limit cycle coexists with a stable active state (Supplementary Figure 11). Generally, this behaviour occurs between the oscillation and stochastic anti-pulsing regions. Under our scheme, this behaviour is classified as stochastic anti-pulsing.

- **Single response period**: While this behaviour can be derived from our classification grid (Supplementary Figure 12, it was in practice never observed in the model.

- **Heterogeneous deactivation**: The reverse of heterogeneous activation, and practice we have classified this as no activation. This behaviour (occurring between stable bistability and no activation) has two steady state, however, the active one is not long-term stable. Here, an inactive system will remain inactive. Meanwhile, a system (which, through some external means, has been put) in an active state will deactivate after some random time.

### 7 Behaviour descriptions

This section gives a brief overview of the observed behaviours, as well as sample trajectories (Supplementary Figures 17-26).

#### 7.1 No activation

The no activation behaviour is characterised by the system staying constantly inactive (Supplementary Figure 17). Given small values of $S$, or $D \gg S$, the system adopts the no activation behaviour. This is the default behaviour of any system where the input is absent ($S = 0$), or negligible ($S \approx 0$). Since this is a default behaviour, we do not further investigate how the parameters affect its prevalence.

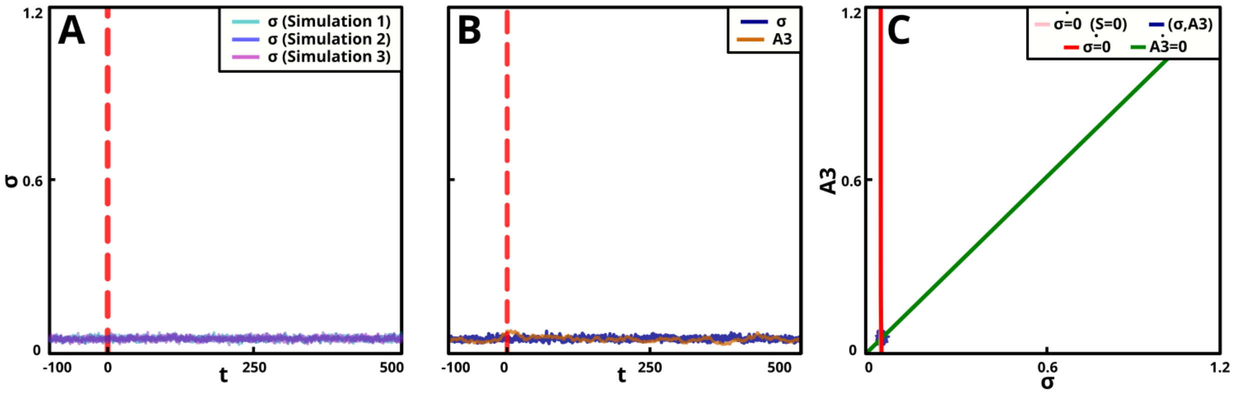

Supplementary Figure 17: **The no activation behaviour.** (A-C) Simulations of the no activation behaviour for the parameter set $(S, D, \tau, v_0, n, \eta) = (1.0, 0.1, 10.0, 0.1, 2, 0.1)$. (A) Three different simulations of the behaviour. (B-C) The same simulation, shown both over time and in phase space. (B) The concentration of the sigma factor ($\sigma$, blue line) and the (third) intermediary ($A = A_3$, yellow line) over time. (C) The state of the system in $[\sigma]$-$[A]$ space (blue trajectory). The nullclines: $NC_A$ (green), and $NC_\sigma$ (pink for input absent, $S = 0$, and red for input present), are also displayed.

#### 7.2 Stochastic pulsing

In the stochastic pulsing behaviour, the system contains only a single steady state, which corresponds to an inactive state. However, random fluctuations may push the system over a boundary that triggers a single pulse of activity. Since this is driven by noise, the pulses occur randomly in time (Supplementary Figure 18). This is a so-called excitable behaviour [Lindner et al., 2004].

The stochastic pulsing behaviour occurs only in the rightmost ($D \gtrsim 1$) region of the $S$-$D$ map. The stochastic pulsing behaviour requires $S$ to be relatively small (so that the system is inactive in its single steady state). For large values of $\eta$, the behaviour can also occur in the ($D \lesssim 1$) region, but this is likely rather a result of a very noisy system occasionally passing the threshold to being active, rather than a genuine pulsing behaviour (Supplementary Figure 28). The behaviour is strongly dependent on the parameters $\tau$ and $\eta$, increasing with their values. Conversely, stochastic pulsing cannot be maintained when either $\tau$ or $\eta$ becomes small. Finally, there is also a slight trend of the behaviour becoming increasingly prevalent with the value of $v_0$ (Supplementary Figure 18D).

Depending on parameter values, the stochastic pulsing behaviour may or may not be accompanied by an initial pulse of $\sigma$ activity at the input's onset (like in the single response pulse behaviour). Typically this occurs for larger values of $S$, while for smaller values the system still exhibits asymptotic pulses, but not necessarily the transient activation pulse. We deemed these two behaviours not different enough to merit specific classifications, but we note the possible distinction.

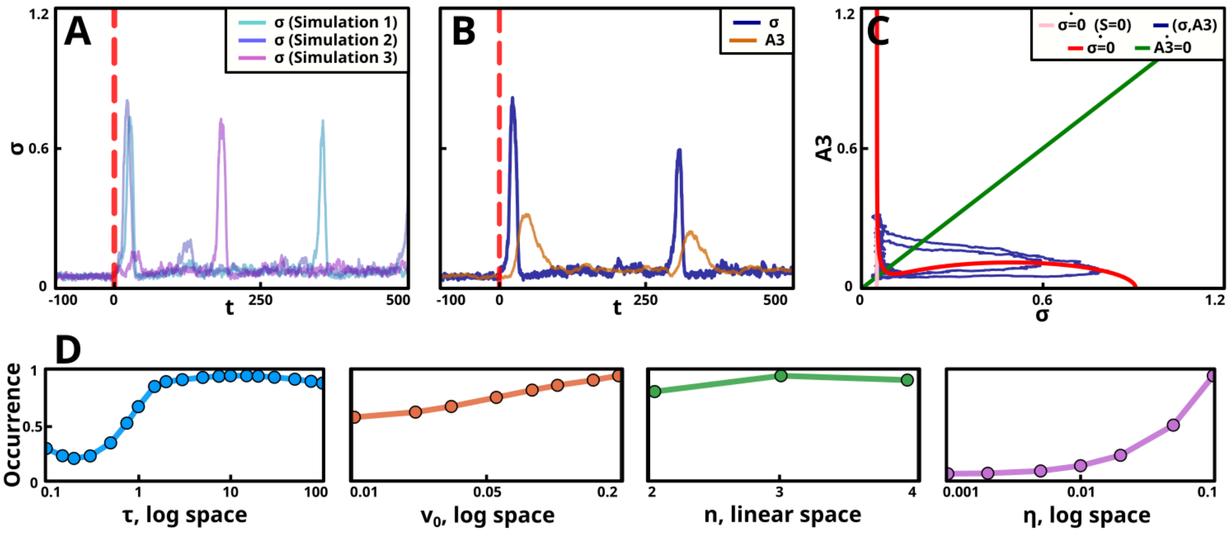

Supplementary Figure 18: **The stochastic pulsing behaviour.** (A-C) Simulations of the stochastic pulsing behaviour for the parameter set $(S, D, \tau, v_0, n, \eta) = (5.0, 50.0, 10.0, 0.05, 2, 0.05)$. (A) Three different simulations of the behaviour. (B-C) The same simulation, shown both over time and in phase space. (B) The concentration of the sigma factor ($\sigma$, blue line) and the (third) intermediary ($A = A_3$, yellow line) over time. (C) The state of the system in [$\sigma$]-[$A$] space (blue trajectory). The nullclines: $NC_A$ (green), and $NC_\sigma$ (pink for input absent, $S = 0$, and red for input present), are also displayed. Note that, for this plot, the y-axis is for a different range than what is used in the other behaviour plots (to ensure that the trajectories can be distinguished clearly) (D) The occurrence analysis of the behaviour shows how its frequency depends on the parameters $\tau$, $v_0$, $n$, and $\eta$ (Supplementary Section 7.11)).

Stochastic pulsing has been observed in a large number of sigma factors ($\sigma^B$, $\sigma^D$, $\sigma^M$, $\sigma^W$, and $\sigma^X$ in *B. subtilis*, as well as RpoS in *E. coli*, [Patange et al., 2018, Park et al., 2018]). It has been suggested as a way to create population heterogeneity, possibly as a part of a bet-hedging behaviour [Patange et al., 2018]. Finally, it has been suggested that several sigma factors may each adopt a stochastic pulsing behaviour as a way of sharing a common resource (RNA Polymerase). Here, at each point in time, only a single sigma factor would be active, and thus utilising the resource [Park et al., 2018].

#### 7.3 Oscillations

In the oscillation behaviour, the system adopts a limit cycle where it oscillates around a single, unstable, steady state (Supplementary Figure 19). Potentially the behaviour could also be generated by the system regularly switching between an active and an inactive state. While this mechanism is accounted for in our classification scheme, we have not confirmed whether it actually generates oscillations in the model.

The oscillation behaviour occurs only in the rightmost ($D \gtrsim 1$) region of the $S$-$D$ map. Here, oscillations require the region's single steady state to be unstable, something which typically occurs for intermediate levels of $S$ (Supplementary Figure 28). The prevalence of the behaviour is mostly unaffected by the levels of $v_0$, $n$, and $\eta$. It does require a relatively large time-delay (large $\tau$) and is absent for small values of $\tau$. This makes sense since negative feedbacks ($D$ non-negligible), when subject to time-delays ($\tau$ large enough), are known to induce limit cycles (Supplementary Figure 19D).

Oscillations have been observed in a cell-free *E. coli* $\sigma^{28}$ system (it is also observed in the $\sigma^B$ mutants investigated in this article). In addition, it occurs in a large number of other cellular systems [Goldbeter, 1991, Nelson et al., 2004, Friedman et al., 2005]. Most notably it is ubiquitous among various circadian rhythm

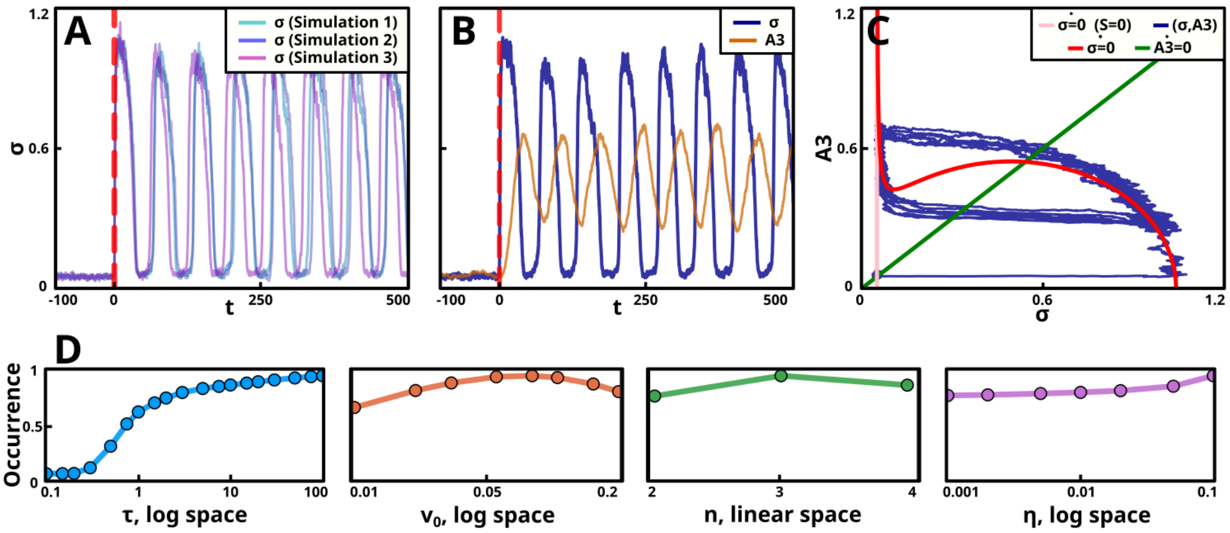

Supplementary Figure 19: **The oscillation behaviour.** (A-C) Simulations of the oscillation behaviour for the parameter set $(S, D, \tau, v_0, n, \eta) = (5.0, 5.0, 10.0, 0.05, 2, 0.05)$. (A) Three different simulations of the behaviour. (B-C) The same simulation, shown both over time and in phase space. (B) The concentration of the sigma factor ($\sigma$, blue line) and the (third) intermediary ($A = A_3$, yellow line) over time. (C) The state of the system in $[\sigma]$-$[A]$ space (blue trajectory). The nullclines: $NC_A$ (green), and $NC_\sigma$ (pink for input absent, $S = 0$, and red for input present), are also displayed. (D) The occurrence analysis of the behaviour shows how its frequency depends on the parameters $\tau$, $v_0$, $n$, and $\eta$ (Supplementary Section 7.11).

systems [Bollinger and Schibler, 2014]).

#### 7.4 Stochastic anti-pulsing

In the stochastic anti-pulsing behaviour, the system contains only a single steady state, which corresponds to an active state. However, random fluctuations may push the system over a boundary, triggering a single (anti-)pulse of inactivity (Supplementary Figure 20). In many ways, it is the reverse of the stochastic pulsing behaviour.

Like the stochastic pulsing behaviour, stochastic anti-pulsing occurs in the rightmost ($D \gtrsim 1$) region of the $S$-$D$ map. However, while stochastic pulsing occurs in the region between the no activity and oscillation behaviours, stochastic anti-pulsing occurs in the region between the oscillation and homogeneous activation behaviours. Like for stochastic pulsing, occurrences of stochastic anti-pulsing in the $D \lesssim 1$ region (when $\eta$ is large) is likely an artefact (Supplementary Figure 28). Just like stochastic pulsing, stochastic anti-pulsing becomes increasingly prevalent with increasing values of $\tau$ and $\eta$ (longer time-delay and more noise, respectively) and disappears when either value becomes small (Supplementary Figure 20D).

To our knowledge, stochastic anti-pulsing has not been observed in any experimental system (neither among sigma factors nor in the wider literature).

#### 7.5 Homogeneous activation

In the homogeneous activation behaviour, at the time of input onset, the system switches from an inactive to an active state. The switch is immediate (and thus homogeneous across an ensemble of activations, as compared to the heterogeneous activation behaviour where it is not). The system is monostable, with only the active steady state (Supplementary Figure 21).

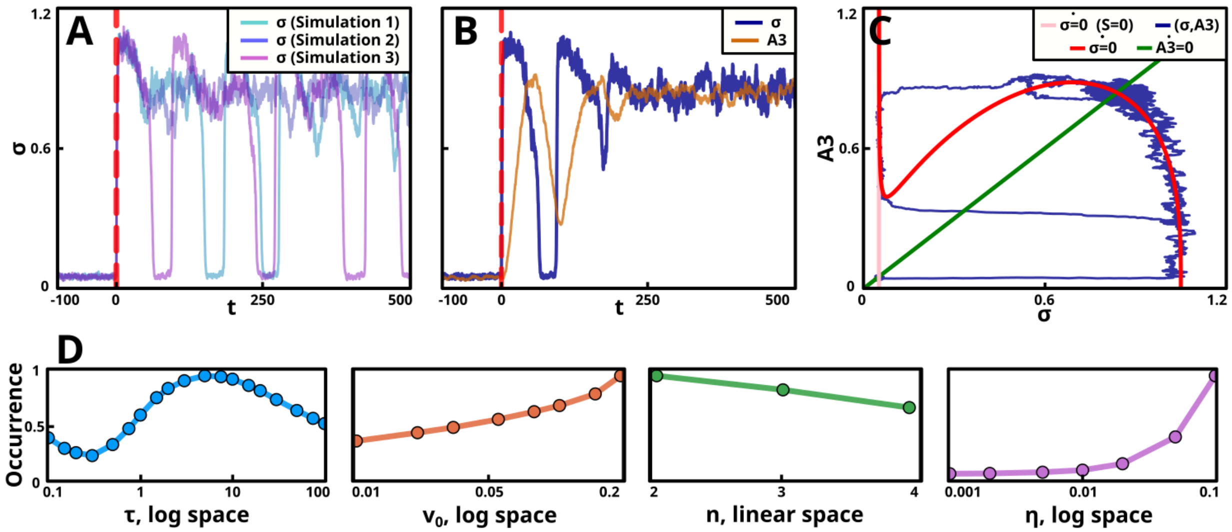

Supplementary Figure 20: **The stochastic anti-pulsing behaviour.** (A-C) Simulations of the stochastic anti-pulsing behaviour for the parameter set $(S, D, \tau, v_0, n, \eta) = (75.0, 50.0, 10.0, 0.05, 2, 0.05)$. (A) Three different simulations of the behaviour. (B-C) The same simulation, shown both over time and in phase space. (B) The concentration of the sigma factor ($\sigma$, blue line) and the (third) intermediary ($A = A_3$, yellow line) over time. (C) The state of the system in $[\sigma]$-$[A]$ space (blue trajectory). The nullclines: $NC_A$ (green), and $NC_\sigma$ (pink for input absent, $S = 0$, and red for input present), are also displayed. (D) The occurrence analysis of the behaviour shows how its frequency depends on the parameters $\tau$, $v_0$, $n$, and $\eta$ (Supplementary Section 7.11).

Given any combination of parameters, if $S$ becomes sufficiently large, a homogeneous activation behaviour is always produced. Also, if $S$ is higher than some threshold value, given a small enough value of $D$ ($D \lesssim S$), a homogeneous activation behaviour will also be produced (Supplementary Figure 28). The occurrence analysis for this behaviour shows similar occurrence frequencies for all parameters. The variations we see are likely due to some parameters enlarging the regions of the other behaviours, reducing the counts for homogeneous activation, rather than the parameters affecting this behaviour directly. (Supplementary Figure 21D).

Theoretically, given a strong enough input, most sigma factors should activate homogeneously. However, the inherent properties of their circuits may put an upper limit to the degree to which the circuit can be activated (preventing this behaviour). The activation of many sigma factors is tied to stress. A strong enough stress will likely kill the bacteria (or cause other secondary effects which might prevent proper activation of the sigma factor), preventing high enough input magnitudes to generate homogeneous activation.

#### 7.6 Heterogeneous activation

In the heterogeneous activation behaviour, at the time of input onset, the system switches from an inactive to an active state, but with highly heterogeneous response times. Typically, the system will have two stable steady states, one with low and one with high activity. Stochastic fluctuations may, however, push the system from the inactive state (where it is at the input's onset) across a separatrix and into the active state. Since this transition is dependent on noise, the activation time of the system is random (and thus heterogeneous across a population). While the inactive state is only stable in the short term (due to noise eventually pushing the system out of it), the active state is long-term stable, with noise-driven transitions from it to the inactive state being impossible in non-absurd timeframes (Supplementary Figure 22).

Since the heterogeneous activation behaviour is dependent on bistability, it only occurs in the leftmost ($D \lesssim 1$) region of the $S$-$D$ map, and only for an intermediate range of $S$ values (the region where bistability

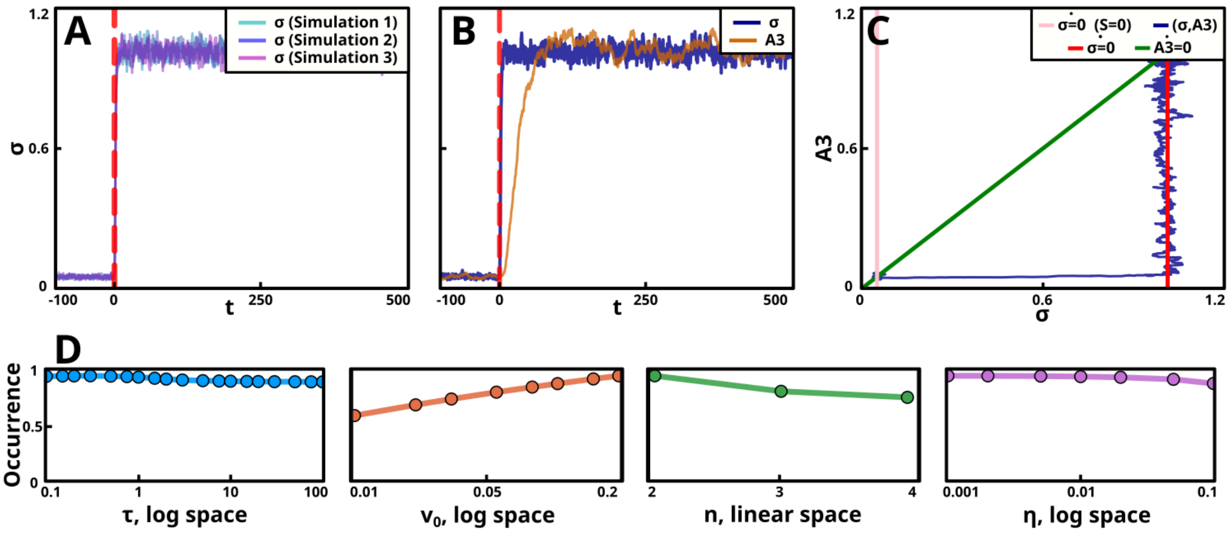

Supplementary Figure 21: **The homogeneous activation behaviour.** (A-C) Simulations of the homogeneous activation behaviour for the parameter set $(S, D, \tau, v_0, n, \eta) = (3.0, 0.1, 10.0, 0.05, 2, 0.05)$. (A) Three different simulations of the behaviour. (B-C) The same simulation, shown both over time and in phase space. (B) The concentration of the sigma factor ($\sigma$, blue line) and the (third) intermediary ($A = A_3$, yellow line) over time. (C) The state of the system in $([\sigma], [A]$ space (blue$NC_\sigma$ trajectory). The nullclines: $NC_A$ (green), and $NC_\sigma$ (pink for input absent, $S = 0$, and red for input present), are also displayed. (D) The occurrence analysis of the behaviour shows how its frequency depends on the parameters $\tau$, $v_0$, $n$, and $\eta$ (Supplementary Section 7.11).

occurs, Supplementary Figure 28). In addition, the prevalence of the behaviour increases with the value of $n$ (increased ultrasensitivity is associated with more distinct bistability) and $\eta$ (the system is clearly dependent on noise to make the inactive state long-term unstable). For large values of $v_0$ the behaviour is diminished (this is in agreement with Supplementary Figure 11, which suggests small $v_0$ promotes bistability). This also holds as $\eta$ becomes large, likely due to high amplitude noise enabling transitions from the active to the inactive state (yielding a stochastic switching behaviour), or alternatively, makes the inactive state so unstable that activations become homogeneous (Supplementary Figure 22D).

Just like there exists bistable systems where the active state is long-term stable and the inactive one is not, there exist those with a long-term stable inactive state and an active state which is only short-term stable. Here, when the input parameter is changed from the value 0 to $S$, the system remains inactive. However, if the parameter is increased further (enough to activate the system), and then reduced back $S$ again, one would see a heterogeneous deactivation system. We deemed this feature not interesting enough to distinguish it from the no activation behaviour.

The $\sigma^V$ system of *B. subtilis* has been observed to activate heterogeneously in response to lysozyme stress [Schwall et al., 2021]. In addition, the behaviour has been observed in non-sigma factor systems [Weber and Buceta, 2013, Uphoff et al., 2016].

#### 7.7 Stochastic switching

In stochastic switching, the system has two steady states, neither of which are long-term stable. Noise may drive the system from either steady state, across the separatrix, and into the other state. These switches occur randomly in time (Supplementary Figure 23).

The stochastic switching behaviour occurs in the leftmost ($D \lesssim 1$) region of the $S$-$D$ map, typically for val-

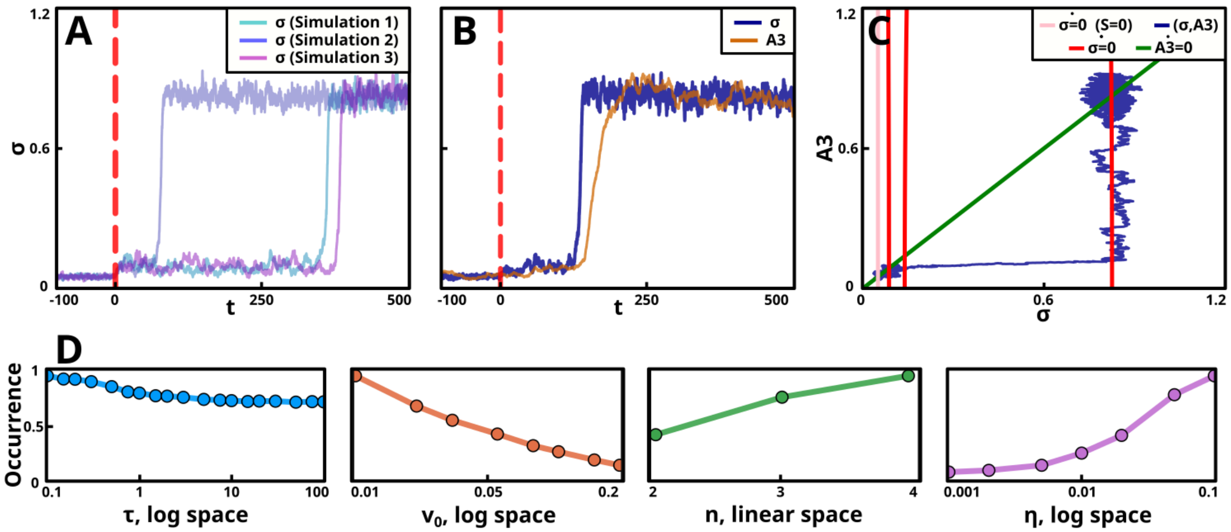

Supplementary Figure 22: **The heterogeneous activation behaviour.** (A-C) Simulations of the heterogeneous activation behaviour for the parameter set $(S, D, \tau, v_0, n, \eta) = (2.2, 0.1, 10.0, 0.05, 2, 0.05)$. (A) Three different simulations of the behaviour. (B-C) The same simulation, shown both over time and in phase space. (B) The concentration of the sigma factor ($\sigma$, blue line) and the (third) intermediary ($A = A_3$, yellow line) over time. (C) The state of the system in $[\sigma]$-$[A]$ space (blue trajectory). The nullclines: $NC_A$ (green), and $NC_\sigma$ (pink for input absent, $S = 0$, and red for input present), are also displayed. (D) The occurrence analysis of the behaviour shows how its frequency depends on the parameters $\tau$, $v_0$, $n$, and $\eta$ (Supplementary Section 7.11).

ues of $S$ lower than those that yield heterogeneous activation, but larger than what yields no activity. The behaviour also occurs in the centre of the plot, at $D \approx S \approx 1$ (Supplementary Figure 28). As a noise-driven behaviour, it requires a large enough value of $\eta$, and does not occur when $\eta$ is small. It is also slightly less prevalent for small values of $\tau$ and large values of $v_0$ (Supplementary Figure 23D).

The stochastic switching behaviour has been observed in several cellular systems [Acar et al., 2008, Sureka et al., 2008], but not to our knowledge among bacterial sigma factors.

#### 7.8 Stable bistability

In the stable bistability behaviour, the system has two steady states, both of which are long-term stable (no asymptotic switching between the two states occurs). At the activation of the input, the system will remain in its inactive state, and is in that sense no different from the no activation behaviour. However, some external factor could cause a switch to the active state (at which point it will remain there until further external perturbations cause a switch to the inactive state). Such a factor could be increasing $S$ so that the system activates through a heterogeneous or homogeneous activation behaviour. When $S$ is then reduced to the value yielding stable bistability, the system will be active instead of inactive. Alternatively, an external source of $\sigma$ could be added, pushing the system to the active state (Supplementary Figure 24).

Like the previous bistability-based behaviours, stable bistability occurs in the leftmost ($D \lesssim 1$) region of the $S$-$D$ map, and only for an intermediate range of $S$ values (for higher values of $S$ than no activity, but lower than heterogeneous activation, Supplementary Figure 28). Likely due to its dependency on bistability, the behaviour becomes more prevalent with increasing $n$. It is also dependent on having a low value for $\eta$ (since increasing noise may create transitions between the two steady states). The behaviour also disappears for large values of $v_0$. Since it occurs for low values of $D$, where self-deactivation is weak, $\tau$ has little effect on its prevalence (Supplementary Figure 24D).

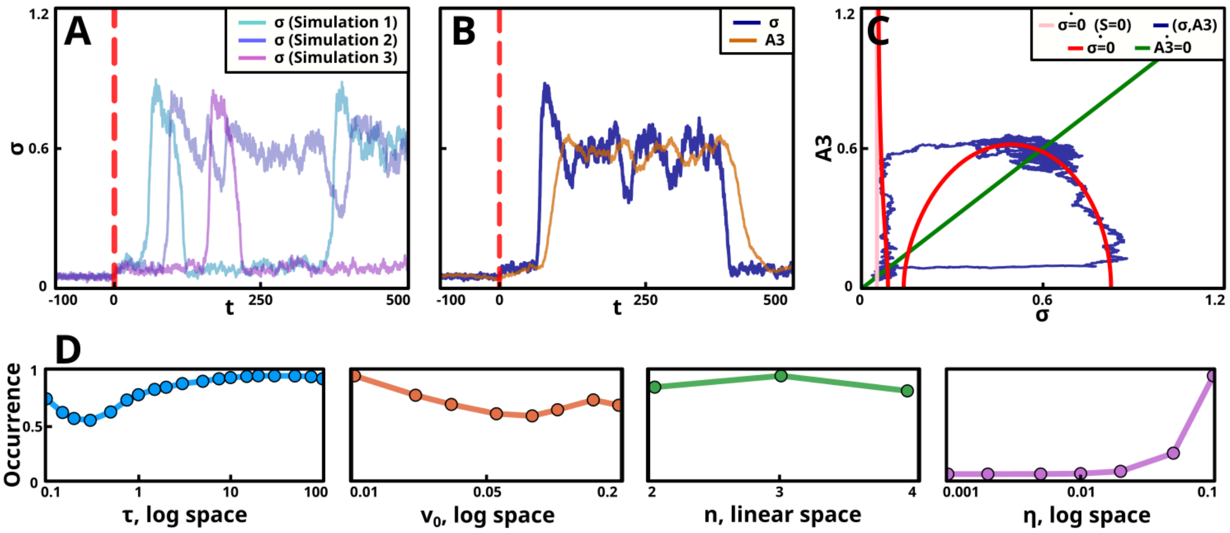

Supplementary Figure 23: **The stochastic switching behaviour.** (A-C) Simulations of the stochastic switching behaviour for the parameter set $(S, D, \tau, v_0, n, \eta) = (2.25, 1.1, 10.0, 0.05, 2, 0.05)$. (A) Three different simulations of the behaviour. (B-C) The same simulation, shown both over time and in phase space. (B) The concentration of the sigma factor ($\sigma$, blue line) and the (third) intermediary ($A = A_3$, yellow line) over time. (C) The state of the system in $[\sigma]$-$[A]$ space (blue trajectory). The nullclines: $NC_A$ (green), and $NC_\sigma$ (pink for input absent, $S = 0$, and red for input present), are also displayed. (D) The occurrence analysis of the behaviour shows how its frequency depends on the parameters $\tau$, $v_0$, $n$, and $\eta$ (Supplementary Section 7.11).

Stable bistability type behaviours are well known in development, where the selection of a system steady state may act as an irreversible decision of the developmental fate of a cell [Choi et al., 2008, Frigola et al., 2012]. To our knowledge, it has not been observed among bacterial sigma factors.

#### 7.9 Single response pulse

In the single response pulse behaviour, the system is inactive as the input persists. However, the input's onset causes a large enough perturbation to the system that it displays a single pulse of activity (Supplementary Figure 25). This is an excitable behaviour [Lindner et al., 2004]. Especially for low-noise regimes, this behaviour may be generated by a damped oscillation (Supplementary Figure 16).

Since the behaviour is dependant on excitability, it occurs in the rightmost ($D \gtrsim 1$) region of the $S$-$D$ map. It requires small values of $S$ (where the steady state is inactive). However, the values should be larger than what yield a no activation behaviour. We note that the single response pulse behaviour typically occupies a very slim range of $S$ values. However, since the model is nondimensionalised, this region may correspond to a large region in a real system. As an excitable behaviour, it is reliant on a large enough time-delay (and thus disappears as $\tau$ becomes small). The behaviour also requires $\eta$ to be small. Since the pulse is due to a deterministic phenomenon (input onset changing the steady state properties of the system) it is not dependent on noise. However, for larger $\eta$, stochasticity becomes prominent enough to cause the excitability to be triggered by stochastic fluctuations as well, turning the behaviour into a stochastic pulsing one. The behaviour becomes more prevalent with larger values of $v_0$ and smaller values of $n$ (Supplementary Figure 25D). The behaviour typically occurs in very slim regions of parameter space (Supplementary Figure 29), however, due to the nondimensionalised nature of the model, we cannot draw major conclusions for what this means for real systems.

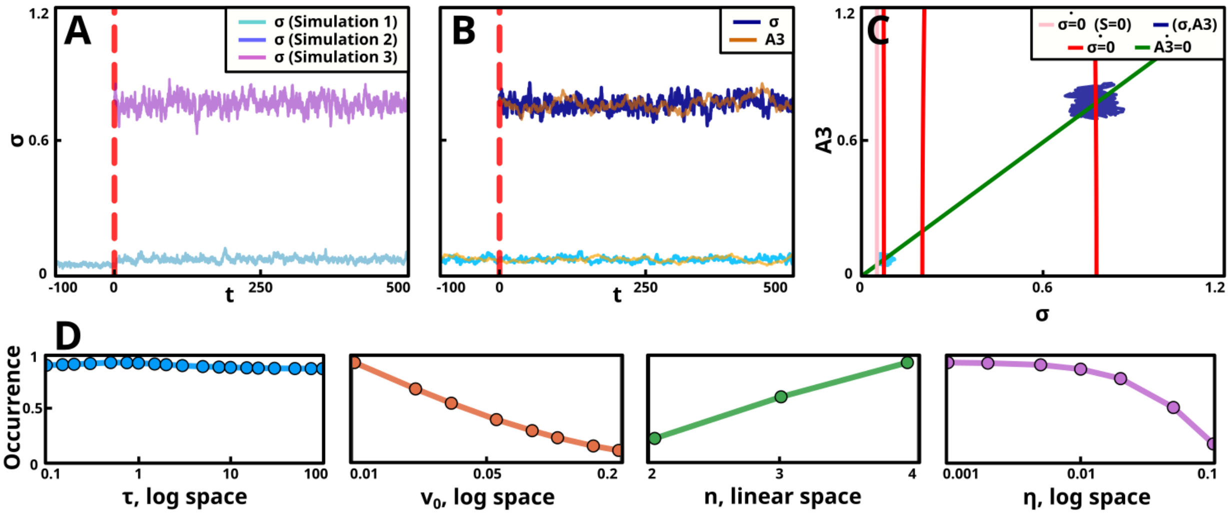

Supplementary Figure 24: **The stable bistability behaviour.** (A-C) Simulations of the stable bistability behaviour for the parameter set $(S, D, \tau, v_0, n, \eta) = (2.5, 0.1, 10.0, 0.025, 2, 0.05)$. (A) Three different simulations of the behaviour. Simulations 1 and 2 are performed in the inactive state, and simulation 3 in the active state (only present in the presence of the input). (B-C) Two sets of simulations, each one shown over time and in phase space. (B) Two simulations of the model, one in the system's inactive state and one in its active state. For each simulation, both the concentration of the sigma factor ($\sigma$, blue line) and the (third) intermediary ($A = A_3$, yellow line) are shown (lighter colours correspond to the inactive state simulation and darker colours to the active state simulation). (C) The same simulations as in B, with their trajectories shown in $[\sigma]$-$[A]$ space (blue trajectories). The nullclines: $NC_A$ (green), and $NC_\sigma$ (pink for input absent, $S = 0$, and red for input present), are also displayed. (D) The occurrence analysis of the behaviour shows how its frequency depends on the parameters $\tau$, $v_0$, $n$, and $\eta$ (Supplementary Section 7.11).

The $\sigma^B$ system in *B. subtilis* yields a single response pulse to the onset of environmental stress [Young et al., 2013]. In addition, the antithetic feedback loop (a network motif which maintains the concentration of a regulated species at some stable level) can also generate single response pulses when its input changes [Briat et al., 2016]. However, while these motifs have similarities, they seem qualitatively different, as the antithetic feedback loop has not been related to stochastic pulsing.

##### 7.10 Intermediate activation

In the intermediate activation behaviour, the system instantaneously activates in response to the input, but instead of reaching an active state (like in homogeneous activation), the system stops at some intermediate value (Supplementary Figure 26).

The intermediate activation behaviour occurs in the rightmost ($D \gtrsim 1$) region of the $S$-$D$ map. It can be found for intermediate values of $S$, where the system's single steady state is neither inactive nor active (Supplementary Figure 28). The behaviour is also dependent on the value of $\tau$. For large values of $\tau$ the steady state is unstable for intermediate values, yielding an oscillation. Only as $\tau$ becomes small does intermediate activation occur. The behaviour's prevalence has little dependency on the other parameter values. The slight increase for large $\eta$ is likely due to noise causing otherwise inactive states to spend long enough time in intermediate values to be deemed as intermediate activation (Supplementary Figure 26D).

For $D \approx 1$, one sometimes encounters a behaviour that can be considered heterogeneous intermediate activation, where the system reaches an intermediate steady state but with a random (non-instant) activation time. This behaviour only occurs for a slim parameter region (adjacent to the intermediate activation region) and reaches a state very close to being active. Also, unlike the normal heterogeneous activation behaviour,

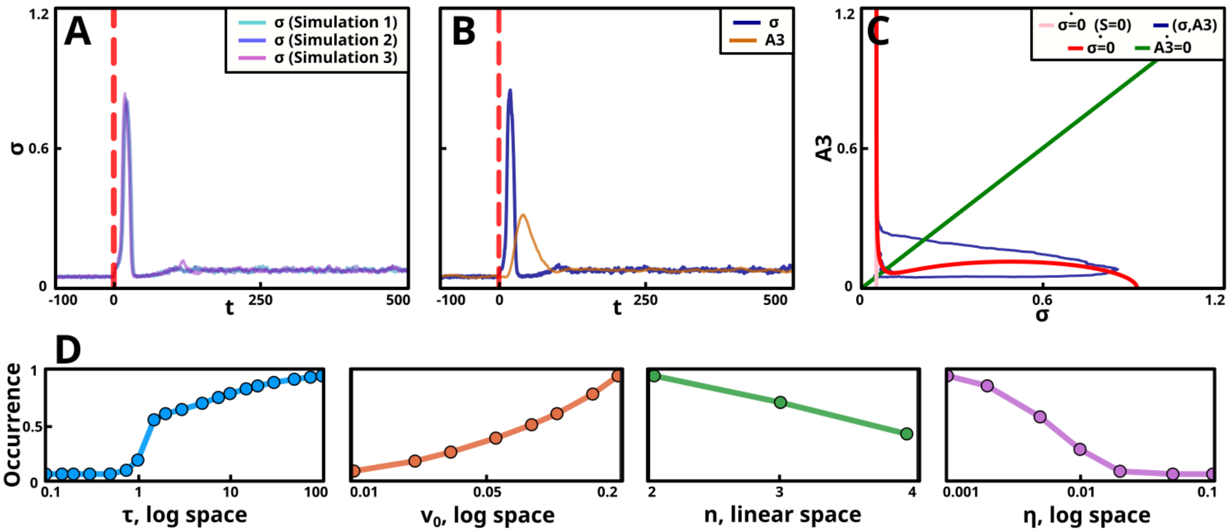

Supplementary Figure 25: **The single response pulse behaviour.** (A-C) Simulations of the single response pulse behaviour for the parameter set $(S, D, \tau, v_0, n, \eta) = (2.2, 6.0, 10.0, 0.1, 2, 0.025)$. (A) Three different simulations of the behaviour. (B-C) The same simulation, shown both over time and in phase space. (B) The concentration of the sigma factor ($\sigma$, blue line) and the (third) intermediary ($A = A_3$, yellow line) over time. (C) The state of the system in $[\sigma]$-$[A]$ space (blue trajectory). The nullclines: $NC_A$ (green), and $NC_\sigma$ (pink for input absent, $S = 0$, and red for input present), are also displayed. (D) The occurrence analysis of the behaviour shows how its frequency depends on the parameters $\tau$, $v_0$, $n$, and $\eta$ (Supplementary Section 7.11).

i

where the mean activation time can get arbitrarily large, the mean activation time for this behaviour seems bound to a relatively narrow range. Hence we decided this behaviour did not merit its own classification.

In practice, it may be hard to distinguish intermediate activation from homogeneous activation. This could likely only be done when one may vary $S$ to see whenever the single steady state smoothly transitions from inactive to active, assuming intermediately active states in between (e.g. like what was done in Figure 4). Also, for larger values of $\eta$, the intermediate activation behaviour displays large fluctuations, from inactive to active states. This makes it hard to distinguish whether noisy experimental data is an intermediate activation behaviour, or simply large degrees of technical noise added upon some other behaviour. So while intermediate activation has likely been observed in both sigma factor and non-sigma factor systems, it is hard to determine whenever it actually was this behaviour that was observed.

#### 7.11 Behaviour occurrence analysis

In Supplementary Figures 17-26 we, for each behaviour, measure how the frequency of its occurrence depends on the four parameters $\tau$, $v_0$, $n$, and $\eta$. This section details how this analysis was performed. Consider our parameter grid of evaluated behaviours, $B(S_i, D_i, \tau_i, v_{0,i}, n_i, \eta_i)$. We define the count $\hat{C}(b, \tau_j)$ as the number of times the behaviour $b$ occurs in the grid $B(...)$ for $\tau_i$ assuming some specific value $\tau_j$. Here,

$$b \in \{\text{no activation, stochastic pulsing, oscillation, ...}\}$$

We then create the occurrence vector for the behaviour $b$ and the grid $\{\tau_i\}$: $\hat{C}_{\tau,i}(b)$. Finally, we normalise $\hat{C}_{\tau,i}(b)$ by dividing every element by its maximum value. This creates $C_{\tau,i}(b)$, for which $0 \leq C_{\tau,i}(b) \leq 1 \;\forall\; i$ holds. We create similar occurrence vectors for $v_0$, $n$, and $\eta$ (vectors could also have been created for $S$ and $D$, but we found it simpler to consider the maps in Supplementary Figure 28).

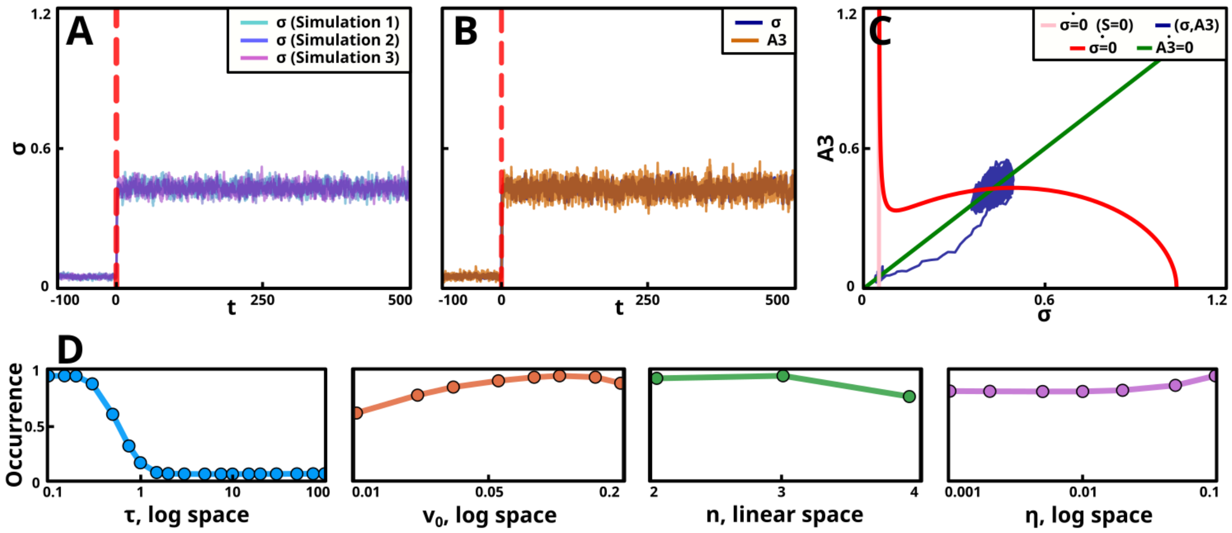

Supplementary Figure 26: **The intermediate activation behaviour.** (A-C) Simulations of the intermediate activation behaviour for the parameter set $(S, D, \tau, v_0, n, \eta) = (5.0, 5.0, 0.5, 0.05, 2, 0.05)$. (A) Three different simulations of the behaviour. (B-C) The same simulation, shown both over time and in phase space. (B) The concentration of the sigma factor ($\sigma$, blue line) and the (third) intermediary ($A = A_3$, yellow line) over time. (C) The state of the system in $[\sigma]$-$[A]$ space (blue trajectory). The nullclines: $NC_A$ (green), and $NC_\sigma$ (pink for input absent, $S = 0$, and red for input present), are also displayed. (D) The occurrence analysis of the behaviour shows how its frequency depends on the parameters $\tau$, $v_0$, $n$, and $\eta$ (Supplementary Section 7.11).

During the count, $n = 2$ was discounted from the grid (except for when measuring the occurrence for the parameter $n$). The reason was that, for $n = 2$, for some values of $v_0$, the system's nullcline lacked local minimum and maximum, and the system could only be classified as no activation or homogeneous activation (Supplementary Section 6.2). This produced artefacts in the occurrence plots. Also, the highest value of $\eta$ (0.20) was discounted as for this value a few grids had missing grid values (where we had avoided making a classification since the condition of the CLE had been broken).

### 8 Behaviour maps

#### 8.1 Generating behaviour maps

We have divided the possible behaviours of our model into ten different classes (Supplementary Section 6.1). Our classification algorithm can be used to automatically assign a parameter set to one of these classes (Supplementary Section 6.3). To create our behaviour maps, we classified the system for all parameter values in a parameter grid: $\{S_i, D_i, \tau_i, v_{0,i}, n_i, \eta_i\}$. The values in $S_i$ and $D_i$ (for which a larger number of values was required to yield high resolution maps) were log10-distributed. The values of $\tau_i$, $v_{0,i}$, and $\eta_i$ were approximately log-distributed (but hand-picked, to avoid long decimals when writing them out). $n_i$ was set to $\{2, 3, 4\}$.

The minimum and maximum values of $S_i$ and $D_i$ were chosen so that further decreases (or increases) did not noticeably change the maps (Supplementary Figure 29-32). The minimum values of $\tau$, $v_0$, and $\eta$ were chosen so that the time-delay, base production, and noise levels, respectively, were negligible. For the maximum value of $\tau$ the time delay was so large that further increases just made limit cycles longer, but no additional changes in behaviours were produced. For the maximum value of $v_0$ (0.2), it constituted a significant portion of the maximum production (1.0). For the maximum value of $\eta$ (0.20) noise became so large that it obscured other behaviours (also, large noise fluctuations would increasingly make the CLE approximation less reliable). It was assumed that $n$ (if $> 1$) would have no major impact on the system, and so only three values were chosen (this assumption is mostly validated in Supplementary Figures 17-26). Finally, we checked that the behaviour maps stopped changing as the parameter values reached their minimum and maximum values (Supplementary Figures 29-32).

Since the classifier was dependant on stochastic simulations, the classification of each parameter value was random. To reduce the noise we postulated that the behaviour of each parameter set was likely similar to that of its neighbours. In a single behaviour map, each grid point has 8 neighbouring grid points (in the $S$-$D$ space). The colour of a grid point was chosen by a single, weighed, vote of the point and its neighbours (where the point's colour was weighed 3, its direct neighbour's colours were weighed 2, and its diagonal neighbour's colours were weighed 1).

#### 8.2 Bifurcation diagrams across the behaviour map

We note that for small values of $S$, the system remains inactive (this corresponds to the behaviour when there is no input). As $S$ is increased, the system passes through a region with some non-trivial behaviour (behaviour which is neither no activity nor homogeneous activation), to reach the homogeneous activation behaviour for large $S$ (corresponding to a system immediately activating, and staying active, once the input is activated). Meanwhile, we see that $D$ splits parameter space into two regions, each with its own distinct set of behaviours. For small $D$ ($D \lesssim 1$) the system experiences a region of bistability as $S$ is varied. This region includes the heterogeneous activation, stochastic switching, and stable bistability behaviours. For large $D$ ($D \gtrsim 1$) the system experiences a smooth, monostable, transition from an inactive to an active state as $S$ is increased (for large $\tau$, the single state may be unstable). This region contains the stochastic pulsing, oscillation, stochastic anti-pulsing, single response pulse, and intermediate activation behaviours (Supplementary Figure 27). The single response pulse and intermediate activation behaviours does not occur for the parameter set used in Supplementary Figure 27, but can be found in Supplementary Figure 28.

#### 8.3 Additional behaviour maps

To complement the behaviour map shown in the main article, we here show a large number of additional behaviour maps, permitting more careful analysis of how the parameters affect the system's behaviour.

First, we consider a succession of $S$-$D$ space behaviour maps, where a single parameter (either $\tau$, $v_0$, $n$, or $\eta$) is modulated across the series. Here, we can see how changing that parameter affects the system's behaviour (Supplementary Figure 28).

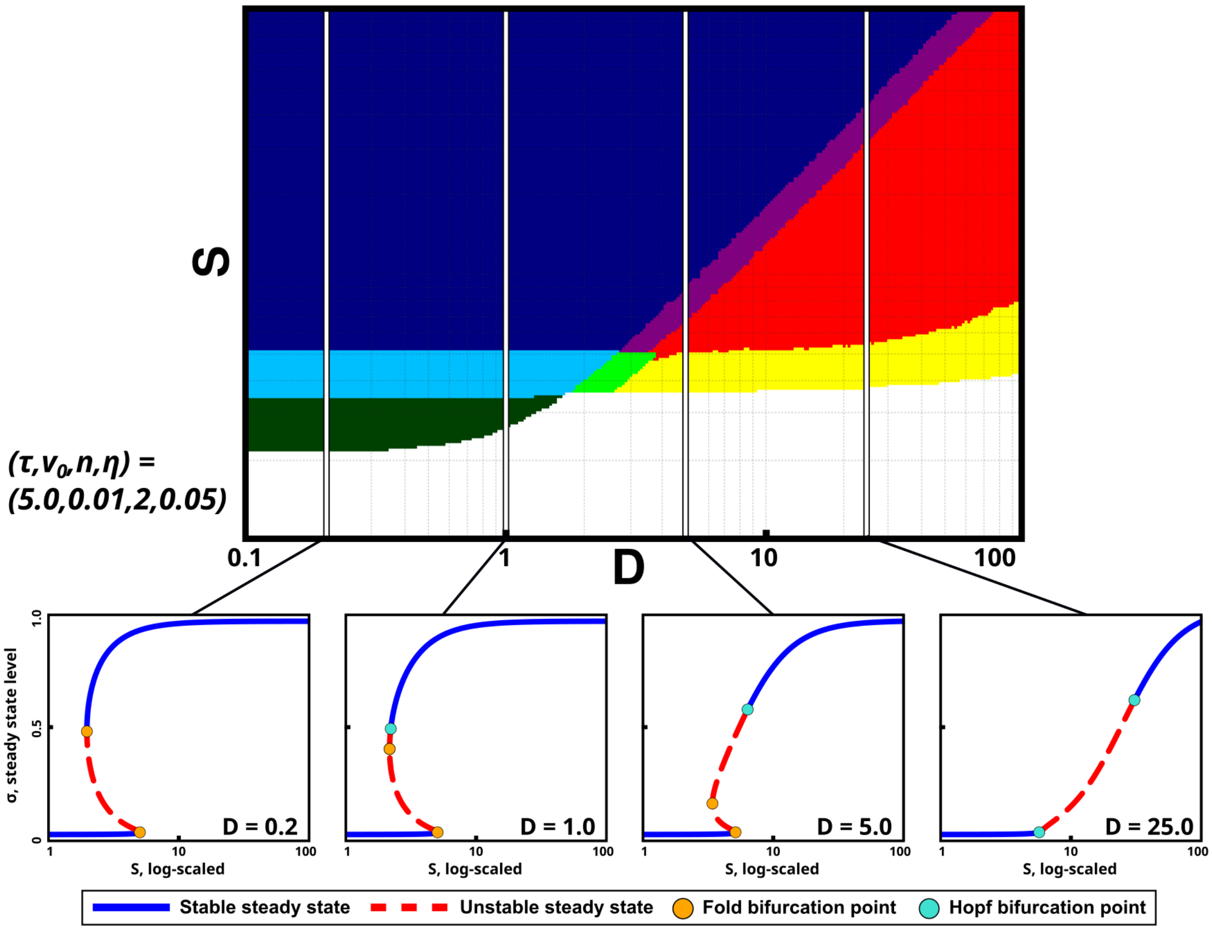

Supplementary Figure 27: **The behaviour map consists of a bistable and a single steady state part.** The same behaviour plot as in Figure 3. In addition, four lines indicate ranges of $S$ for which the displayed bifurcation diagrams are created. In the leftmost part of the behaviour plot ($D \lesssim 1$) the system displays bistability, while in the rightmost part ($D \gtrsim 1$) the system only have a single (possibly unstable) steady state. The two regions are also characterised by two distinct sets of behaviours.

If we consider $\tau$, it has little effect on the leftmost part of the behaviour space (for low $D$ values, where the system is bistable). This makes sense, since in this region self-deactivation is weak (which is the feedback the time-delay affects). In the rightmost part, $\tau$ affects the transition from oscillations (high $\tau$) to intermediate activation (low $\tau$). Again, it makes sense that oscillations require a high enough $\tau$, since these rely on the presence of a time-delay. Finally, reducing $\tau$ also reduces the size of the stochastic pulsing and stochastic anti-pulsing regions. Again, this makes sense since these are excitable behaviours, which are related to oscillations (Supplementary Figure 28A).

Tuning $v_0$ has little effect on the types of behaviours encountered, however, the regions of the non-trivial behaviours are enlarged as $v_0$ is reduced. This especially holds for the behaviours depending on bistability (heterogeneous activation, stochastic switching, and stable bistability), as well as oscillations (Supplementary Figure 28B). This agrees with the bifurcation diagrams, where a low $v_0$ was shown to extend the bistability region (Supplementary Figure 11).

Tuning of the noise parameter, $\eta$, primarily affects the system's noise dependent behaviours. All these (stochastic pulsing, stochastic anti-pulsing, stochastic switching, and heterogeneous activation) diminish as

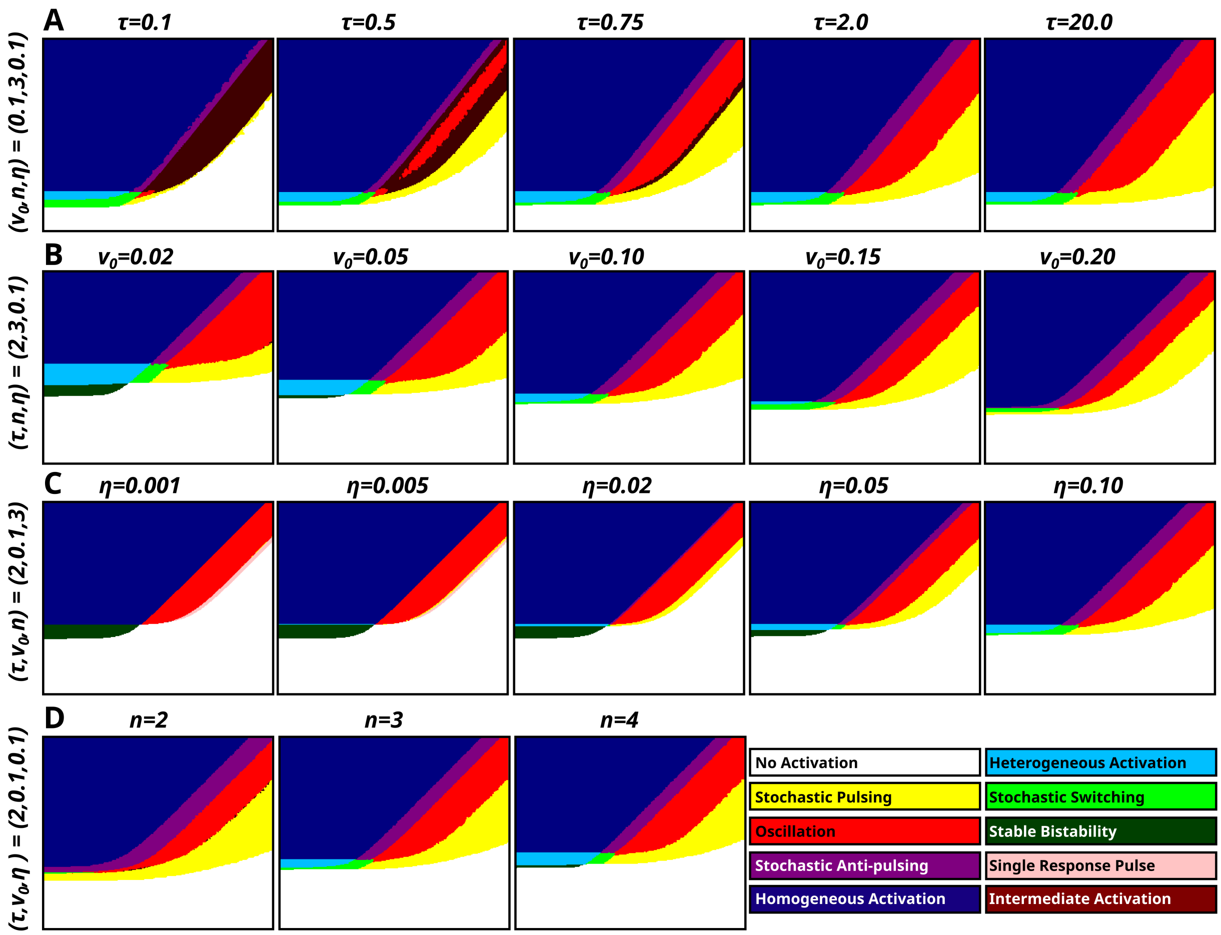

Supplementary Figure 28: **The behaviour map is modulated by the remaining four parameters.** (A-D) For all behaviour maps, the parameter set $(\tau, v_0, n, \eta) = (2.0, 0.1, 3, 0.1)$, with $S \in (1.0, 100.0)$, and $D \in (0.1, 100.0)$ is used (however, for each map one parameter is modified, as described above it in the figure). All maps are created as described in Supplementary Section 8 (A) The behaviour map for a range of $\tau$ values. (B) The behaviour map for a range of $v_0$ values. (C) The behaviour map for a range of $\eta$ values. (D) The behaviour map for a range of $n$ values.

$\eta$ is reduced. The stochastic switching region becomes larger (to a limit) as $\eta$ increases. The heterogeneous activation region also becomes larger, but then shrinks and disappears as $\eta$ becomes large enough. This is likely caused by noise becoming so large that the inactive state cannot be maintained. Finally, the stochastic pulsing and stochastic anti-pulsing regions become larger as $\eta$ increases. Most notably, these regions extend into the $D \lesssim 1$ region for very high $\eta$. This might not be stochastic (anti-)pulsing due to excitability (as in the $D \gtrsim 1$ region), but is rather caused by the stochastic fluctuations becoming so large that the system will not stay reliably inactive/active. Also, the single response pulse behaviour seems to require $\eta$ to be low enough (Supplementary Figure 28C).

The primary effect of changing $n$ (the degree of ultrasensitivity) seems to be to increase the sizes of the non-trivial behaviour regions (Supplementary Figure 28D).

Finally, it is also possible to create a 2d grid of parameter maps, showing the changes with regards to two different parameters. These show similar trends as shown in Supplementary Figure 28. For a number of such grids of maps, please consider Supplementary Figures 29-32.

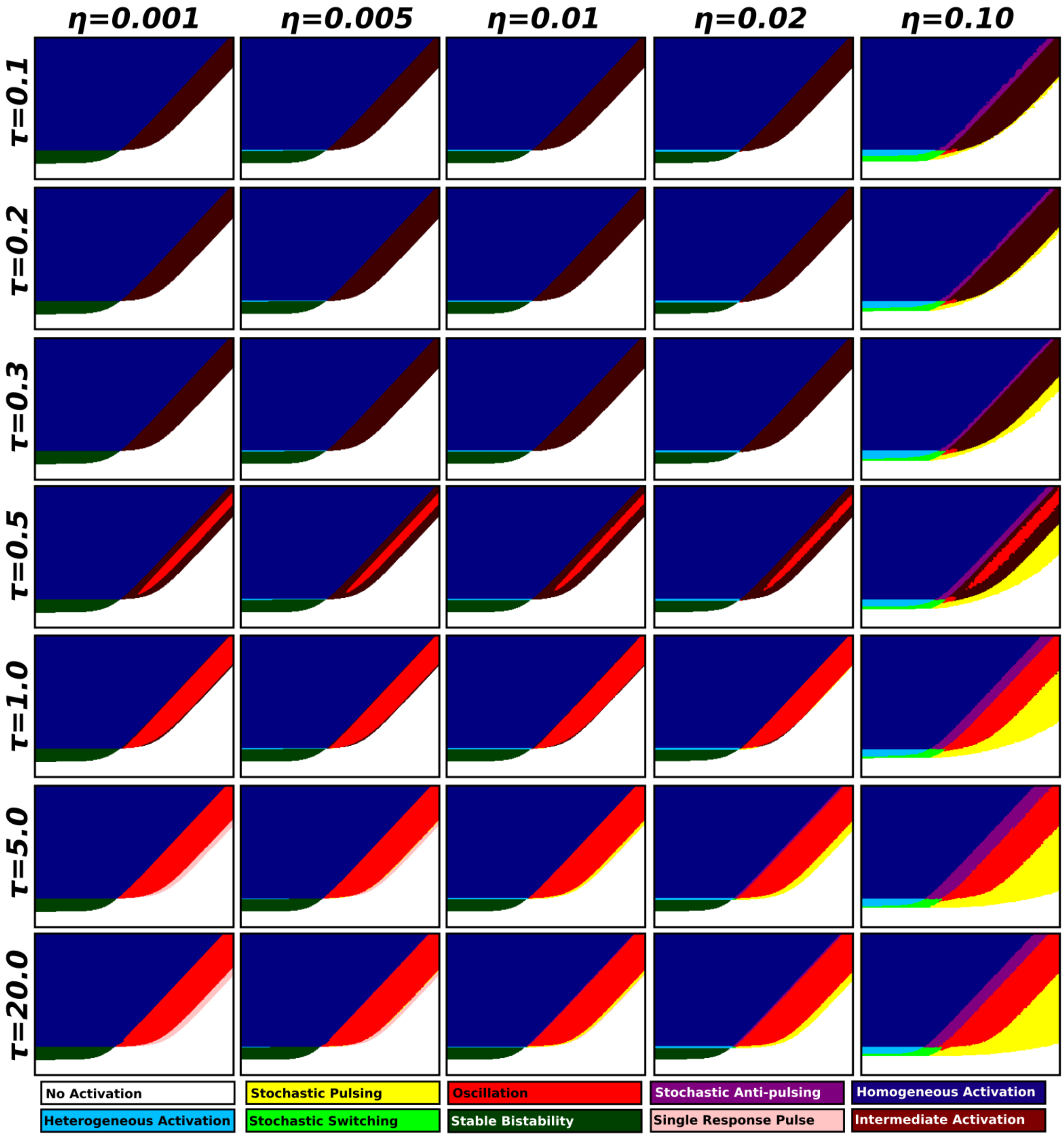

Supplementary Figure 29: **A grid of behaviour maps, for a range of values of the parameters $\boldsymbol{\tau}$ and $\boldsymbol{\eta}$.** The maps are plotted as described in Methods. For each map, we set $(v_0, n) = (0.1, 3)$, with $S \in (1.0, 150.0)$ (y-axis), and $D \in (0.1, 150.0)$ (x-axis) are used. The values of $\tau$ and $\eta$ are modified as indicated by the grid. The four upper maps in the leftmost column ($\eta = 0.001$, $\tau \in \{4/3, 20/9, 20/3, 100/3\}$) contain a slim region of the single response pulse behaviour (between the oscillatory and the no activation regions).

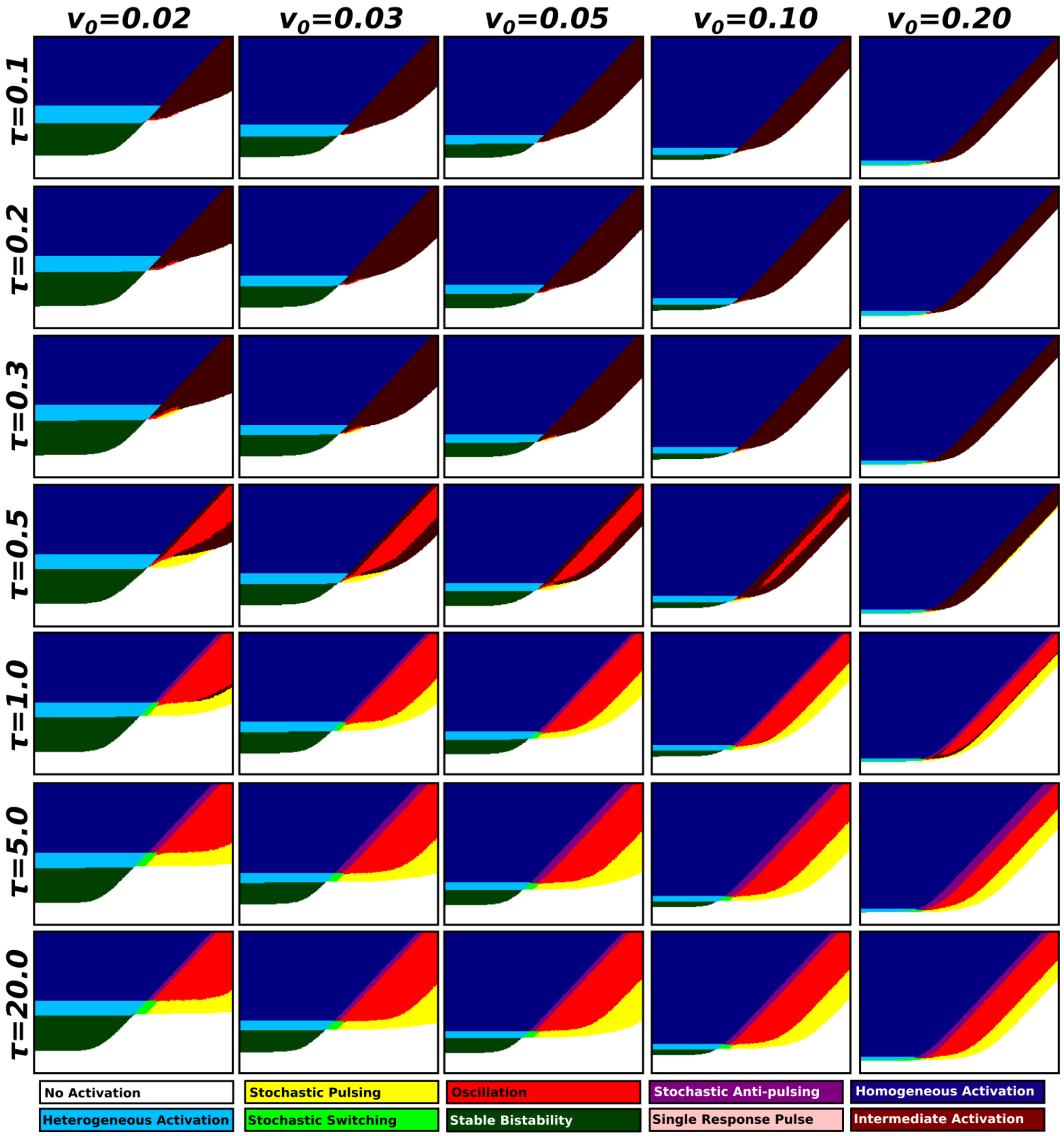

1026 Supplementary Figure 30: **A grid of behaviour maps, for a range of values of the parameters $\tau$ and**
1027 The maps are plotted as described in Methods. For each map, we set $(n, \eta) = (3, 0.05)$, with $S \in (1.0, 150.0)$
1028 (y-axis), and $D \in (0.1, 150.0)$ (x-axis) are used. The values of $\tau$ and $v_0$ are modified as indicated by the grid.

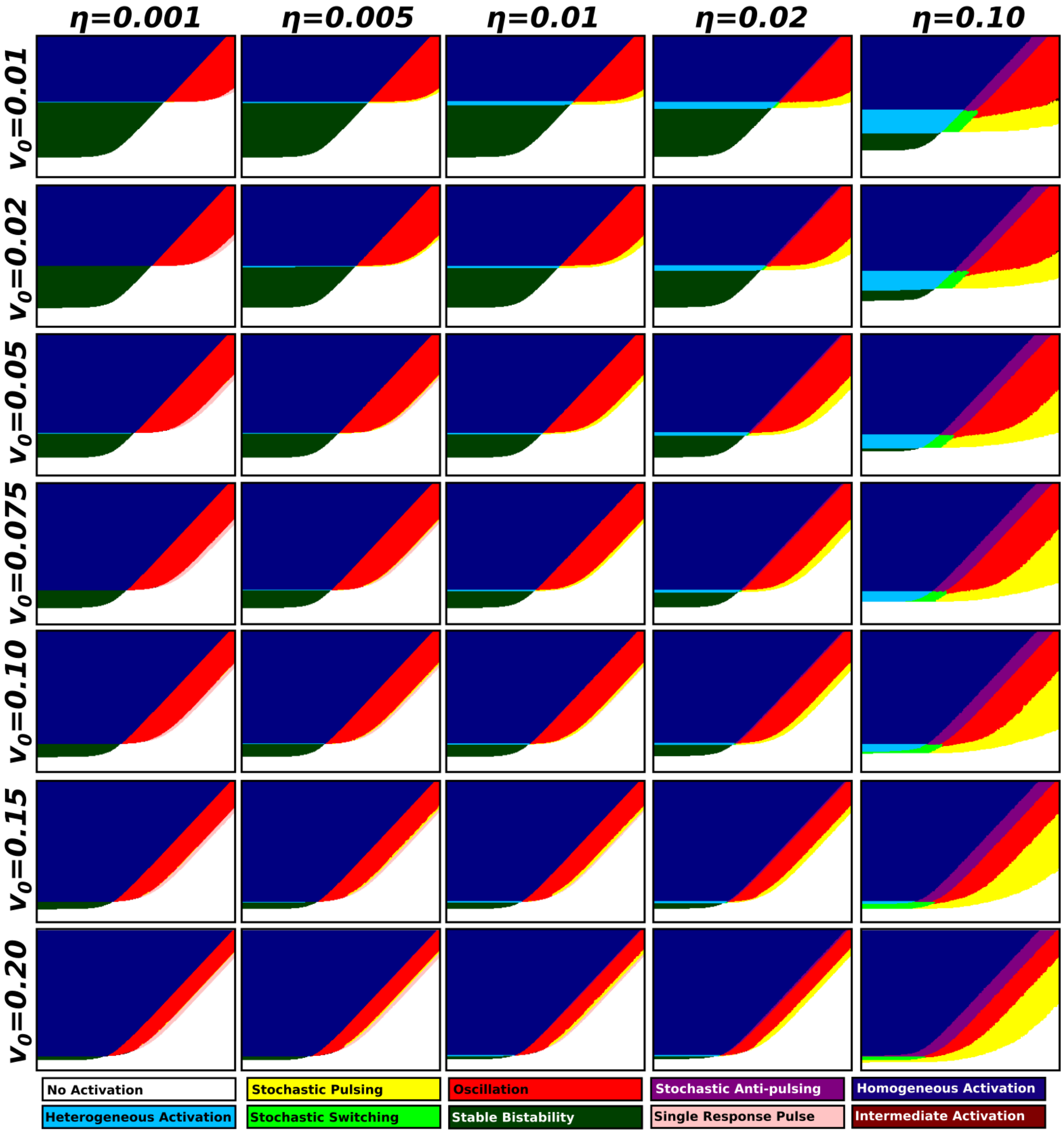

Supplementary Figure 31: **A grid of behaviour maps, for a range of values of the parameters $\eta$ and $v_0$.** The maps are plotted as described in Methods. For each map, we set $(\tau, n) = (10/3, 3)$, with $S \in (1.0, 150.0)$ (y-axis), and $D \in (0.1, 150.0)$ (x-axis) are used. The values of $\eta$ and $v_0$ are modified as indicated by the grid. The maps in the top two rows, and the two leftmost in the third highest row, contain a slim region of the single response pulse behaviour.

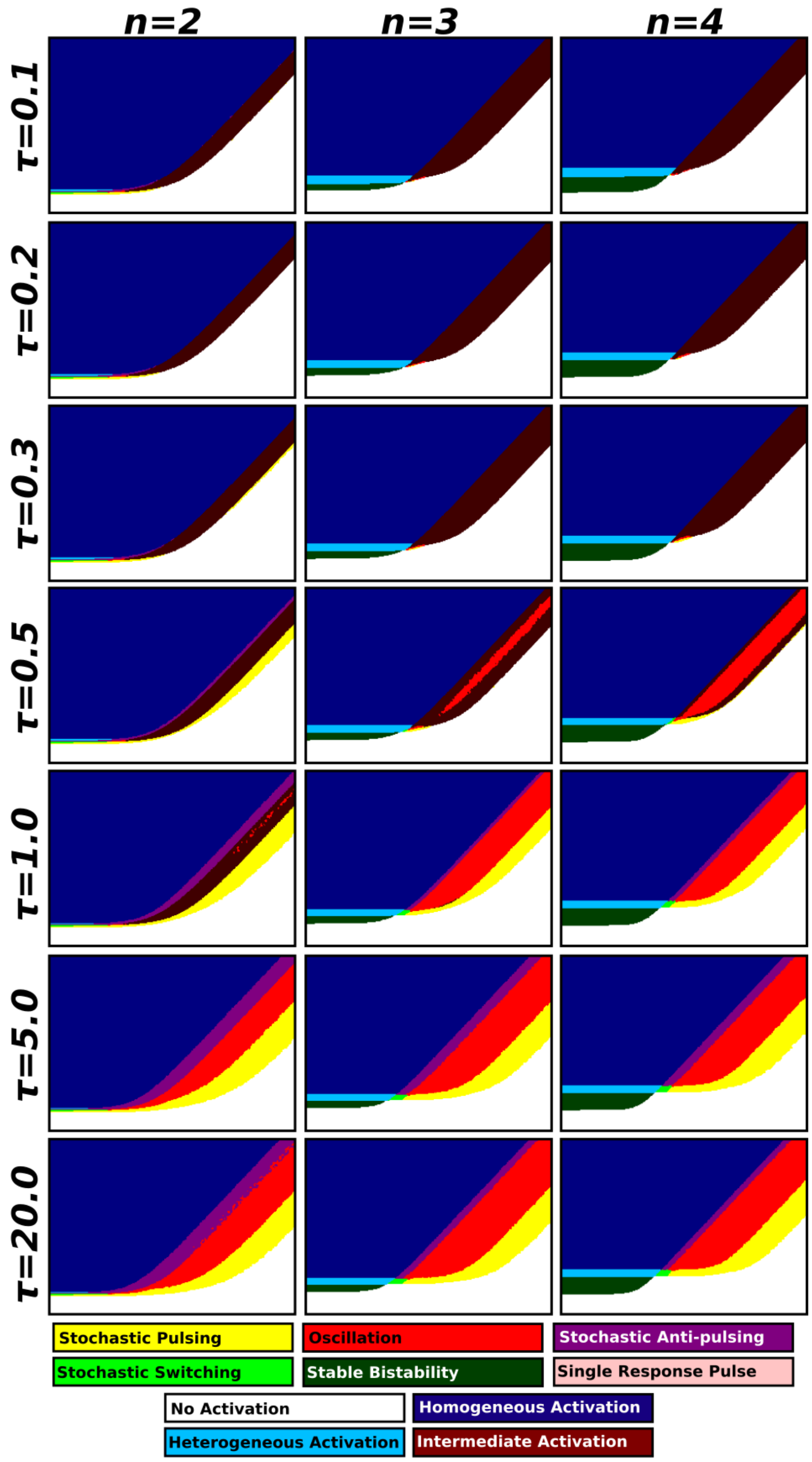

Supplementary Figure 32: **A grid of behaviour maps, for a range of values of the parameters $\boldsymbol{\tau}$ and $\boldsymbol{\eta}$.** The maps are plotted as described in Methods. For each map, we set $(, v_0, \eta) = (0.1, 0.05)$, with $S \in (1.0, 150.0)$ (y-axis), and $D \in (0.1, 150.0)$ (x-axis) are used. The values of $\tau$ and $n$ are modified as indicated by the grid.

### 9 Behaviour transition analysis

For parts of our analysis, we depend on the relative occurrences of the various transitions between different behaviours, and how these depend on the model's parameters. This analysis is presented here.

#### 9.1 Counting the transitions between different behaviour combinations

First, we investigated which (non-$S/D$) parameter(s) had the largest effect on the number of system transitions (i .e. transitions from one behaviour to another one as the critical parameter is changed). Here, $\tau$ had the largest impact on the behaviour of the system displayed. Next, $v_0$ and $\eta$ had similar impact, with $n$ having the smallest impact (Supplementary Table 4). Next, we scanned our 6-dimensional behaviour map, and counted all transitions from one behaviour to another one as the parameter $S$ was varied. A list of such transitions and their counts is presented in Supplementary Table 5. In Supplementary Table 6, we instead counts the occurrence of each transition (including all behaviours as $S$ is varied from small to large). These tables described which transitions are common (and rare) throughout behavioural space.

| **Parameter** | **Normalised transition count** |
|---|---|
| $\tau$ | 2.4980 |
| $v_0$ | 1.0518 |
| $n$ | 0.3944 |
| $\eta$ | 0.9203 |

Supplementary Table 4: **The relative frequency of behavioural transitions per parameter.** We wanted to determine which parameter (after $S$ and $D$ had the highest impact on displayed behaviour. To do this, for each parameter, we computed the average number of (non-trivial) transitions as that parameter was varied across its full range of values. Here, a non-trivial transition is one not including the no activation or homogeneous activation behaviours. Here, $\tau$ had the largest impact on the displayed behaviour, followed by $v_0$ and $\eta$. $n$ had the smallest effect on which behaviours were displayed.

#### 9.2 Noise warps parameter space

It is known that noise has a tendency to "warp" systems beyond just deviations from a mean behaviour. E.g. in [Loman and Locke, 2023] it was noted that the response pulse amplitude (of a single response pulse type system) increased with the amount of noise (rather than creating pulse amplitude variability). Generally, this is a known phenomenon for non-linear systems. It has also been noted that noise can induce oscillations in an otherwise non-oscillating system [Loman, 2025]. However, the phenomenon of noise causing non-intuitive changes to system behaviours has not been studied more systematically. By creating behaviour maps of stochastic behaviours, and comparing these to the corresponding deterministic cases, we have the opportunity to carry out this investigation. Here, we can note that noise does systematically "warps" the behavioural regions, as compared to what would be expected from the deterministic cases.

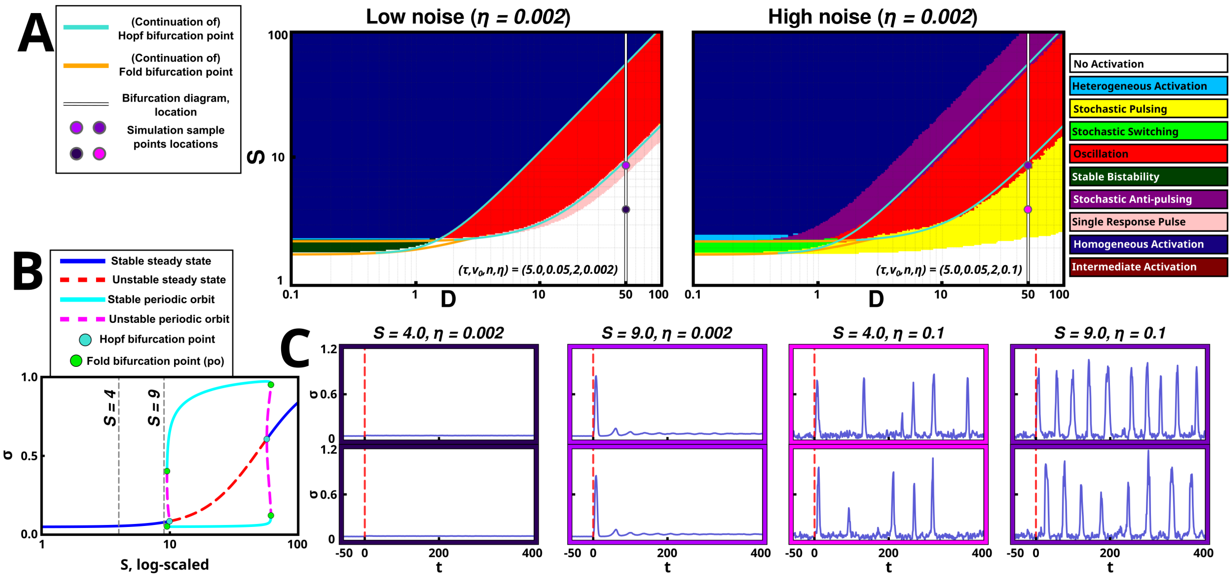

Supplementary Figure 33: **Stochasticity warps parameter space with respect to behavioural transitions.** (A) The same behaviour map, shown for a low and a high noise regime. In the map, we have plotted the bifurcation points of the deterministic system (studied in more detail in Supplementary Figures 11 and 27) across 2d $(S, D)$-space. Generally, the fold bifurcations are associated with bistability and the Hopf bifurcations with oscillations. We note that, for the low-noise regime, the behavioural transitions align well with the bifurcation points of the deterministic system. Meanwhile, as noise is increased, the behaviour regions shift. (B) Simulations for $S = 4$ and $S = 9$ (marked by grey dashed lines in the bifurcation diagram and dots in the behaviour maps in A), for the low and high noise regimes, respectively. We note that noise transitions the no activation behaviour (at $S = 4$) to a stochastic pulsing behaviour. Next, it transitions the single response pulse behaviour (at $S = 9$) to an oscillation behaviour (even if this parameter does not exhibit a periodic orbit for the deterministic case).

| Initial Behaviour | New Behaviour | Transition points |
|---|---|---|
| No activation | Stable bistability | 351372 |
| Heterogeneous activation | Homogeneous activation | 321327 |
| Stable bistability | Heterogeneous activation | 272292 |
| No activation | Stochastic pulsing | 242073 |
| Stochastic pulsing | Oscillation | 242051 |
| Stochastic anti-pulsing | Homogeneous activation | 215812 |
| Oscillation | Stochastic anti-pulsing | 200767 |
| Intermediate activation | Homogeneous activation | 135444 |
| No activation | Intermediate activation | 124724 |
| No activation | Single response pulse | 103933 |
| Oscillation | Homogeneous activation | 95710 |
| No activation | Homogeneous activation | 81993 |
| Stable bistability | Homogeneous activation | 79009 |
| Single response pulse | Oscillation | 56565 |
| Single response pulse | Stochastic pulsing | 43174 |
| Stochastic switching | Heterogeneous activation | 29945 |
| No activation | Stochastic switching | 28820 |
| Stochastic pulsing | Intermediate activation | 28133 |
| No activation | Oscillation | 24667 |
| Intermediate activation | Stochastic anti-pulsing | 20040 |
| Stochastic pulsing | Stochastic switching | 18355 |
| Stochastic switching | Stochastic anti-pulsing | 15824 |
| No activation | Heterogeneous activation | 9248 |
| Stochastic switching | Oscillation | 5226 |
| Single response pulse | Intermediate activation | 4615 |
| Stochastic anti-pulsing | Heterogeneous activation | 2161 |
| Intermediate activation | Heterogeneous activation | 2019 |

Supplementary Table 5: **List of behavioural transition points.** For each behavioural transition, the behaviour before and after the transition (as $S$ increases) is noted. Next, "Transition points" denote the number of points in our 6-dimensional parameter space where the transition occurs as $S$ is increased one step (showing how common the transition is). The list is sorted from the most common transitions to the less common ones.

| Transition | Count |
|---|---|
| no act. → stable bistability → het. act. → hom. act. | 272202 |
| no act. → stoch. pulsing → oscillation → stoch. anti-pulsing → hom. act. | 129105 |
| no act. → int. act. → hom. act. | 84198 |
| no act. → hom. act. | 81557 |
| no act. → stable bistability → hom. act. | 79009 |
| no act. → single response pulse → oscillation → hom. act. | 34748 |
| no act. → int. act. → oscillation → hom. act. | 31226 |
| no act. → stoch. switching → het. act. → hom. act. | 25560 |
| no act. → stoch. pulsing → oscillation → hom. act. | 19613 |
| no act. → single response pulse → stoch. pulsing → oscillation → stoch. anti-pulsing → hom. act. | 18638 |
| no act. → single response pulse → stoch. pulsing → oscillation → hom. act. | 17344 |
| no act. → oscillation → hom. act. | 13413 |
| no act. → stoch. pulsing → int. act. → oscillation → stoch. anti-pulsing → hom. act. | 10737 |
| no act. → stoch. pulsing → stoch. switching → stoch. anti-pulsing → hom. act. | 10208 |
| no act. → single response pulse → oscillation → stoch. anti-pulsing → hom. act. | 9612 |
| no act. → het. act. → hom. act. | 9147 |
| no act. → stoch. pulsing → int. act. → stoch. anti-pulsing → hom. act. | 8616 |
| no act. → oscillation → int. act. → hom. act. | 7270 |
| no act. → single response pulse → oscillation → int. act. → hom. act. | 8616 |
| no act. → stoch. pulsing → oscillation → int. act. → hom. act. | 7270 |

Supplementary Table 6: **List of full behavioural transitions.** For each possible transition where $S$ goes from small to large in our behaviour map, we check the behaviours it transitions through (taking the first occurrence of each behaviour). Here, we list the 20 most common such transitions and how frequently they occur. With one exception (intermediate activation to oscillation to intermediate activation), transition from, and then back into, a behaviour should not occur. To reduce noise, we have filtered out such cases. This includes some cases of transitions that include int. act. → oscillation → int. act., which is a fairly common (valid) transition (and the one exception from this list).

### 10 Additional experimental data

#### 10.1 Additional $\sigma^V$ data

Additional replicate measurements of the $\sigma^V$ system, not displayed in the main manuscript, are displayed in Supplementary Figures 34, 35.

#### 10.2 Additional $\sigma^B$ data

Additional replicate measurements of the $\sigma^V$ system, not displayed in the main manuscript, are displayed in Supplementary Figure 36-39.

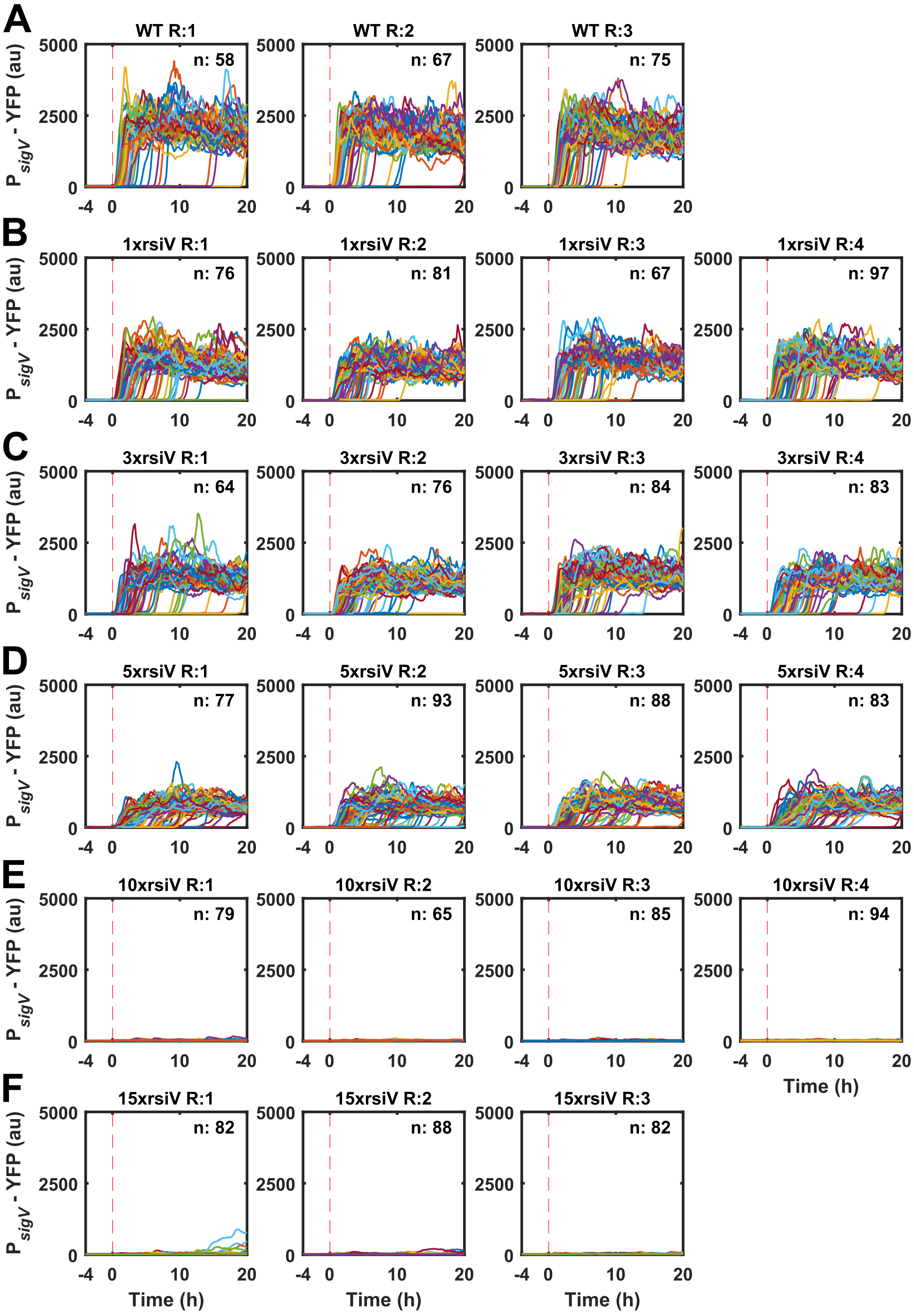

Supplementary Figure 34: **With increasing copy number of *rsiV* single cell $P_{sigV}$-YFP is decreasing under 1 µg/ml Lysozyme stress.** All shown repeats were grown overnight in the mother machine with no stress before exposing them to 1 µg/ml Lysozyme at time 0h. Each row is one *rsiV* mutant and each column is a repeat. A) WT (JLB130), B) 1x *rsiV* (JLB212), C) 3x *rsiV* (JLB288), D) 5x *rsiV* (JLB293), E) 10x rsiV (JLB325), F) 15x *rsiV* (JLB324). The trajectory colour represents the $P_{sigV}$-YFP time trajectory of one cell. n indicates the number of cells in each subplot.

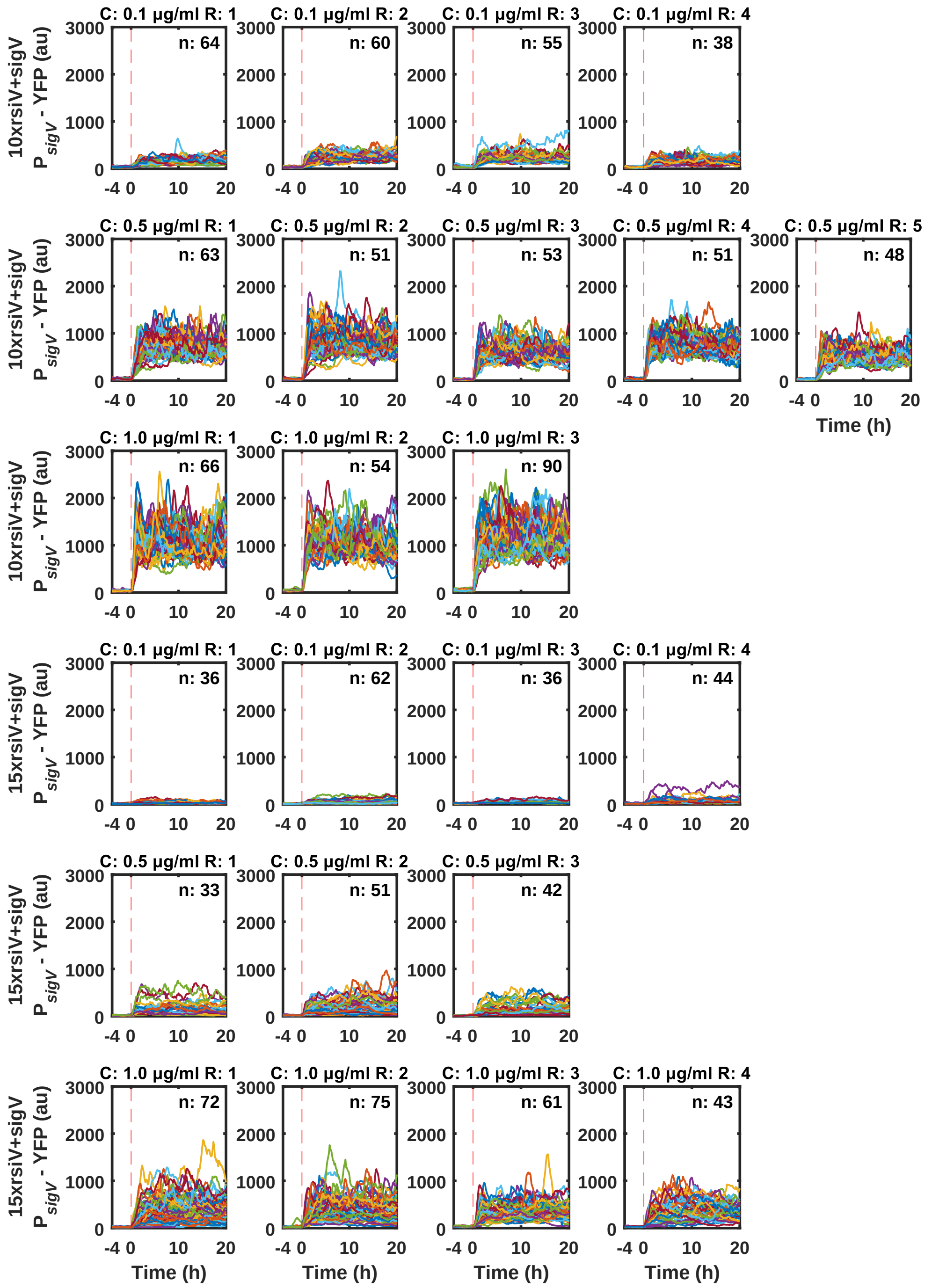

Supplementary Figure 35: **Adding an extra copy of *sigV* to 10x *rsiV* or 15x *rsiV* moves the behaviour into the intermediate regime.** All shown repeats were grown overnight in the mother machine with no stress before exposing them to 0.1, 0.5 or 1.0 µg/ml Lysozyme at time 0h. Each row is one *rsiV* mutant with its respective lysozyme concentration and each column is a repeat. A) 10x *rsiV* and *sigV* (JLB327) + 0.1 µg/ml Lysozyme, B) 10x rsiV and *sigV* (JLB327) + 0.5 µg/ml Lysozyme, C) 10x rsiV and *sigV* (JLB327) + 1.0 µg/ml Lysozyme, D) 15x rsiV and *sigV* (JLB345) + 0.1 µg/ml Lysozyme, E) 15x rsiV and *sigV* (JLB345) + 0.5 µg/ml Lysozyme, F) 15x *rsiV* and *sigV* (JLB345) + 1.0 µg/ml Lysozyme. The trajectory colour represents the $P_{sigV}$-YFP time trajectory of one cell. n indicates the number of cells in each subplot.

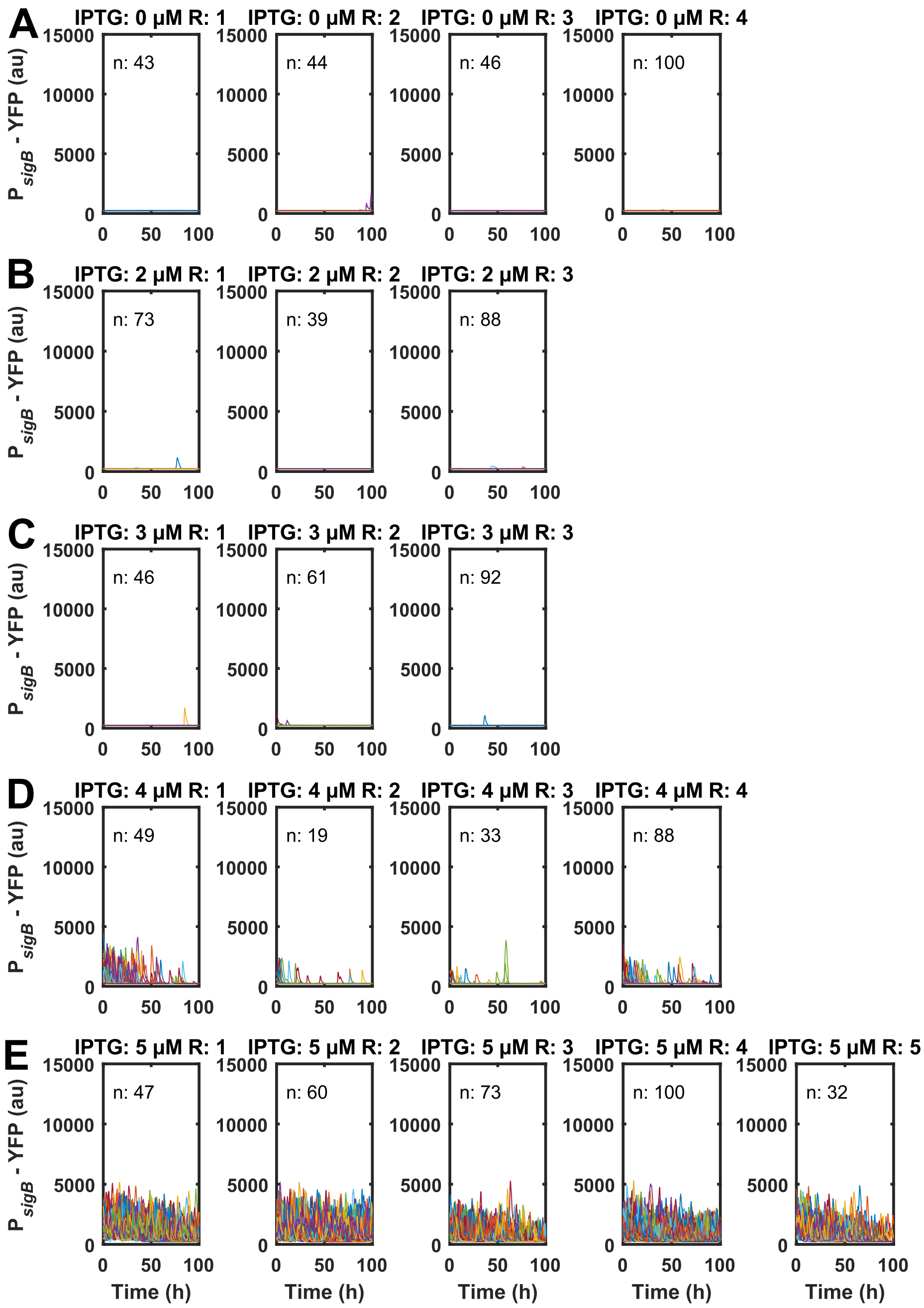

Supplementary Figure 36: **In the Phs-rsbPΔ4 (ΔPAS-CC) ΔrsbQ ΔrsbU mutant we can tune the $P_{sigB}$-YFP output behaviour based on the IPTG levels.** The Phs - rsbPΔ4 (ΔPAS-CC) ΔrsbQ ΔrsbU mutant (JLB346) was grown with the following IPTG levels: A) 0 µM, B) 2 µM, C) 3 µM, D) 4 µM, E) 5 µM. The trajectory colour represents the $P_{sigB}$-YFP time trajectory of one cell. n indicates the number of cells in each subplot.

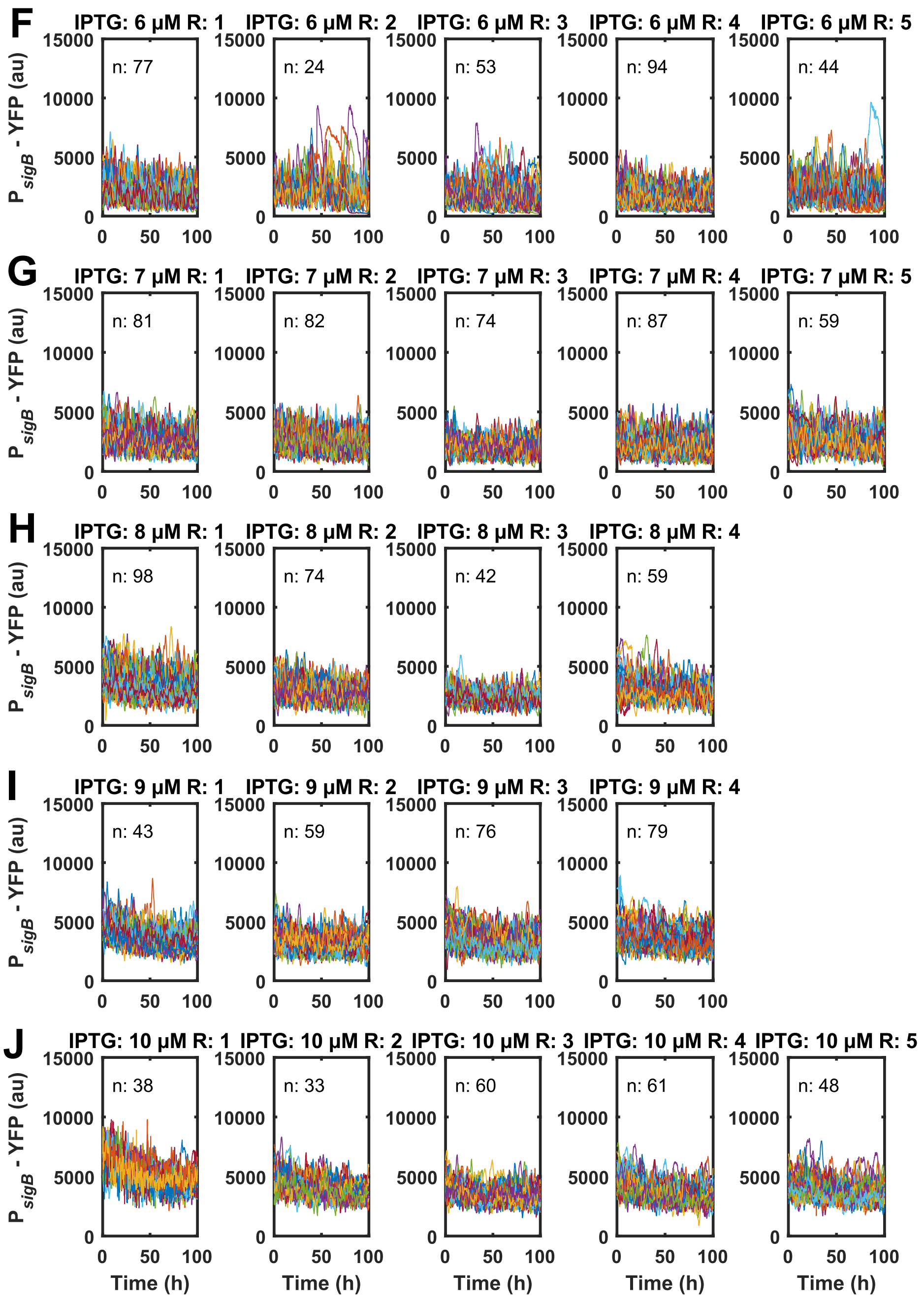

Supplementary Figure 37: **In the Phs - rsbPΔ4 (ΔPAS-CC) ΔrsbQ ΔrsbU mutant we can tune the $P_{sigB}$-YFP output behaviour based on the IPTG levels.** The Phs - rsbPΔ4 (ΔPAS-CC) ΔrsbQ ΔrsbU mutant (JLB346) was grown with the following IPTG levels: F) 6 µM, G) 7 µM, H) 8 µM, I) 9 µM, J) 10 µM. The trajectory colour represents the $P_{sigB}$-YFP time trajectory of one cell. n indicates the number of cells in each subplot.

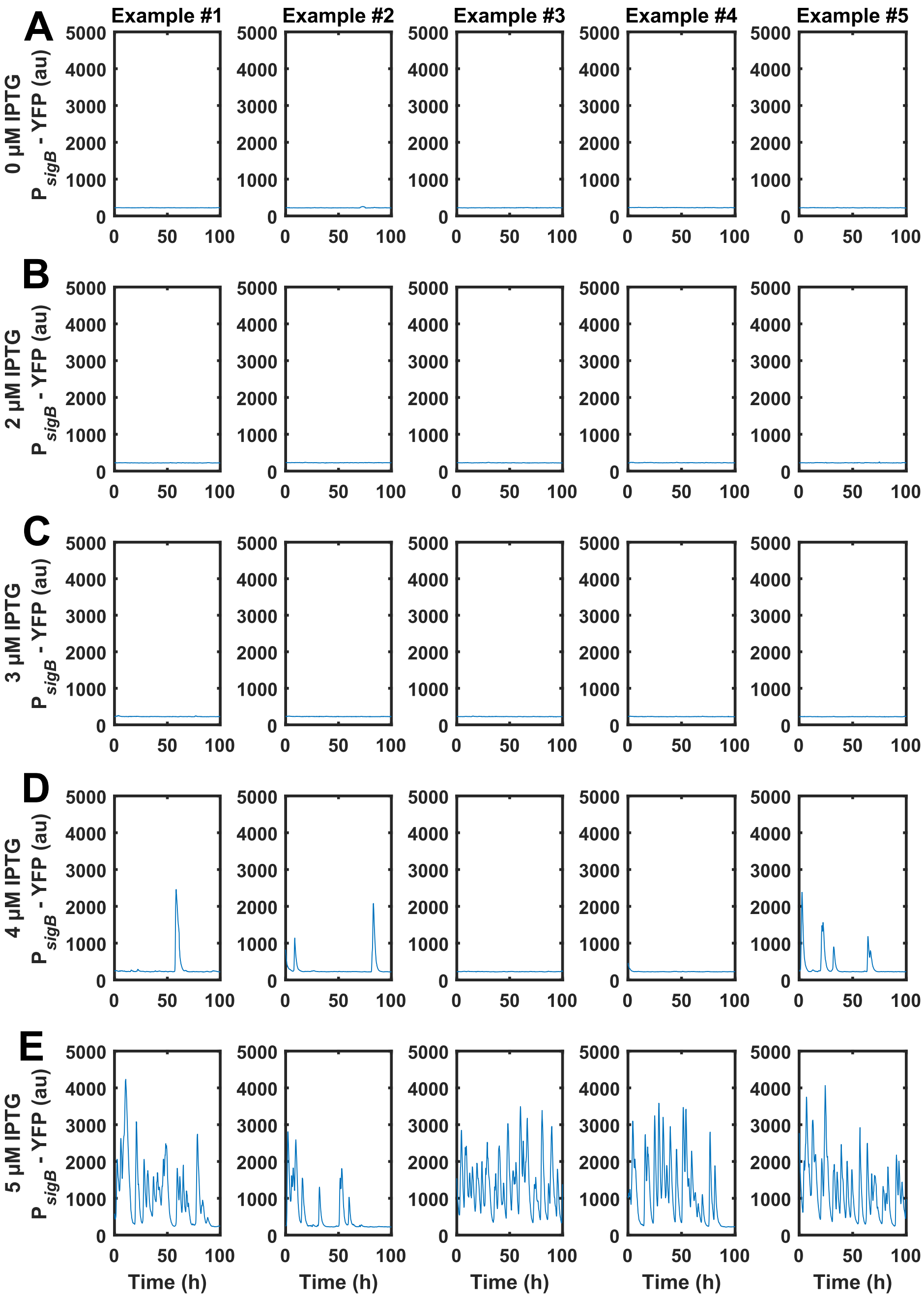

Supplementary Figure 38: **In the Phs - rsbPΔ4 (ΔPAS-CC) ΔrsbQ ΔrsbU mutant we can tune the $P_{sigB}$-YFP output behaviour based on the IPTG levels on a single cell level.** The Phs - rsbPΔ4 (ΔPAS-CC) ΔrsbQ ΔrsbU mutant (JLB346) was grown with the following IPTG levels: A) 0 µM, B) 2 µM, C) 3 µM, D) 4 µM, E) 5 µM. In each subfigure the $P_{sigB}$-YFP time trajectory of one cell is plotted (blue line).

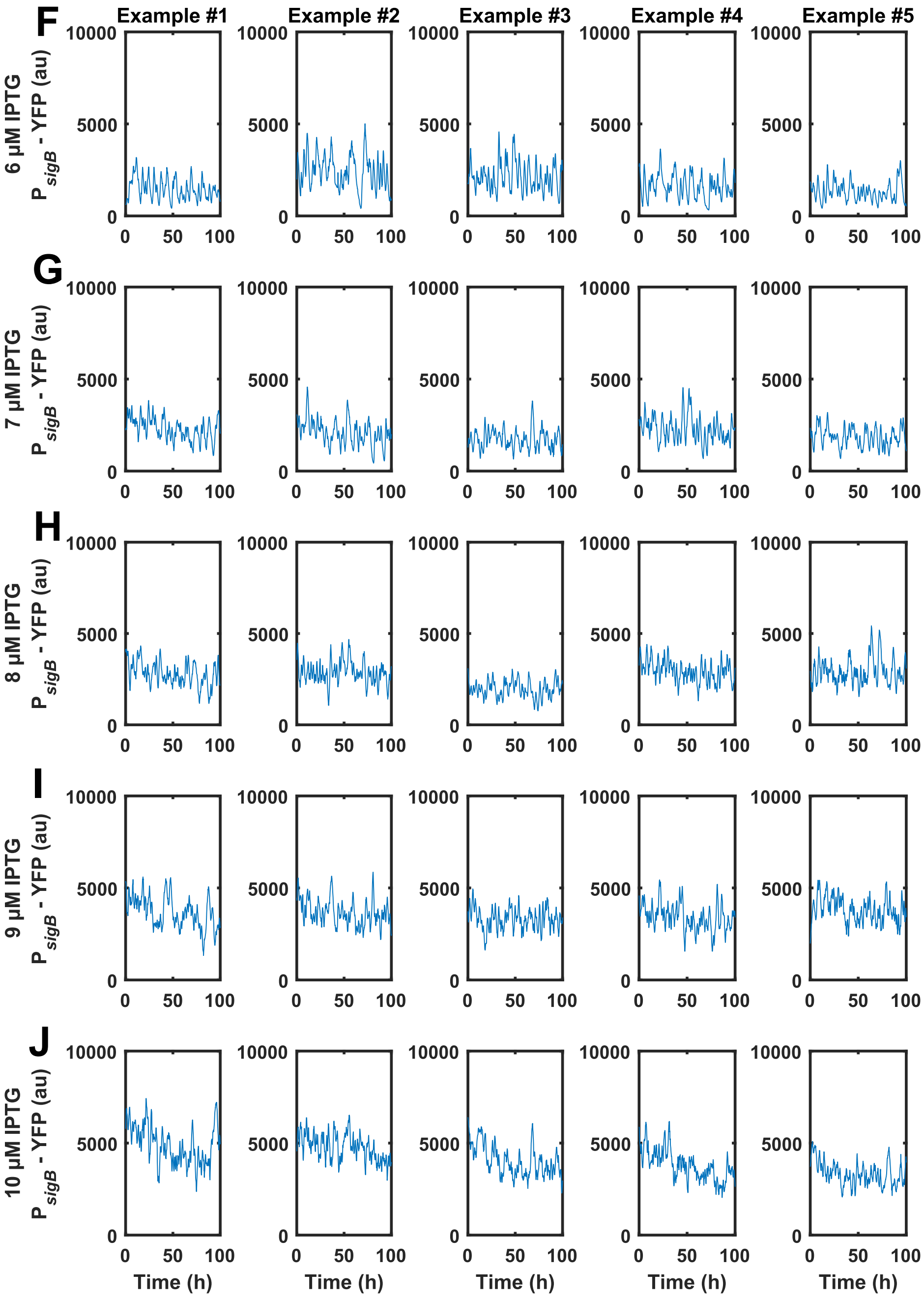

Supplementary Figure 39: **In the Phs - rsbP$\Delta$4 ($\Delta$PAS-CC) $\Delta$rsbQ $\Delta$rsbU mutant we can tune the P$_{sigB}$-YFP output behaviour based on the IPTG levels on a single cell level.** The Phs - rsbP$\Delta$4 ($\Delta$PAS-CC) $\Delta$rsbQ $\Delta$rsbU mutant (JLB346) was grown with the following IPTG levels: F) 6 µM, G) 7 µM, H) 8 µM, I) 9 µM, J) 10 µM. In each subfigure the P$_{sigB}$-YFP time trajectory of one cell is plotted (blue line).

### 11 Combined experiments/simulations classifier

Our original classifier depends on computing system steady state properties (using a combination of additional simulations and analysis of the model's analytic expression). Because of this, it cannot be applied to experimental trajectories. To validate whether the transition seen, when $S$ is varied in the $\sigma^{\mathrm{B}}$ system, replicates the pulsing/oscillatory transition predicted by the model, we design a new classifier. This was inspired by our previous ones (by attempting to compute the distribution of transition times), but could classify time-trajectories directly (while being agnostic to whether these are generated by simulations or experimental measurements).

For a given time trajectory, $t_{ii=1}^{n}$, the classifier assigns one of the following behaviours:

- **No activation:** To trajectories which are inactive
- **Stochastic pulsing:** To trajectories which exhibit activity pulses from an otherwise inactive state
- **Oscillation:** To trajectories that oscillate
- **Homogeneous activation:** To trajectories which are active
- **Unclassified:** This behaviour is assigned to trajectories which does not fall into one of the above classes. It is primarily used as a validation of our classifier's validity (i.e. a large number of "unclassified" classifications would indicate a problem with our classifier).

The model assigns behaviours according to the following heuristic:

1. We noted that the distribution of all values encountered across all trajectories was bimodal. We used this to estimate the inactivation and activation thresholds.
2. The *inactivation threshold* was set as $i_{thres} = 4p_l$ (where $p_l$ is the value of the lower peak). The *activation threshold* was set as $i_{thres} = p_h/2$ (where $p_l$ is the value of the higher peak).
3. For each individual trajectory we computed the *activation type* as follows (and the *deactivation type* in the reverse manner):
   (a) Trajectories which never surpassed the activation threshold were classified as *NA*.
   (b) We counted all times between the system entering an inactive state until it surpassed the activation threshold. Each of these times was classed as either *heterogeneous* or *instantaneous* (depending on whether they were longer than a fixed threshold).
   (c) If the number of *heterogeneous* activation times were more than the number of *instantaneous* activation times, the activation type was classified as *heterogeneous* (else as *instantaneous*).
4. Trajectories with activation type NA were classified as **no activation**.
5. Trajectories with activation type heterogeneous and deactivation type instantaneous or NA were classified as **stochastic pulsing**.
6. Trajectories with activation and deactivation type instantaneous were classified as **oscillation** of their Welch-type periodogram peak surpassed a specific threshold. Else, if their median value surpassed the activation threshold, they were classified as **homogeneous activation**.
7. Trajectories with deactivation type NA were classified as **homogeneous activation**.
8. Trajectories with deactivation type heterogeneous, and activation type instantaneous or NA, were classified as **homogeneous activation**.
9. Trajectories not assigned to any classes according to this scheme were classified as **unclassified**.

To further demonstrate the soundness of this classification scheme, we plotted randomly selected trajectories (simulated and experimentally measured) and their corresponding classification (Supplementary Figures 40, 41). By considering these plots, we can confirm that behaviours are correctly classified.

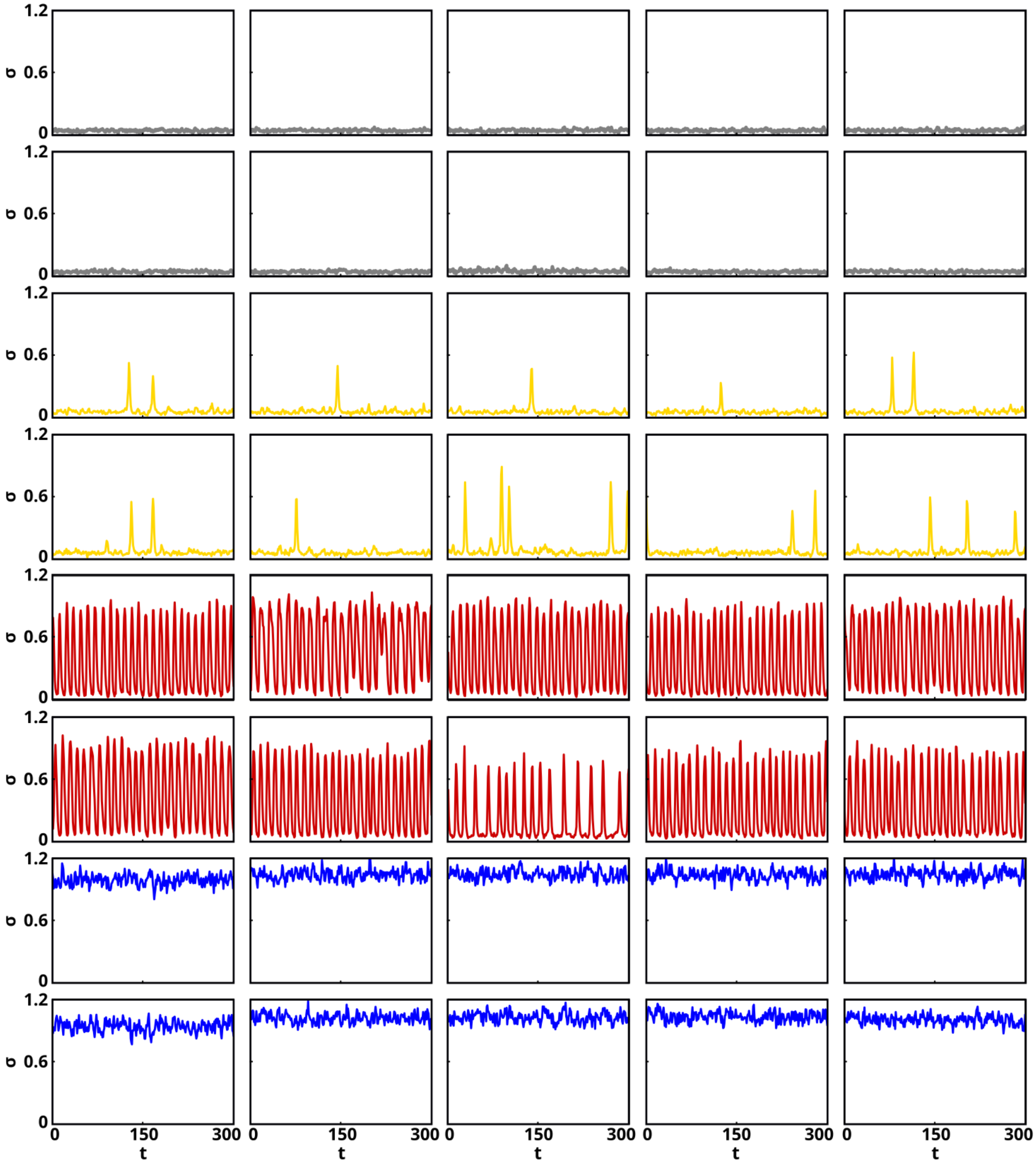

Supplementary Figure 40: **The combined classifier successfully assigns the correct class to simulated trajectories.** The combined classifier classifies a trajectory as either ”no activation” (grey), ”stochastic pulsing” (yellow), ”oscillation” (red), or ”homogeneous activation” (blue). For each behaviour, we have randomly selected 10 simulated trajectories that were assigned to that behaviour and plotted them (here selected from the behaviour map used in Figure 5). Considering the trajectories, they have all been correctly classified. A final class, ”unclassified”, was assigned to behaviours that couldn't be assigned to any other behaviour. Out of 3,020,000 simulated trajectories that were classified, none were assigned as unclassified, further suggesting the validity of the classifier.

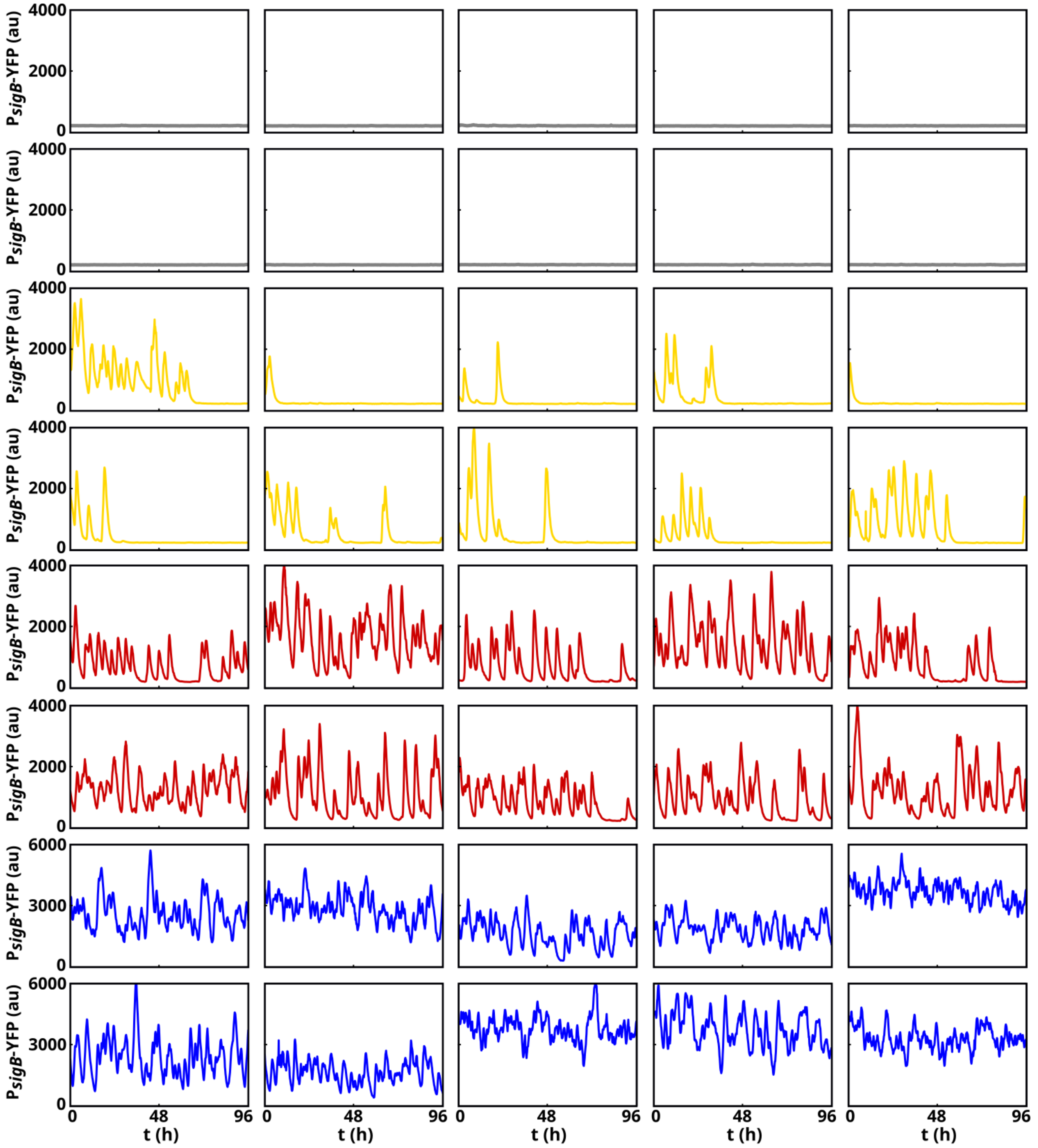

Supplementary Figure 41: **The combined classifier successfully assigns the correct class to experimental trajectories.** The combined classifier classifies a trajectory as either "no activation" (grey), "stochastic pulsing" (yellow), "oscillation" (red), or "homogeneous activation" (blue). For each behaviour, we have randomly selected 10 experimental trajectories that were assigned to that behaviour and plotted them. Considering the trajectories, they have all been correctly classified. A final class, "unclassified", was assigned to behaviours that couldn't be assigned to any other behaviour. Out of 2395 experimental trajectories that were classified, only 4 were assigned as unclassified, further suggesting the validity of the classifier.

### 12 Experimental set-up

#### 12.1 Wafer fabrication

The microfluidic master was fabricated using maskless photolithography by exploiting a Dilase 650 Kloe direct laser writer (Kloe, France). The microfluidic design is based on the popular mother machine design (29) but allows for up to eight simultaneous conditions to be investigated. The devices were designed using AutoCad (Autodesk, USA). Briefly, for the first layer (growth channels) SU8 6001 (Kayaku Advanced Material, Japan) was spun coated (LabSpin, Suss, MicroTec,Germany) onto a silicon substrate (Si-Mat, Germany) at 1500 rpm. For the second layer (feeding channels) SU8 2050 (Kayaku Advanced Material, Japan) was spun at 2000 rpm.

##### Wafer cleaning

- The silicon substrate was cleaned in an acetone bath (Merck, USA) for 5 min.
- Then the wafer was rinsed with IPA (Merck, USA), blow dried with Nitrogen (BOC, UK) and dehydrated on a hot plate (Stuart, UK) at 200°C for 5 min.

##### Focus test

We found that doing a focus test on each wafer in the different areas where the final design would go before the actual exposure improved our results greatly. This was done before any exposure of the growth channels. Ramping the temperatures was critical to improve adhesion.

- SU8 6001 (Kayaku Advanced Material, Japan) was spun coated (LabSpin, Suss MicroTec, Germany) at 1500 rpm.
- Softbaking: 30 s 65°C → 4 min 110°C → 30 s 65°C.
- The exposure was done with a Dilase 650 direct laser writer (Kloe, France) using a beam size of 1 um. Speed: 100 mm/s and 100% power.
- Post exposure bake: 30 s 65°C → 30 s 110°C → 30 s 65°C.
- The photoresist was then developed in PGMEA (Merck, USA) for 3 min.
- For each area of a final design the focus was noted. The results greatly depended on the focus and small changes in focus could make a big difference.
- Finally, the crosslinked SU8 was stripped off by wiping the wafer with acetone and cleaned again as described in the section on wafer cleaning.

##### Growth Channels

The parameters for the growth channel were the same as for the focus test except for a hard bake. ThesePost exposure bake: 30 s 65°C → 30 s 110°C → 30 s 65°C. parameters result in a channel height of 1 um.

- SU8 6001 (Kayaku Advanced Material, Japan) was spun coated (LabSpin, Suss MicroTec, Germany) at 1500 rpm.
- Softbaking: 30 s 65°C → 4 min 110°C → 30 s 65°C.
- The exposure was done with a Dilase 650 direct laser writer (Kloe, France) using a 1 µm beam size. Speed: 100 mm/s and 100% power.
- Post exposure bake: 30 s 65°C → 30 s 110°C → 30 s 65°C.
- Development in PGMEA for 3 min.
- Hard bake: 30 s 65°C → 30 s 110°C → 5 min 200°C → 30 s 110°C → 30 s 65°C.

**Feeding Channels**

The outlined parameters below resulted in a channel height of 80 um.

- SU8 2050 (Kayaku Advanced Material, Japan) was spun coated (Polos Spin 150i, Netherlands) at 2000 rpm.
- Softbaking: 3 min 65°C → 9 min 95°C.
- The exposure was done with a Dilase 650 direct laser writer (Kloe, France) using a 10 µm beam. Speed: 100 mm/s and 25% power.
- Post exposure bake: 2 min 65°C → 7 min 95°C.
- Development in PGMEA for 7 min.
